# Dynamic structural adaptations enable the endobiotic predation of *bdellovibrio bacteriovorus*

**DOI:** 10.1101/2022.06.13.496000

**Authors:** Mohammed Kaplan, Yi-Wei Chang, Catherine M. Oikonomou, William J. Nicolas, Andrew I. Jewett, Stefan Kreida, Przemysław Dutka, Lee A. Rettberg, Stefano Maggi, Grant J. Jensen

## Abstract

*Bdellovibrio bacteriovorus* is an endobiotic microbial predator that offers promise as a living antibiotic for its ability to kill Gram-negative bacteria, including human pathogens. Even after six decades of study, fundamental details of its predation cycle remain mysterious. Here, we used cryo-electron tomography to comprehensively image the lifecycle of *B. bacteriovorus* at nanometer-scale resolution. In addition to providing the first high-resolution images of predation in a native (hydrated, unstained) state, we also discover several surprising features of the process, including novel macromolecular complexes involved in prey attachment/invasion and a flexible portal structure lining a hole in the prey peptidoglycan that tightly seals the prey outer membrane around the predator during entry. Unexpectedly, we find that *B. bacteriovorus* does not shed its flagellum during invasion, but rather resorbs it into its periplasm for degradation. Finally, following replication and division in the bdelloplast, we observe a transient and extensive ribosomal lattice on the condensed *B. bacteriovorus* nucleoid.

**Graphical abstract:** 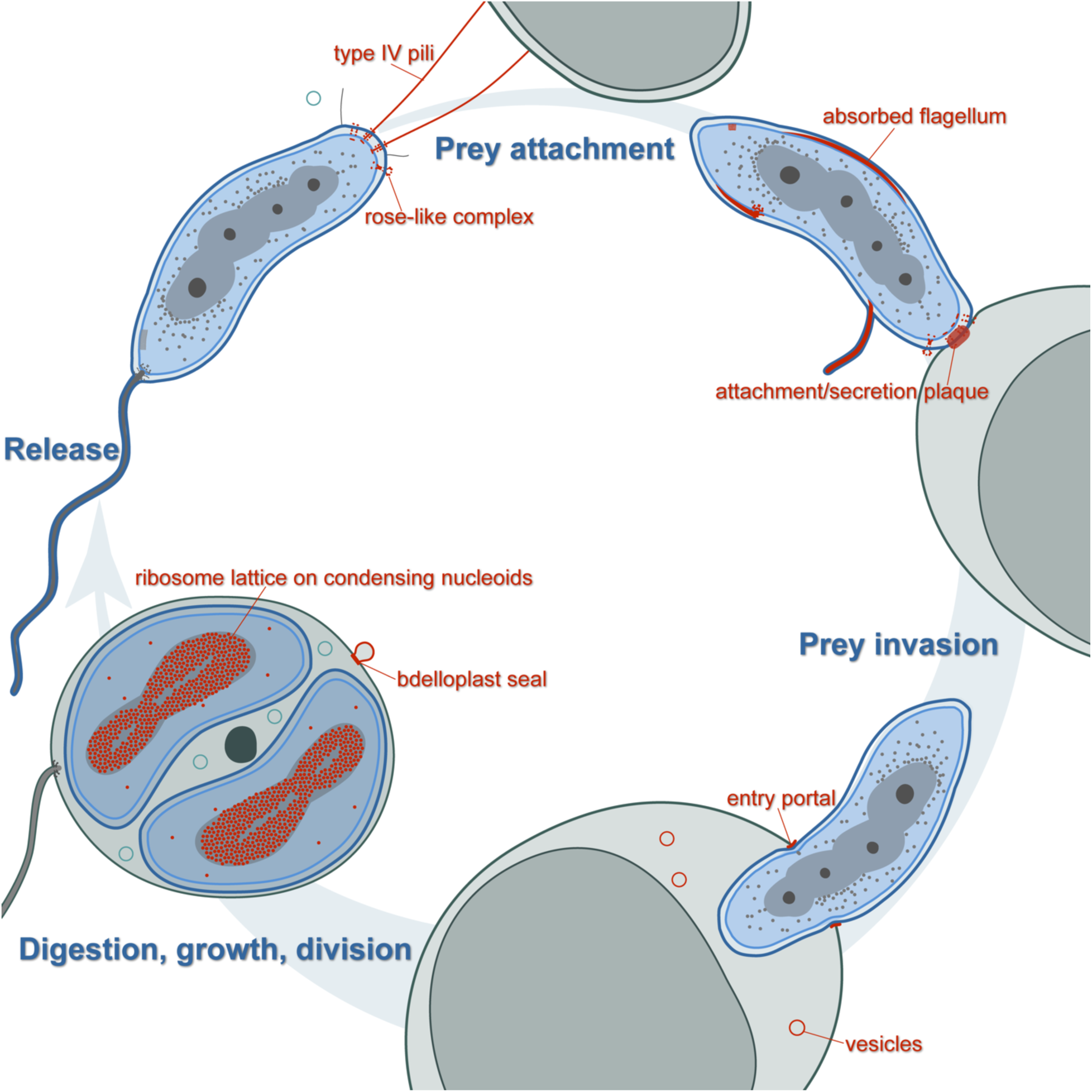

## Introduction

Predation is a widespread behavior, from the largest eukaryotes to the smallest viruses, that drives evolution and energy flow in biological communities. In bacteria, predatory behavior is common and multiple predation types have been described, including epibiotic strategies in which the predator remains outside the prey and endobiotic strategies in which the predator invades the prey’s cytoplasm or periplasm (Pérez et al., 2016). Since its description in 1963 as the first known bacterial parasite of bacteria (Stolp and Starr, 1963), *Bdellovibrio bacteriovorus* has been a paradigm of periplasmic endobiotic predation, in which a small predator takes up residence in a larger diderm prey’s periplasm (Laloux, 2020; Sockett, 2009). The ability of *Bdellovibrio*-and-like-organisms (BALOs) to invade other Gram-negative bacteria, including human pathogens, has rendered them potential candidates for living antibiotics to tackle the crisis of antimicrobial resistance that has emerged in the past few decades (Bratanis et al., 2020; Cavallo et al., 2021; Harini et al., 2013; Iebba et al., 2014; Madhusoodanan, 2019; Negus et al., 2017; Pantanella et al., 2018; Raghunathan et al., 2019; Russo et al., 2018; Shatzkes et al., 2017a, 2017b), or even as a possible probiotic agent (Bonfiglio et al., 2020). For example, it has been shown that treating zebrafish infected with *Shigella flexneri* with *B. bacteriovorus* increases the animals’ survival rate (Willis et al., 2016). In addition, microbial predation has been suggested to have been involved in pivotal evolutionary events including the genesis of eukaryotic cells, the rise of multicellularity, and pathogenicity (Davidov and Jurkevitch, 2009; Erken et al., 2013; Lyons and Kolter, 2015).

The predatory lifecycle of *B. bacteriovorus* has been extensively studied by methods including conventional transmission electron microscopy, light/fluorescence microscopy, helium-ion microscopy, atomic force microscopy and biochemical assays (Burnham et al., 1968; Kuru et al., 2017; Makowski et al., 2020; Núñez et al., 2003; Said et al., 2019; Stolp and Starr, 1965), and is the subject of several excellent reviews (see for example: (Cavallo et al., 2021; Laloux, 2020; Negus et al., 2017; Rotem et al., 2014; Sockett, 2009)). In the free-living attack phase, the predator is transcriptionally streamlined, with a highly compacted spiral nucleoid (Butan et al., 2011), and a vibrioid cell shape caused by the asymmetric activity of a peptidoglycan hydrolase (Banks et al., 2022). At one pole of the cell, a sheathed unipolar flagellum enables high-velocity (up to 160 µm/s (Lambert et al., 2006; Rendulic, 2004)) motile collisions with prey. The flagellar filament has a distinctive damped waveform due to segments made up of subunits with different helical properties (Thomashow and Rittenberg, 1985a). Initial interaction with, and attachment to, prey is mediated by type IVb (Avidan et al., 2017) and type IVa pili (T4P) (Evans et al., 2007; Mahmoud and Koval, 2010; Milner et al., 2014) at the pole opposite the flagellum (the “biting pole”). Following this initial interaction, *B. bacteriovorus* uses flagellum-independent gliding motility to reach an attachment site on the side of rod-shaped prey cells (Lambert et al., 2011).

Upon initial attachment, prey quality is assessed in a cyclic-di-GMP-dependent process (Caulton and Lovering, 2020; Hobley et al., 2012; Meek et al., 2019), and if the prey is found to be suitable and not already under attack by another predator (Lerner et al., 2012), the attachment is made permanent and the process of invasion begins. Initially, it was suggested that a “drilling” mechanism caused by flagellar rotation might play a role in the invasion process (Burnham et al., 1968; Stolp and Starr, 1965), but later studies found that flagellar motility-compromised mutants can still invade prey, which fits with the roles of pili and gliding motility mentioned above (Lambert et al., 2006). Another early penetration model hypothesized that entry is passive and occurs due to osmotic forces from the flux of solutes and water resulting from structural changes in the prey envelope (Abram et al., 1974).

Prey invasion involves enzymatic modification of the prey cell wall by *B. bacteriovorus* at the predator-prey contact point (Kuru et al., 2017; Thomashow and Rittenberg, 1978c, 1978b, 1978a; Tudor et al., 1990) and is associated with the secretion, by type I and II secretion systems, of many enzymes (Pasternak et al., 2014; Rendulic, 2004), including glycanases (Harding et al., 2020; Thomashow and Rittenberg, 1978a), peptidases (Lerner et al., 2012; Tudor et al., 1990) and deacetylases (Lambert et al., 2016). A self-protection protein in *B. bacteriovorus* inhibits the predator’s peptidases, thereby allowing specific modification of the prey cell wall (Lambert et al., 2015). These modifications round the prey cell and greatly expand the periplasmic space, providing room for the predator to enter. Entry itself is rapid, occurring in a few minutes or less (Abram et al., 1974; Capeness et al., 2013). With a few exceptions noted (Iida et al., 2009; Lambert et al., 2006), flagella are no longer visible on *B. bacteriovorus* when they enter their prey (Shilo, 1969; Thomashow, L. S., 1979).

Once *B. bacteriovorus* enters the prey’s periplasm, the entry pore is sealed and transpeptidases modify the prey cell wall to render it more robust to osmotic pressure (Kuru et al., 2017; Lerner et al., 2012), forming what is known as the bdelloplast (Starr and Baigent, 1966). If prey-derived cues indicate the availability of sufficient nutrients in the bdelloplast (Rotem et al., 2015), *B. bacteriovorus* enters its growth phase. The predator consumes the cytoplasmic contents of the prey, facilitated by translocation of outer membrane (OM) pore proteins into the prey’s cytoplasmic membrane (Tudor and Karp, 1994) and secretion of nucleases to digest the prey’s nucleic acids (Bukowska-Faniband et al., 2020). This fuels predator growth, through bidirectional elongation, and replication of its genetic material (Makowski et al., 2019). Once available nutrients are exhausted, the predator septates synchronously into multiple progeny cells by non-binary fission, with the number of progeny cells depending on the size of the prey (Fenton et al., 2010a; Kaljević et al., 2021). Finally, the progeny cells, reset to the attack phase, use an enzyme specific for deacetylated peptidoglycan to lyse the prey cell wall (Harding et al., 2020; Lambert et al., 2016), creating pores through which they exit the bdelloplast and move on in search of new prey (Fenton et al., 2010a). If the first invasion does not yield sufficient nutrients for replication and division, a *B. bacteriovorus* cell may leave the first bdelloplast and complete its lifecycle in a second prey (Makowski et al., 2019).

Despite extensive study, the structural details of much of the invasion process remain unclear. Most of the previous work relied on low-resolution imaging methods or conventional electron microscopy preparations, in which dehydration and fixation disrupt cell membranes and obscure macromolecular details. Cryogenic electron tomography (cryo-ET) allows the investigation of cellular processes in a fully-hydrated frozen state with macromolecular resolution (Ghosal et al., 2019a; Kaplan et al., 2021a; Oikonomou and Jensen, 2017), but so far it has only been applied to study the ultrastructure of individual *B. bacteriovorus* cells in the attack phase. While these studies revealed important structural features of this stage (Borgnia et al., 2008; Butan et al., 2011; Fenton et al., 2010b), much remains to be learned about the full invasion cycle of *B. bacteriovorus*.

Here, we used cryo-ET to image the predation cycle of *B. bacteriovorus* invading three types of prey: *Vibrio cholerae, Escherichia coli*, and *E. coli* minicells. Our work reveals, for the first time, the macromolecular details of each stage of the *B. bacteriovorus* lifecycle and uncovers several unexpected features of the process, including absorption of the extracellular flagellum into the predator’s periplasm during attachment, a flexible portal structure associated with the prey peptidoglycan surrounding the predator during entry, and formation of a ribosome lattice around the predator’s nucleoid after the prey is consumed in the bdelloplast.

## Results

To visualize the predatory lifecycle of *B. bacteriovorus* in a near-native state, we applied cryo-ET imaging to samples of *B. bacteriovorus* HD100 at various timepoints, from 10 minutes to 16 hours, after addition of prey (*Vibrio cholerae*, *Escherichia coli*, or *E. coli* minicells). Compared to thicker cells, the thinness of *E. coli* minicells yielded higher-resolution details about the predator-prey interaction. The small size of the minicells also prevented complete entry of the predator, allowing us to capture otherwise fleeting intermediates in the rapid invasion process. Table S1 lists the number of cryo-tomograms we acquired at each stage of the predatory lifecycle of *B. bacteriovorus*, and Table S2 the number of examples we observed of each of the features described below.

### I Anatomy of the attack-phase *B. bacteriovorus* cell

Our cryo-tomograms of attack-phase *B. bacteriovorus* showed features previously described, including a compact spiral nucleoid occupying the center of the cell (Fig. 1A) (Borgnia et al., 2008; Butan et al., 2011). While the nucleoid excluded ribosomes, they were occasionally abundant at its periphery (Fig. S1 and Movie S1), as seen previously by cryo-ET (Borgnia et al., 2008; Butan et al., 2011). Again consistent with previous cryo-ET of attack-phase cells (Borgnia et al., 2008), we saw unidentified tubes (on average, two per cell) in the cytoplasm. The tubes had a uniform diameter of ∼8 nm and were typically a few tens of nanometers in length (Fig. S2). Each cell contained a single polar flagellum, sheathed in outer membrane (Fig. 1A), and subtomogram averaging of 79 particles revealed that the structure of the flagellar motor is similar to that recently published from a host-independent strain of *B. bacteriovorus* (Chaban et al., 2018) (Fig. 1B).

**Figure 1:**
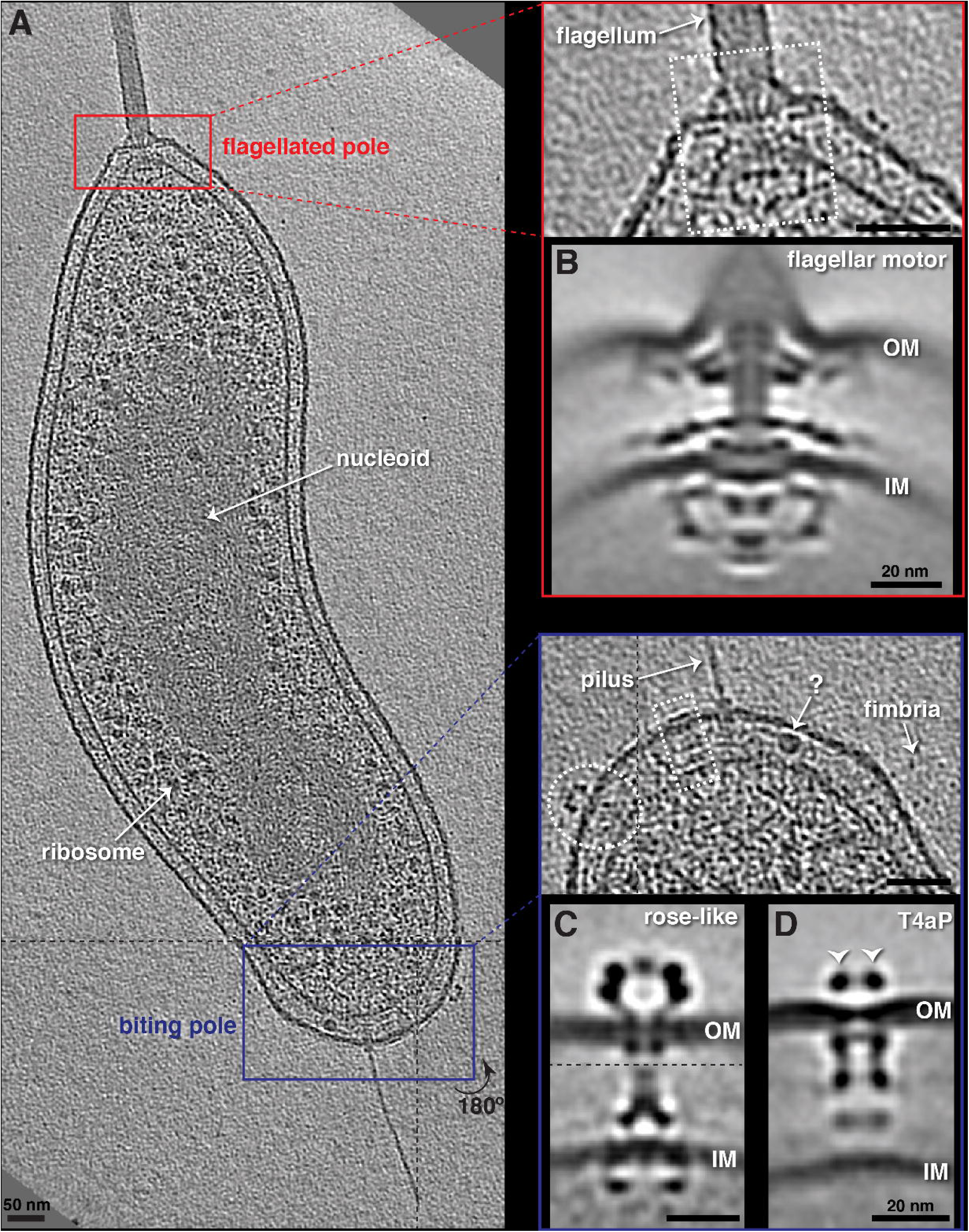
Anatomy of attack-phase *B. bacteriovorus*. **A)** A slice through an electron cryo-tomogram of an attack phase *B. bacteriovorus* cell, with enlarged views of the flagellated (red) and biting (blue) poles. White rectangles highlight the flagellar motor (upper panel) and non-piliated T4aP basal body (lower panel), and the white ellipse highlights a rose-like complex. Question mark points to a cross-section through a periplasmic tubular structure, and white arrow to a type IVa pilus. Scale bars are 50 nm. **B**-**D)** Central slices through subtomogram averages of the *B. bacteriovorus* flagellar motor (B), rose-like complex (C), and non-piliated T4aP basal body (D). White arrows in D point to the extracellular ring of the T4aP basal body. Scale bars are 20 nm. Dashed black lines indicate a composite of slices through the tomogram at different z-heights in (A), and a composite of subtomogram averages aligned on the outer and inner membrane, respectively, in (C). OM = outer membrane, IM = inner membrane.

We also saw novel features, including extracellular vesicles with a uniform diameter of ∼25 nm near cells (Fig. S3A). On the “biting” pole opposite the flagellum, we noted several characteristic features that have not been described before. Rarely, we saw spherical and short filamentous structures in the periplasm (Figs. 1A and S3B). We observed abundant, thin (∼3-4 nm wide) fimbriae on the cell surface, with lengths ranging from ∼50 to more than 100 nm (Fig. S3A&C). We also observed a novel complex, spanning the periplasm and with a prominent extracellular rosette of density. We call this unidentified structure the “rose-like complex.” On average, each cell had 2-3 rose-like complexes at the biting pole (Fig. 1A). Subtomogram averaging of 132 particles revealed a molecular complex spanning the entire periplasmic space with associated cytoplasmic densities (a ring ∼17 nm in diameter) and extensive extracellular densities. Five distinct extracellular densities could be distinguished in cross-section: two stacked rings and a central cap (Fig. 1C).

We observed both piliated and non-piliated T4aP basal bodies on the biting pole (Figs. 1A and S4 and S5), with empty more abundant than piliated (Table S2). A subtomogram average of 335 non-piliated T4aP basal bodies revealed the architecture, including a distinctive extracellular ring present in both non-piliated and piliated basal bodies (Figs. 1D and S4) which was not seen in subtomogram averages of T4P in other species (Chang et al., 2016, 2017; Gold et al., 2015; Treuner-Lange et al., 2020). While all piliated basal bodies had the ring, not all non-piliated basal bodies did (Fig. S5). This could either be because the external ring had disassembled or not yet assembled, or because these structures were not in fact T4aP but rather, e.g. T4bP, which are also involved in adhesion to prey (Avidan et al., 2017). In addition, classifying the non-piliated T4aP using principal component analysis revealed that ∼1/2 of the particles had only the Secretin outer membrane channel and extracellular ring and lacked the lower periplasmic ring (Fig. S6). This might reflect different assembly stages, as it has been shown that T4P in other species utilize an outside-in assembly pathway starting from the Secretin (Friedrich et al., 2014).

### II Attachment of *B. bacteriovorus* to prey

We observed attachment of *B. bacteriovorus* to *V. cholerae* and *E. coli* minicells (of various sizes). The earliest event we identified was connection of *B. bacteriovorus* to prey by T4aP, with the tip of the extended pilus clearly in contact with the prey’s outer membrane. At the resolution of our cryo-tomograms, no distinctive features were visible at the pilus-prey attachment point (Fig. 2A). Occasionally, we also saw thin fimbriae apparently contacting the outer membrane of the prey cell (Fig. S7).

**Figure 2:**
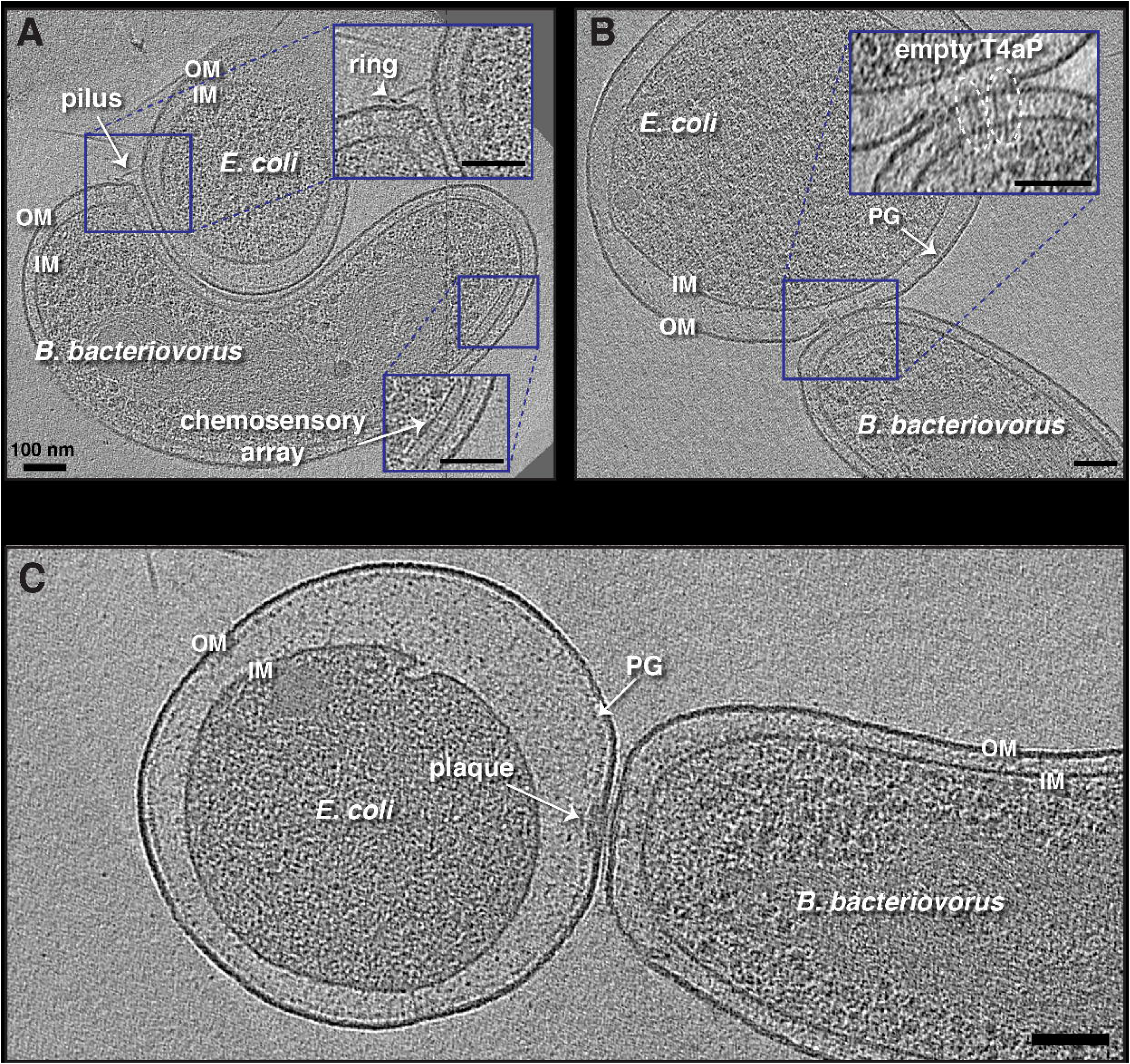
Attachment of *B. bacteriovorus* to prey. **A)** A slice through an electron cryo-tomogram showing a *B. bacteriovorus* cell attached to a prey (*E. coli* minicell) vi a T4aP. OM = outer membrane, IM = inner membrane. **B)** A Slice through an electron cryo-tomogram of *B. bacteriovorus* attached to prey (*E. coli* minicell) showing non-piliated T4aP basal bodies (white ellipses) penetrating to the prey’s PG layer. **C)** A slice through an electron cryo-tomogram of *B. bacteriovorus* attached to prey with a polar attachment plaque. PG = peptidoglycan. Scale bars 100 nm.

Next, we observed predator and prey in close apposition. In some cases, pilus attachments were still visible; in others the pili had retracted completely. In all cases, we observed non-piliated T4aP basal bodies aligned at the contact site, with rose-like complexes nearby (Figs. 2B and S8). In several attachments to *E. coli* minicells, where finer detail could be resolved, we observed that the non-piliated T4aP basal bodies extended through the prey outer membrane, with their extracellular rings apparently embedded in the prey peptidoglycan (PG) (Figs. 2B and S9). At this stage, we began to observe enlargement of the prey periplasm. In agreement with previous studies (Lambert et al., 2011), with rod-shaped prey the contact site was usually located on the side of the prey cell, and on the pole of the predator (Fig. S10A). In some cases, particularly with smaller, spherical *E. coli* minicells, the attachment site was displaced slightly off the biting pole onto the side of the predator (Fig. S10B-D).

In the next stage, we observed an electron-dense but relatively unstructured plaque of material (henceforth referred to as the “attachment plaque”) at the contact point between the predator and the prey. The diameter of this plaque ranged from 15-70 nm and the thickness usually extended from the predator’s OM to the prey’s PG cell wall (Figs. 2C, S10C and D), suggesting a modification of the prey cell wall at the predator-prey contact point in agreement with previous reports (Kuru et al., 2017). We often observed nonpiliated T4aP basal bodies near or in the plaque (Fig. S10D), as well as rose-like complexes and, very occasionally, prey-attached pili nearby. While we sometimes observed two or even three *B. bacteriovorus* attached to the same prey cell with T4aP, only one ever formed an attachment plaque, consistent with the committed attachment observed in previous studies (Fig. S11).

At around the same time that the attachment plaque formed at the biting pole, an unexpected process began at the other pole. The sheathed flagellum of the *B. bacteriovorus* was resorbed into the periplasmic space, wrapping around the cell. The process seems to initiate with breakage of the flagellar motor at its OM-embedded ring (the L- (lipopolysaccharide) ring, Fig. S12). The L-ring remained in the OM. Early in the process, the P- (peptidoglycan) ring was still visible around the flagellar rod but was located more than 20 nm from the OM, compared to 10-11 nm in attack-phase cells (Figs. S13 and S14 and Movie S2). Consistent with a decoupling of the P- and L-rings, we observed two examples of *B. bacteriovorus* cells attached to prey with a plaque and with disrupted OM around the flagellum; in both cases, only the P-ring, and not the L-ring, was visible surrounding the rod of the motor (Fig. S15).

As flagellar resorption continued, the motor (lacking the L-ring but including the P-ring and rod) was completely internalized to the periplasm and moved off the cell pole, with the hook and basal portion of the filament entering the periplasm (Figs. 3, S16 and Movie S3). The L-ring still encircled the filament at the junction of OM and flagellar sheath (Figs. 3, S16 and Movie S3). Eventually, most or all of the filament was internalized to the periplasm, wrapping around the cell (Figs. S17, S18 and Movies S4-S7). In some cases, we saw the motor further up the side of the cell. In other cases, we could not find the motor, either because it was degraded or because it was located in a part of the cell not visible due to the effect of the missing wedge of information in cryo-ET (Baumeister, 1999). During the absorption process, the exit point of the flagellum sometimes shifted from the pole up the side of the cell (Figs. S18, S19 and Movie S6). In cells with fully internalized filaments, we could no longer identify L-rings in the outer membrane. In some cases, the wrapped filament broke in the periplasm (Movie S4). Figure S20 summarizes this absorption process based on our cryo-ET data.

**Figure 3:**
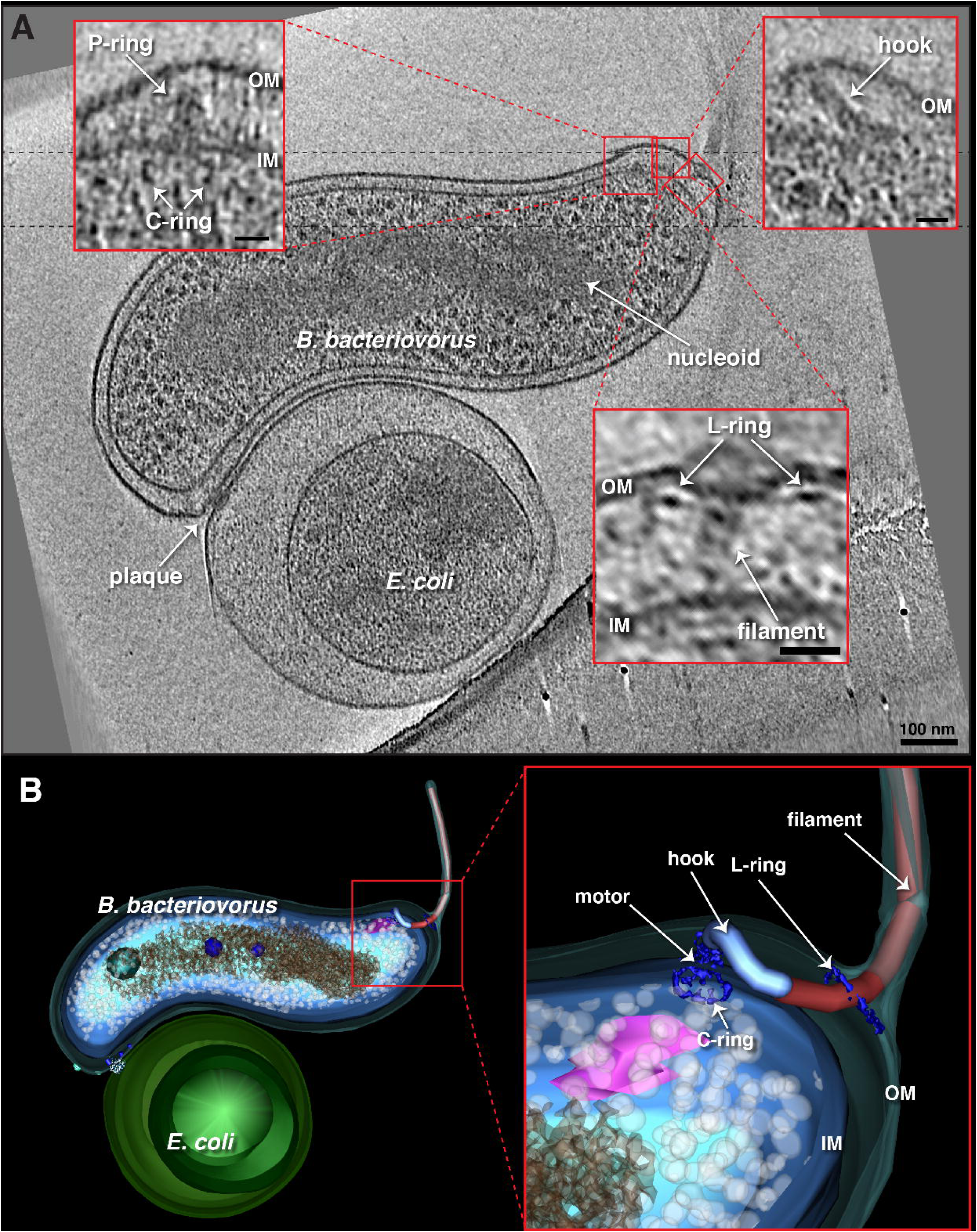
*B. bacteriovorus* flagellar absorption. **A)** A slice through an electron cryo-tomogram showing a *B. bacteriovorus* cell attached to a prey via an attachment plaque with its flagellum resorbing into the periplasm. Enlargements in the red-boxed areas highlight different parts of the absorbed flagellum. Scale bar is 50 nm in the main panel, 20 nm in the enlargements. **B)** A 3D segmentation of panel (A) and an enlarged view illustrating different parts of the absorbed flagellum.

Chemosensory arrays remained visible throughout the attachment phase, although they appeared to be partially degraded (smaller in diameter than in attack-phase cells) in some cases. As in attack-phase cells, we observed small, uniformly-sized membrane vesicles in the vicinity of the predator and attached prey (Fig. S21). In some vesicles, densities were visible either inside and/or on the surface (Fig. S21F). We also saw vesicles near the sheath of the flagellum during resorption (Fig. S22).

### III Invasion of the prey periplasm

We captured 18 cryo-tomograms of stalled invasions of *B. bacteriovorus* entering the periplasm of *E. coli* minicells. Note that in these stalled invasions we do not know how long before sample freezing a particular predator entered its prey or whether the non-permissive size of the small prey had incidental effects on the entry process. Still, this paradigm provided an unique opportunity to view fleeting stages of invasion at high resolution. In invasion, the attachment plaque at the contact site was replaced by a portal structure through which the predator entered the prey periplasm (Figs. 4, S23, and Movie S8). This portal ring, which bridged the outer membranes of predator and prey, appeared in cross-section as a thin (<5 nm), dark density extending from the prey’s PG layer to the outside of the cell, capping the open end of the prey outer membrane (Figs. 4A and S23A). The height of the portal in cross-section, on the order of a few tens of nanometers, varied between cells, and even on opposite sides of the same cell (e.g. compare Figs. 4A and S23A). In eight examples, we observed what appeared to be prey OM blebbing out from the portal (Movie S9). Consistent with a water-tight seal model, the portal appeared to exert considerable force on the *B. bacteriovorus* cell, as previously observed (Abram et al., 1974), reducing the distance between the outer and inner membranes by ∼50% at the entry point compared to elsewhere in the cell, and constricting the deformation-resistant cell wall (Figs. 4B-C, S23B-C). Consistent with previous reports of cell flexibility (Borgnia et al., 2008), we observed that *B. bacteriovorus* could bend considerably to maximally occupy the prey periplasm (Fig. 4A).

**Figure 4:**
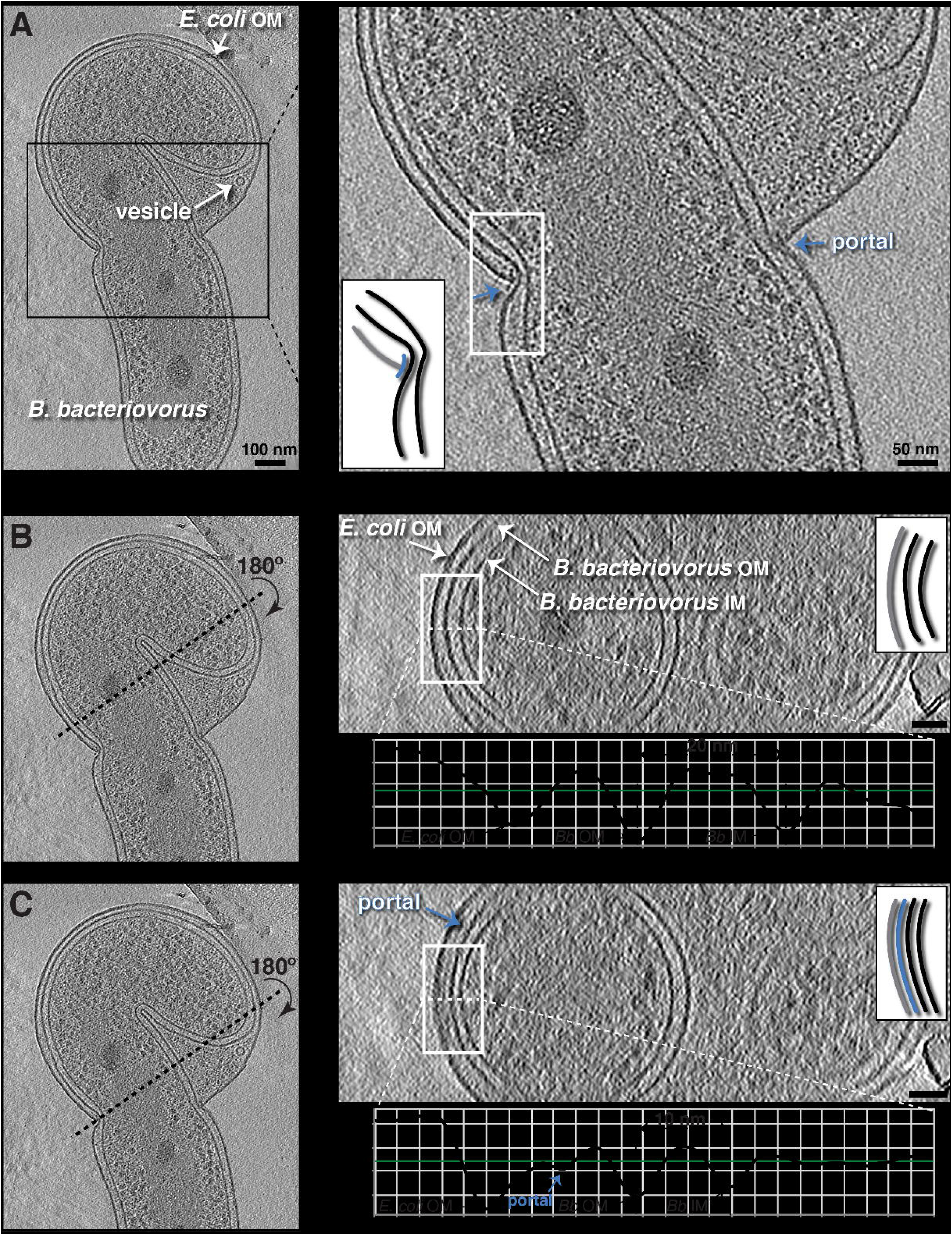
*B. bacteriovorus* prey invasion. **A)** A slice through an electron cryo-tomogram (left) and enlarged view (right) showing a stalled invasion by a *B. bacteriovorus* of an *E. coli* minicell. Blue arrows and inset schematic highlight the portal. **B)** Right (top): Cross-section through the *yz* plane of the tomogram shown in (A) along the black dotted line indicated on the left. Note that this slice does not include the portal. Right (bottom): Average density profile taken along the white dashed line (inside the white rectangle) in the top panel. The distance between the predator’s inner (IM) and outer membranes (OM) is indicated (20 nm). **C)** Similar to (B) but for a *yz* slice where the portal is visible. The distance between the predator’s inner and outer membranes is indicated (10 nm). The schematics in the right panels of (A-C) represent the white-boxed areas in the corresponding slices, with the portal shown in blue, the prey outer membrane in grey, and the predator inner and outer membranes in black. Scale bars 100 nm in the left panel of (A), and 50 nm in other panels.

In all stalled invasions, we observed that the *B. bacteriovorus* cells had fully degraded their absorbed periplasmic flagella and lacked rose-like complexes and fimbriae. Chemosensory arrays were also partially or completely degraded (Fig. S24). Interestingly, while complexes morphologically similar to non-piliated T4aP basal bodies were still present in the predator, they lacked the characteristic extracellular ring found at earlier stages of invasion (Fig. S25), and were less abundant than T4aP basal bodies in attack-phase cells. This could be because either the extracellular ring is lost during the invasion process, or these are different complexes, e.g. type II secretion systems.

In two examples where a non-flagellated *B. bacteriovorus* cell was in the vicinity of a lysed prey, we observed knob-like densities on the predator’s biting pole (Fig. S26). Given the lysed prey cell nearby and the lack of attack-phase structures such as flagella or chemosensory arrays in the *B. bacteriovorus*, we think it likely that these cells were pulled out of the prey post-invasion during sample preparation. When present, the knob-like structures could be abundant; we identified 23 examples on one cell (Table S2). While leg-like densities could be seen extending from the extracellular domains to the PG layer in individual examples, subtomogram averaging failed to resolve a consistent structure, suggesting flexibility or differing stoichiometry.

Even though entry was incomplete, *B. bacteriovorus* in stalled invasions of *E. coli* minicells showed signs of entering growth phase. Some predators had at least partially decondensed their nucleoids, and the prey cytoplasm was considerably reduced in size, presumably consumed by the predator. Perhaps related to this digestion, we observed multiple vesicles with a consistent size of ∼25 nm in the prey periplasm (Figs. 4A and S23A). We cannot tell whether they originated from prey or predator membrane, but their size is identical to those we observed in the vicinity of isolated attack-phase and prey-attached *B. bacteriovorus*.

### IV Growth phase in the bdelloplast

We captured 54 cryo-tomograms of the bdelloplast stage from samples of *B. bacteriovorus* invading *V. cholerae* cells and *E. coli* minicells large enough to accommodate the entire predator. Once the predator fully entered the prey’s periplasm, the entry hole was sealed by a scar consisting of an extracellular bubble of what appeared to be membrane and an amorphous electron density associated with the prey OM and PG beneath it (Figs. 5, S27 and S28 and Movies S10-S13). In invaded *V. cholerae*, the prey’s (non-functional) flagellum remained attached to the bdelloplast (Figs. 5F and S27B, C). The flagellum remained connected to the part of the motor embedded in the OM and PG: the PL-rings and part of the rod. Presumably these parts were separated from the rest of the motor by periplasmic expansion. We did not observe any motor components still associated with the inner membrane. The fact that flagellar relics remained only on *V. cholerae* bdelloplasts, and not *E. coli*, could be because the flagellar sheath aids in retention. In *V. cholerae* bdelloplasts, we sometimes also observed PL-subcomplexes (without associated filaments) resulting from previous flagellar loss events (Ferreira et al., 2019; Kaplan et al., 2020) (Fig. S27B).

**Figure 5:**
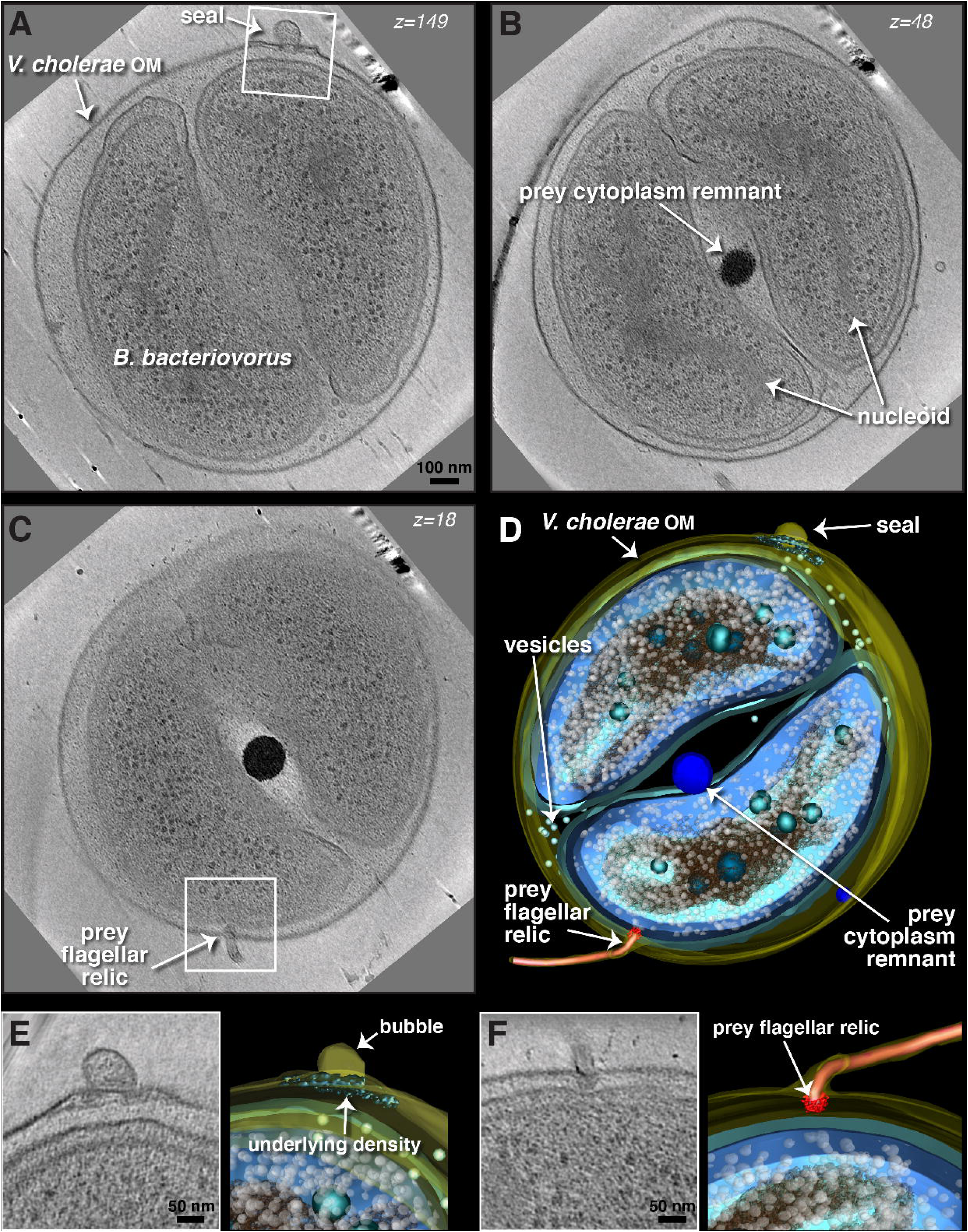
Anatomy of the bdelloplast. **A-C)** Slices (at different z-levels) through an electron cryo-tomogram of a *V. cholerae* bdelloplast containing two *B. bacteriovorus* after predator division. **D)** A 3D segmentation of the bdelloplast shown in (A-C). **E)** Enlargement (left) and 3D segmentation (right) of the white-boxed area in (A), highlighting the features of the seal. **F)** Enlargement (left) and 3D segmentation (right) of the white-boxed area in (C), rotated 180°, highlighting the prey flagellar relic. OM= outer membrane. Scale bars 100 nm in (A-C) and 50 nm in (E, F).

In predators inside bdelloplasts, we saw neither chemosensory arrays nor any relics of flagella, although we occasionally observed filamentous structures in the periplasm that may be remnants of flagellar digestion (Fig. S29). We also sometimes observed these in (flagellated) attack-phase cells (Fig. S3B). Interestingly, they were always located at the pole (the biting pole of attack-phase cells), perhaps reflecting spatial differences in proteolysis. We also could not find any rose-like complexes or T4aP basal bodies with external rings in *B. bacteriovorus* inside bdelloplasts. As in stalled invasions, we did identify putative nonpiliated T4aP basal bodies lacking the external ring. Again, they were less abundant than in attack-phase cells; from 47 cryo-tomograms of stalled invasions or early bdelloplasts, we identified 25 such particles (Table S2). We did occasionally observe 8-nm-wide cytoplasmic tubes, as seen in other lifecycle stages (Figs. S30). In addition, we saw variously-sized spherical, nested and horseshoe-shaped vesicles in the predator’s cytoplasm, morphologically similar to those reported in other species (Dobro et al., 2017) (Fig. S31).

As in stalled invasions of *E. coli* minicells, we observed many uniformly-sized (∼25 nm) vesicles in the bdelloplast periplasm (Figs. 5 and S31 and Movies S11-S13). Consistent with active growth, *B. bacteriovorus* nucleoids were less condensed than in earlier stages and, in concert with the prey cytoplasm shrinking, the predator cell elongated and curled to fill most of the bdelloplast (Fig. S32). Contrary to previous observations by traditional EM (Abram et al., 1974), we could not unambiguously identify a connection between the predator OM and prey inner membrane (Figs. S28, S32, S33 and Movies S10-S11).

When nearly all of the prey cytoplasm was consumed, the elongated *B. bacteriovorus* cell divided. The number of progeny depends on the size of the bdelloplast (Fenton et al., 2010a), and we observed two or three progeny cells in *E. coli* and *V. cholerae* prey (e.g., Fig. 5). In a few cases, division produced an extra, small spherical product, in accordance with previous reports (Burnham et al., 1970) (Fig. S34). In some cases, bdelloplasts contained a very dense sphere of material, presumably containing the remnants of the prey cytoplasm (Figs. 5 and S34). In other cases, not even this remained (a characteristic we use to define an “end-stage bdelloplast”). The characteristic ∼25 nm vesicles, however, were still present in end-stage bdelloplasts even after no prey cytoplasm remained.

In some elongated or divided *B. bacteriovorus* in end-stage bdelloplasts of *E. coli*, we observed a remarkable hexagonal lattice of ribosomes coating the nucleoid (Fig. 6 and Movies S13-S17). The lattice spacing was ∼20 nm, consistent with maximally dense packing of ribosomes (Fig. S35). This arrangement was much more extensive than we and others observed in attack-phase cells (Butan et al., 2011). We measured the distances from ribosomes to the apparent surface of the nucleoid in two tomograms. Of 2,304 ribosomes in one (Fig. 6) and 1,109 in the other (Fig. S36), ∼80% were located within 10 nm of the nucleoid surface. By comparison, in a simulation of the same number of randomly-packed 20-nm spheres in the same tomographic volumes (see Materials and Methods), only ∼20-25% were expected to be located within 10 nm of the nucleoid surface (Figs. 6E and S36), suggesting that the association we observed does not arise simply by chance. To investigate whether the ordered ribosomes shared the same orientation, we produced an ∼4.7 nm-resolution subtomogram average of the ribosomes in a *B. bacteriovorus* cell and mapped it back into the tomographic volume using the positions and orientations determined during averaging. We found that individual particles were apparently randomly oriented on the nucleoid surface (Fig. S37).

**Figure 6:**
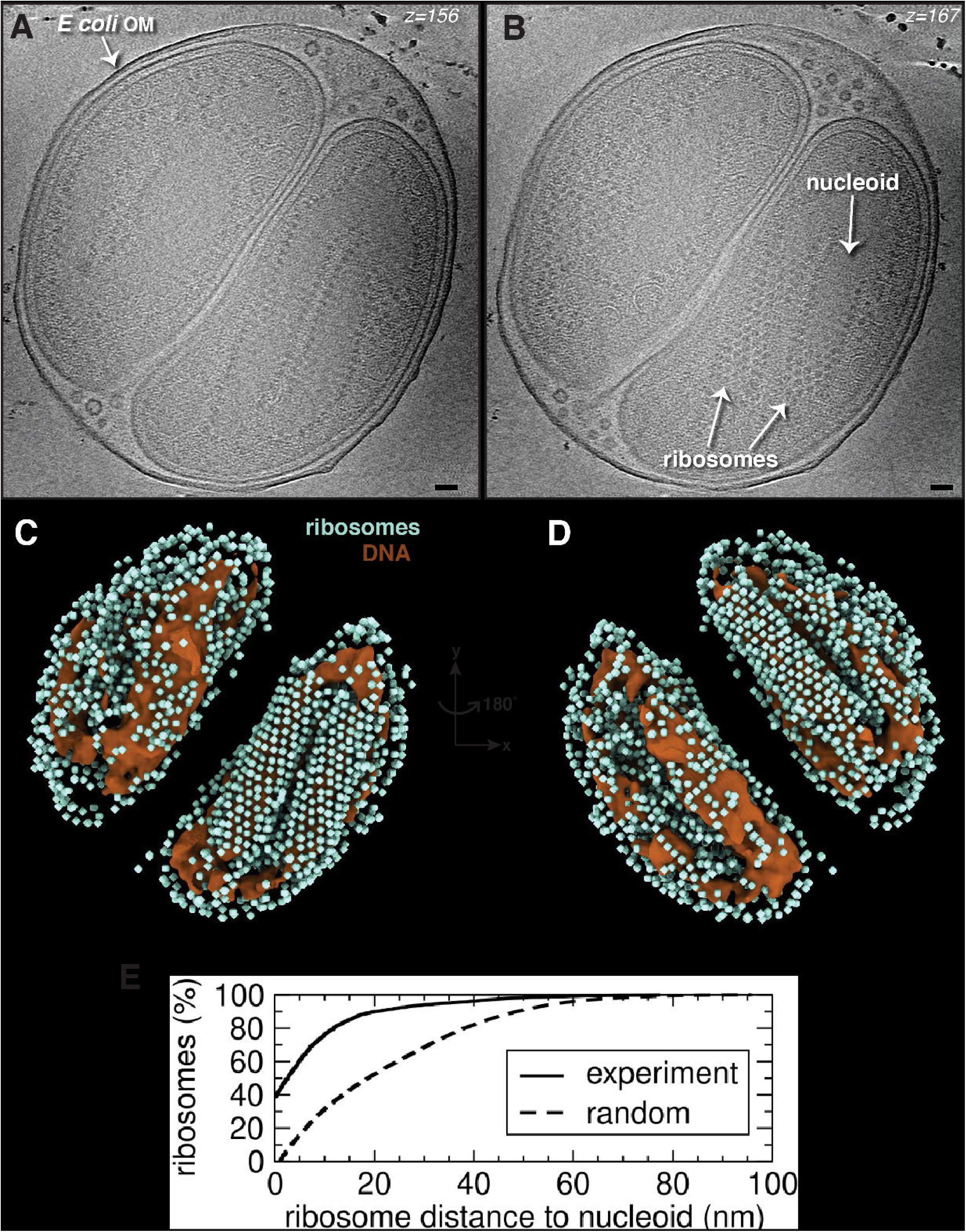
Ribosomal nucleoid lattice in the end-stage bdelloplast. **A, B)** Slices (at different z-levels) through an electron cryo-tomogram of an end-stage *E. coli* bdelloplast containing two *B. bacteriovorus* cells after predator division, highlighting the hexagonal arrangement of ribosomes around the nucleoids. **C, D)** Rotated views of a 3D segmentation of the nucleoids and ribosomes of the cryo-tomogram shown in (A, B). Scale bars 50 nm. **E)** Distances of individual ribosomes from the nucleoid surface measured in the 3D segmentations of the cryo-tomogram shown in (A&B) (“experiment,” solid line), compared to a simulation of randomly distributed 20 nm-wide spheres packed in the same segmented volume (“random,” dashed line). See also Movie S16.

Movie S18 offers an animated summary of all stages of the *B. bacteriovorus* predatory lifecycle that we observed in this study.

## Discussion

Here we used cryo-ET imaging to reveal the predation cycle of *B. bacteriovorus in situ* at nanometer-scale resolution (Movie S18). Our results contextualize decades of research on BALO predation and uncover many surprising new details of the process.

In addition to previously-characterized structures in attack-phase cells such as the flagellum, poly-phosphate storage granules, highly-condensed nucleoid, chemosensory array, and type IV pili (Borgnia et al., 2008; Butan et al., 2011; Chaban et al., 2018; Evans et al., 2007; Mahmoud and Koval, 2010), we observed several unidentified structures. At all stages of invasion, *B. bacteriovorus* cells contained 8 nm-wide cytoplasmic tubes, typically two per cell. The identity and function of these tubes, which were also seen previously by cryo-ET in attack-phase cells (Borgnia et al., 2008), remains unknown, but they may serve a cytoskeletal role. In a few attack-phase cells, we observed spherical or tubular structures in the periplasm at the biting pole which might be fragments of digested flagella. The biting pole of attack-phase cells also contained abundant fimbriae, shorter (∼100 nm or less) and thinner than T4aP and without obvious machinery at their base. Their location suggests a role in prey interaction. Interestingly, mutant strains of *B. bacteriovorus* lacking either T4aP and T4bP genes can still attach to prey (Avidan et al., 2017; Milner et al., 2014); perhaps these fimbriae mediate such adhesion.

The most intriguing new structure we observed is what we call the rose-like complex, present in ∼2-3 copies on the biting pole of nearly every attack-phase cell we imaged. The periplasmic portion of the rose-like complex is morphologically similar to a recent structure of a tripartite efflux pump (Alav et al., 2021) and since related *B. bacteriovorus* type I secretion systems (T1SS) secrete enzymes that modify the prey during invasion (Rendulic, 2004), it is possible that the rose-like complex is a T1SS. Consistent with a role in early invasion, we observed rose-like complexes on attack-phase cells and at prey contact sites, but not in cells during or after invasion. The function of the elaborate extracellular domains extending nearly 20 nm out from the cell is of particular interest; perhaps they interact with, or breach, the prey envelope.

Another machine with a notable extracellular domain is the T4aP basal body. The pili observed on the biting pole of *B. bacteriovorus* in early micrographs (Abram and Davis, 1970; Abram et al., 1974; Shilo, 1969) were previously identified as T4aP by mutant analysis and immunolocalization (Evans et al., 2007; Mahmoud and Koval, 2010). Our higher-resolution imaging here revealed a novel extracellular ring surrounding the base of the pilus, not seen in previous subtomogram averages of related T4aP in *Thermus thermophilus* and *Myxococcus xanthus* (Chang et al., 2016; Gold et al., 2015). The ring was also present in non-piliated basal bodies, indicating that it is stable in the absence of the pilus. Interestingly, these extracellular densities were observed on attack-phase cells in a previous cryo-ET study, but their relation to T4aP was not resolved (Borgnia et al., 2008). We observed some non-piliated complexes lacking the extracellular ring, both in attack-phase cells and in bdelloplasts, where no complexes with the outer ring were observed. It is possible that these complexes are not T4aP, but rather a related machine containing a Secretin pore in the outer membrane, such as a type II secretion system, which is structurally similar (Ghosal et al., 2019b) and thought to be involved in *B. bacteriovorus* secretion (Dori-Bachash et al., 2008; Rendulic, 2004). In addition, *B. bacteriovorus* also contains a large repertoire of T4bP genes, which are dispensable for attachment but required for invasion (Avidan et al., 2017; Schwudke et al., 2005). Their products have not been located on the cell and it is possible that some or all of the ring-less basal bodies we saw were T4bP. Alternatively, the extracellular ring may be a transient component of the *B. bacteriovorus* T4aP, perhaps dissociating during the prey entry process.

The pilin protein PilA is required for invasion and T4aP have been suggested to pull cells into prey, perhaps by attaching to the cell wall (Evans et al., 2007; Mahmoud and Koval, 2010; Milner et al., 2014), but mutants lacking the disassembly PilT ATPase are still capable of invasion (Chanyi and Koval, 2014) and the role of T4aP in the process remains a major open question in the field (Sockett, 2009). In our tomograms, multiple T4aP can be seen attached to prey cells and clearly exerting force as they retracted, pulling the prey OM and PG into close contact with the predator. As the membranes were brought into contact, the shortening pili fully disassembled, leaving empty basal bodies. Interestingly, these non-piliated basal bodies continued to mediate attachment and could be seen extending through the prey OM, with their extracellular rings located in the PG layer of the prey, suggesting the function of this novel component. Our results thus suggest that pili themselves do not drive entry, but rather force the initial connection. If the basal bodies without external rings we observed in bdelloplasts were in fact T4aP, it is possible that the rings remained embedded in the prey PG.

With a few exceptions noted by (Lambert et al., 2006), *B. bacteriovorus* are known to lose their flagella when entering prey. Our images reveal a surprising mechanism: while attached to a prey cell, the flagellar motor is broken at the L-ring and the filament absorbed into the predator’s periplasm, where it is digested. This mechanism differs from all previous observations of flagellar loss due to lifecycle-programmed ejection, response to nutrient deprivation or mechanical breakage, all of which leave a stable subcomplex of the P- and L-rings in the cell wall and outer membrane (Ferreira et al., 2019; Kaplan et al., 2019, 2020, 2021b, 2021b; Zhu and Gao, 2020; Zhu et al., 2019; Zhuang and Lo, 2020; Zhuang et al., 2020). It will be interesting to see whether the *B. bacteriovorus* motor lacks the inter-subunit interactions that likely stabilize the PL-subcomplex in other species (Johnson et al., 2021; Tan et al., 2021; Yamaguchi et al., 2020). This process also differs from the breakage of the prey flagellum that occurs as the periplasmic space is expanded into the bdelloplast. In that case, we see that the flagellum remains anchored to the cell by a stub of the motor embedded in the outer membrane. Why the hook/filament is not lost is unclear; perhaps it is locked into the remodeled PG. This process is reminiscent of that recently observed in other species upon cell lysis, where the cytoplasmic flagellar switch complex is lost, while the periplasmic and extracellular components remain (Kaplan et al., 2021c).

How does *B. bacteriovorus* absorb its flagellum into the periplasm? It is unlikely to be pulled from the motor, which we occasionally saw drift partway up the side of the cell before being fully degraded. Our observation of filaments partially absorbed up to a junction on the side of the cell suggests that the process involves zippering of the flagellar sheath and the outer membrane. Perhaps this is related to the unique lipid composition of the sheath, which is predicted to be even more fluid than the rest of the outer membrane (Thomashow and Rittenberg, 1985b). The vesicles we observed in the vicinity of some absorbed flagella are consistent with previous reports that rotation of sheathed flagella can lead to shedding of outer membrane vesicles (Aschtgen et al., 2016; Brennan et al., 2014). However, we do not know whether the vesicles formed because the motor was still rotating during absorption or due to the absorption process itself.

Other bacterial species have been observed to wrap their extracellular (unsheathed) flagella around themselves (Alirezaeizanjani et al., 2020; Cohen et al., 2020; Constantino et al., 2018; Hintsche et al., 2017; Kühn et al., 2017; Tian et al., 2022). This behavior, which *Shewanella putrefaciens* uses to burrow back out of a tight spot when stuck, is triggered by a mechanical instability in the flagellum that buckles it when the cell can no longer move to alleviate the torque of flagellar rotation (Kühn et al., 2017). Such a situation likely occurs when a *B. bacteriovorus* cell attaches to a prey. Brief continuing flagellar rotation could conceivably wrap the filament around the cell, where fusion of the outer membrane and sheath would bring it into the periplasm. Such increased resistance is consistent with the lateral shift of the biting pole we sometimes observed on attached *B. bacteriovorus* cells. Interestingly, it was recently shown that a discontinuous flagellar filament formed by two different flagellins facilitates screw-like motility in *S. putrefaciens* (Kühn et al., 2018), and is key to wrapping in *Campylobacter jejuni* (Cohen et al., 2020). Multiple flagellins similarly make up distinct segments of the *B. bacteriovorus* flagellum (Thomashow and Rittenberg, 1985a); the reason for this was unknown but now we speculate that it may facilitate flagellar recycling.

A major open question is the nature of the pore through which *B. bacteriovorus* enter their prey. Secreted PG-remodeling enzymes, presumably part of the dense plaque we observe at the attachment site that extends to the prey cell wall, are known to create, and subsequently seal, a reinforced circular porthole in the prey PG (see figures 2 and 3 in (Kuru et al., 2017)). How the prey outer membrane is modified remains more of a mystery. Prey cells remain intact and transcriptionally active throughout the initial entry process (Lambert et al., 2010a), so it was proposed that the membranes of prey and predator must fuse (Negus et al., 2017). Instead, we observed what seems to be a proteinaceous collar curving out from the prey PG to seal the hole in the outer membrane and prevent interaction of the two membranes. Presumably this structure is associated with the PG remodeling enzymes, and may even be a modified and reinforced extension of the cell wall, as suggested by (Abram et al., 1974). The portal is dynamic, expanding and contracting to match the cross-section of the cell passing through it, and its height varied between cells (and even on opposite sides of the same cell). As was also seen by traditional thin-section TEM (Abram et al., 1974), the portal exerted considerable force on the *B. bacteriovorus* cell, deforming its PG layer, consistent with a water-tight seal between the cells.

What provides the force for entry remains enigmatic. Over several decades, various models have been proposed, including flagellar rotation (Stolp and Starr, 1963), retraction of T4aP attached to the prey cell wall (Evans et al., 2007; Mahmoud and Koval, 2010), and attachment to the prey inner membrane as osmotic pressure rapidly expands the periplasm (Abram et al., 1974). Our results rule out all of these models. We see that the *B. bacteriovorus* flagellum is broken, and at least partially absorbed, prior to entry. Similarly, pili are fully disassembled prior to entry. And finally, we see prey periplasmic expansion even before entry, presumably due to secreted enzymes that de-crosslink and sculpt the prey PG during entry (Lerner et al., 2012), with no visible connection between the predator and the now-distant prey inner membrane. It is possible that an expanding periplasmic space provides a suction force. It is also possible that an active mechanism walks the connection point with the portal down the *B. bacteriovorus* cell.

Following predator entry, we saw that the portal is sealed by a scar of electron dense material, likely related to the resealing of the PG sacculus (Kuru et al., 2017; Snellen and Starr, 1974), with a protruding bubble of what appears to be membrane. While previous, traditional EM imaging also observed an electron-dense ring at the sealed penetration pore (Shilo, 1969), no associated bubble was observed, perhaps reflecting the improved preservation and hydration of membrane structures by cryo-EM. Given the blebs around the invasion portal we noticed on some prey cells in stalled invasions, it may be prey OM. Alternatively, it may be outer membrane pinched off from the end of the invading predator. *B. bacteriovorus* has a highly labile OM, as we saw both with absorption of the sheathed flagellum and in occasional highly-curved cells where the outer membrane fused into a sac, and consistent with previous lipid analysis of its unique membrane (Lambert et al., 2008). In either case, the bubble must be somehow tethered to the seal, perhaps through membrane-embedded protein complexes.

While we did not see the connections to the prey cytoplasm during entry observed by (Abram et al., 1974), we did see a few examples of the predator outer and prey inner membranes in close proximity in the bdelloplast. In most cases, however, we only observed vesicles in the bdelloplast periplasm, suggesting that this may be a mechanism for nutrient transfer. Compared to membrane vesicles of other species (Kaplan et al., 2021d; Toyofuku et al., 2019), these vesicles were notable for their relatively small and uniform size (∼25 nm in diameter). Similar small periplasmic vesicles may be present in previous traditional EM images, but interpretation is limited by the membrane disruption of that sample preparation (Abram et al., 1974). In our tomograms, we cannot tell which cell(s) are producing the vesicles in bdelloplasts, but we also observed them near isolated attack-phase cells as well as budding from *B. bacteriovorus* attached to prey. We also observed them in end-stage bdelloplasts where no remaining prey inner membrane was visible. *B. bacteriovorus* is known to translocate a pore protein from its outer membrane to the prey inner membrane (Tudor and Karp, 1994); perhaps similar transferred machinery produces vesicles to deliver prey cytoplasmic content, and lipids, to the growing predator.

Following complete consumption of the prey, we observed a striking hexagonal lattice of ribosomes on the surface of the *B. bacteriovorus* condensed nucleoid(s). Similar, though more limited, ribosome associations were previously observed in attack phase, particularly in a *B. bacteriovorus* mutant with an even more tightly condensed nucleoid than wild-type cells (Borgnia et al., 2008; Butan et al., 2011), and in some attack-phase cells in this study. One possibility for this ribosome lattice is that it is a depletion effect resulting from entropic forces encouraging large objects to aggregate on a surface (Asakura and Oosawa, 1954, 1958; Rocha et al., 2020). However, we did not observe the effect on other cell surfaces such as the inner membrane, nor in other stages of the cell cycle, so we think this unlikely. Eukaryotic ribosomes have been observed to crystallize in hypothermic conditions (Byers, 1967), but we see no relationship between the orientation of neighboring particles consistent with a crystal, or with a polysome (Brandt et al., 2009). We think a more likely possibility is that the ribosomes are independently translating transcripts from the condensing nucleoid. The switch from growth phase to attack phase involves a transcriptional shift activating a few hundred genes, and inactivating many more (Karunker et al., 2013; Lambert et al., 2010b). The highly condensed attack-phase nucleoid excludes even small monomeric proteins (Kaljević et al., 2021), so the ribosomes we observe on the surface may be, or recently have been, coupled to transcribing polymerases.

Together, our results provide the most complete view to date of the unique structures that enable the complex lifecycle of this endobiotic bacterial predator, and raise new questions. We hope our work spurs further study of this fascinating process and informs potential future applications of bacterial predators as living antibiotics (Atterbury and Tyson, 2021).

## Acknowledgements

This project was funded by the National Institutes of Health (grant R01 AI127401 to G.J.J) and a Baxter postdoctoral fellowship from Caltech to M.K. S.K. is supported by the Swedish Research Council (2019-06293). Cryo-ET work was performed in the Beckman Institute Resource Center for Transmission Electron Microscopy at the California Institute of Technology and the Howard Hughes Medical Institute Janelia Farm CryoEM Facility. We thank Daniel Villanueva Avalos for making the summary animation. We are deeply grateful to Prof. Liz Sockett (University of Nottingham) for the gift of the *B. bacteriovorus* strain and helpful advice and comments.

