## Supplementary file for "Dynamic structural adaptations enable the endobiotic predation of *bdellovibrio bacteriovorus*"

### Supplementary Figures

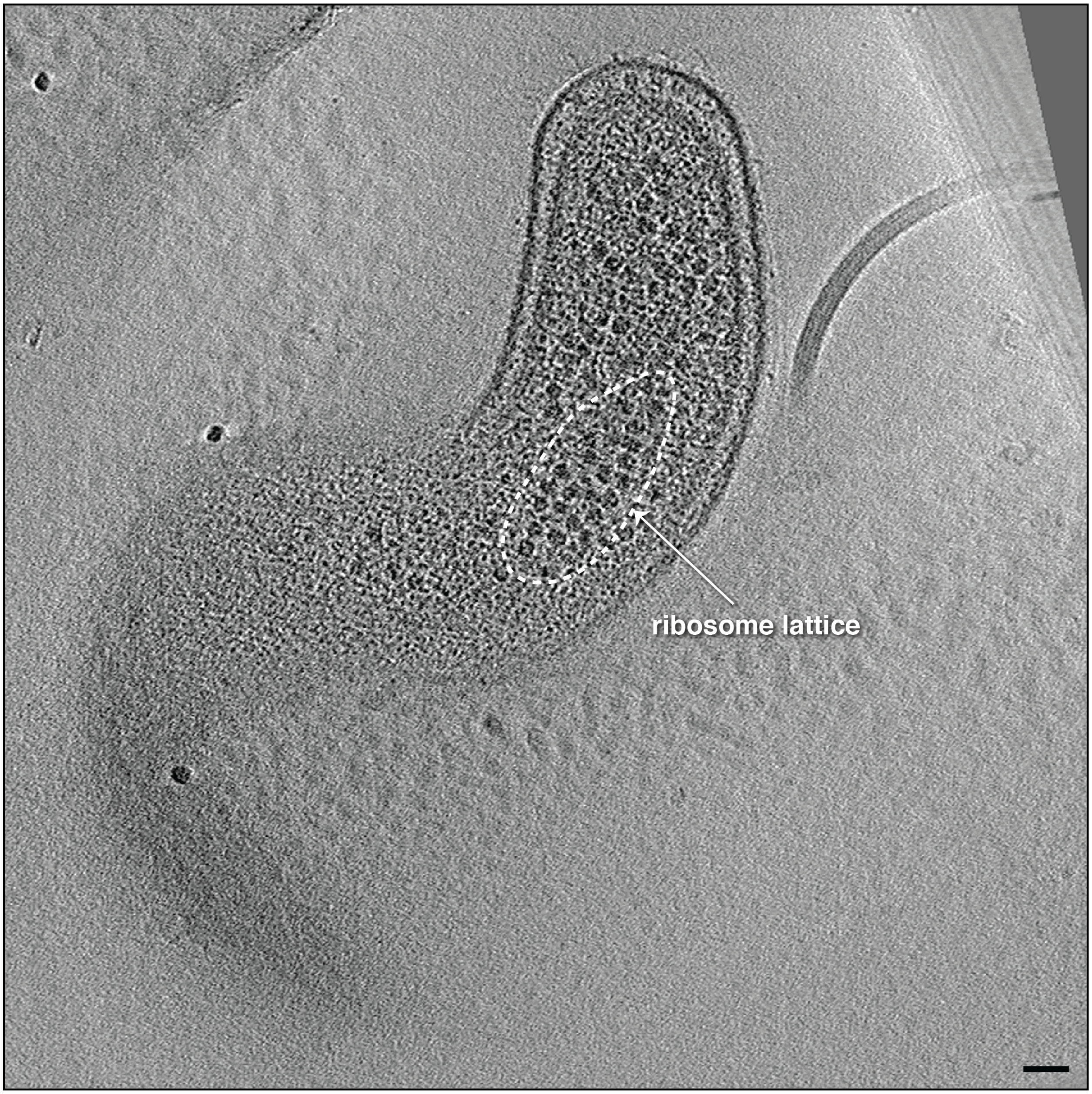


**Figure S1:** A slice through an electron cryo-tomogram of an attack-phase *B. bacteriovorus* cell illustrating a local regular arrangement of the ribosomes (white ellipse). Scale bar is 50 nm.

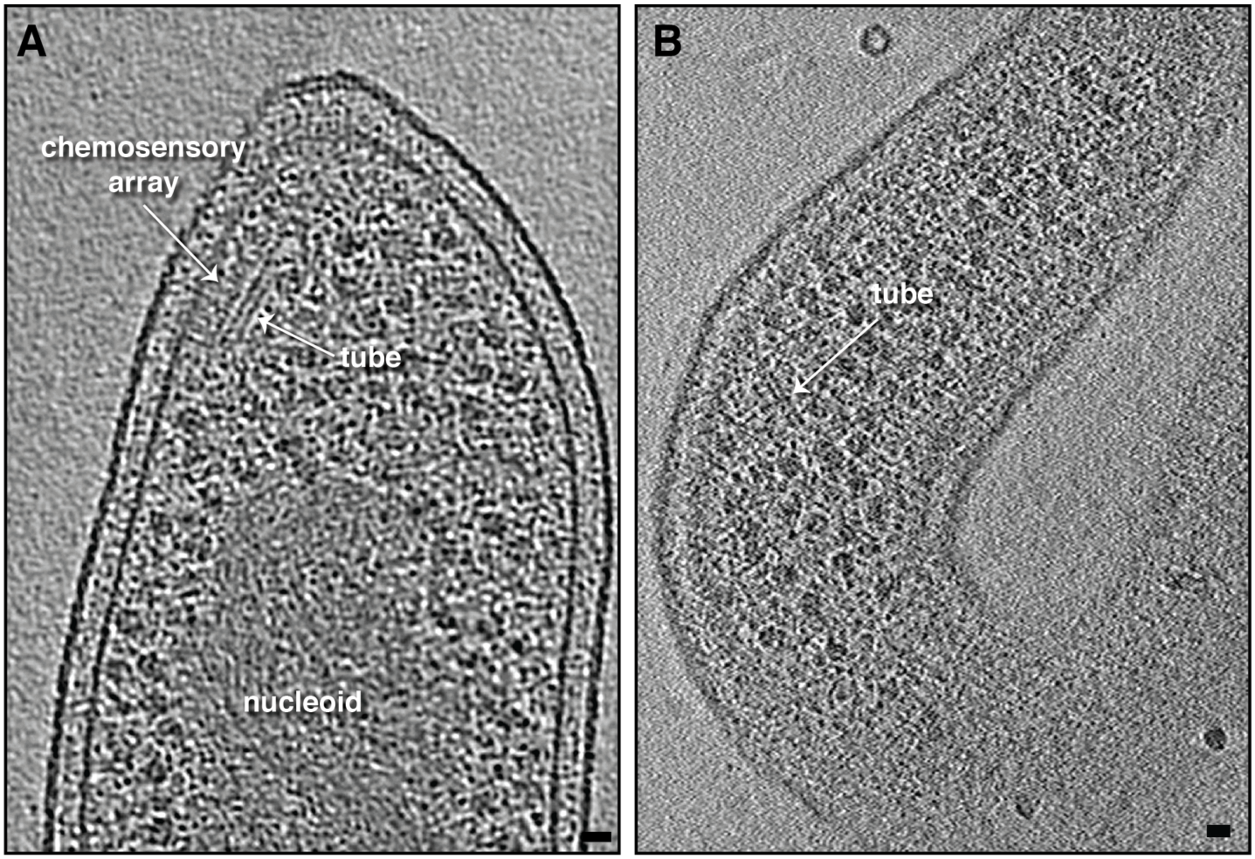


**Figure S2: A-B)** Slices through electron cryo-tomograms of *B. bacteriovorus* attack-phase cells highlighting cytoplasmic tubes. Scale bar is 20 nm.

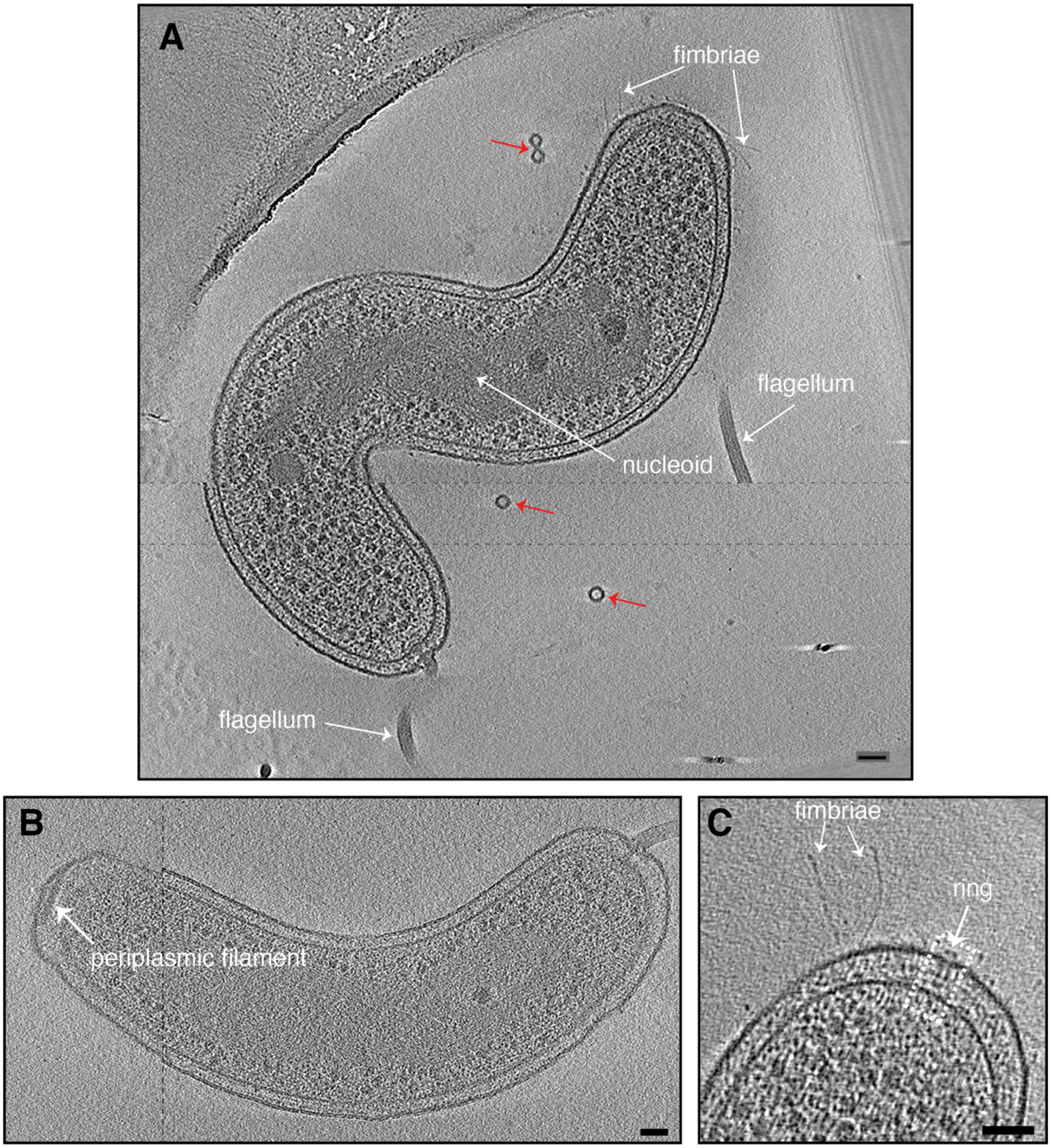


**Figure S3: A-C)** Slices through electron cryo-tomograms of *B. bacteriovorus* attack phase cells indicating the presence of uniformly sized vesicles (20-30 nm in diameter, red arrows) in the vicinity of the cell (A), periplasmic filamentous structure (B), and thin fimbriae (A, C). Dashed black lines in (A and B) indicate a composite of slices through the tomogram at different z-heights. Scale bar 50 nm.

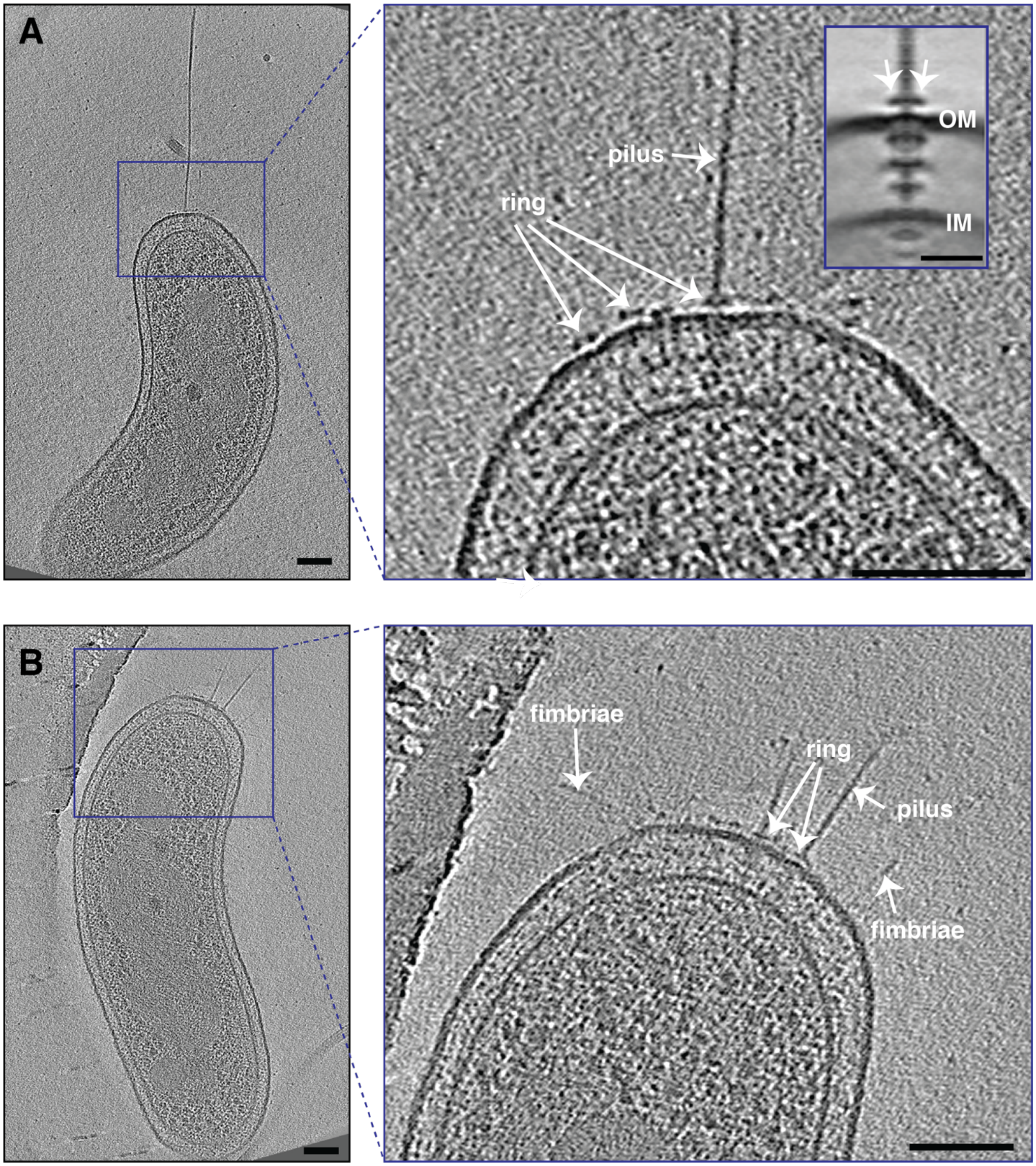


**Figure S4: A-B)** Slices through electron cryo-tomograms of *B. bacteriovorus* attack-phase cells highlighting piliated and non-piliated T4aP. Inset in the right panel of A is a subtomogram average of piliated T4aP basal bodies. Scale bars 100 nm, 20 nm in the subtomogram average inset. OM= outer membrane, IM= inner membrane.

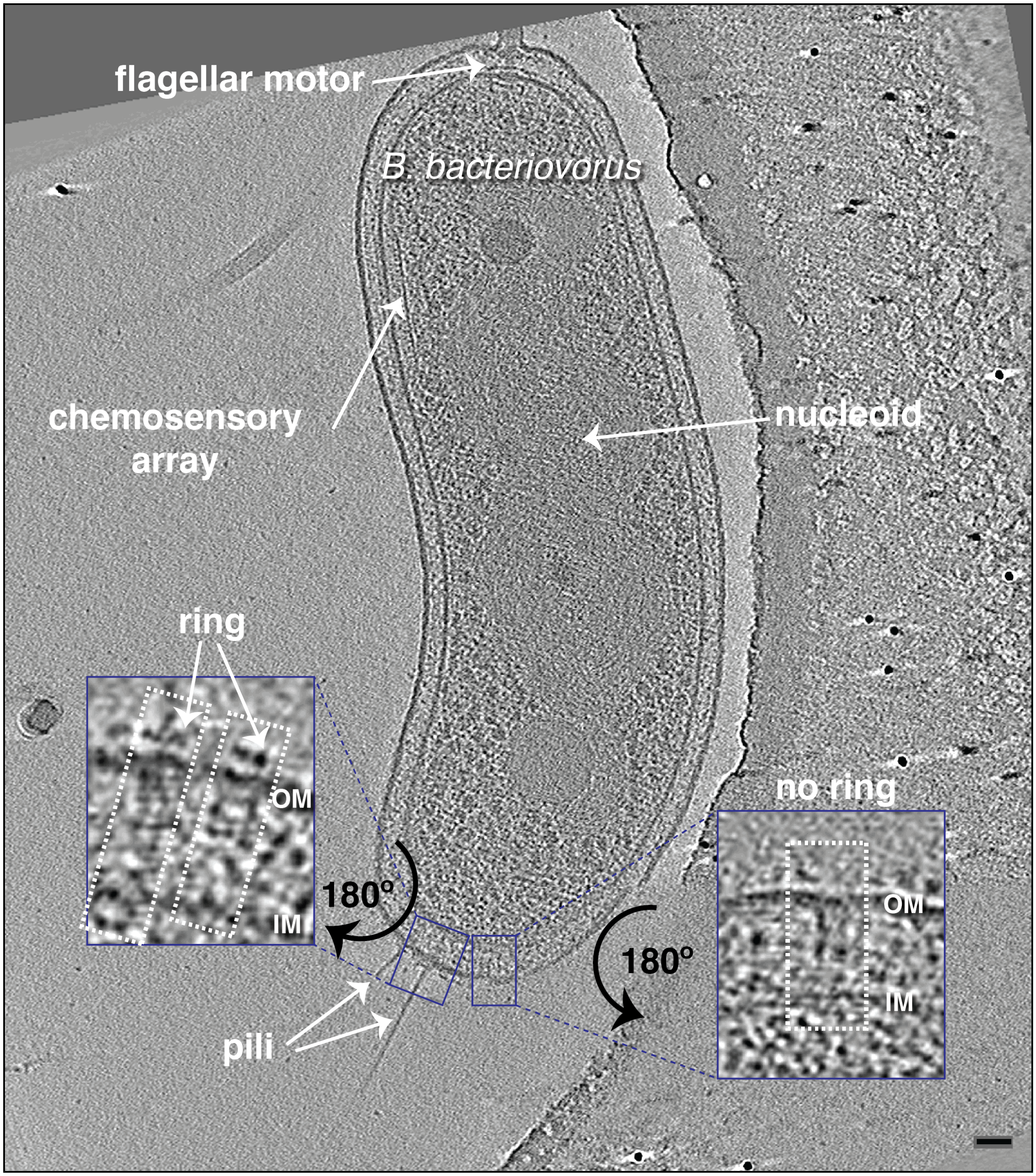


**Figure S5:** A slice through an electron cryo-tomogram of an attack phase *B. bacteriovorus* cell indicating the presence of piliated and non-piliated particles with (left enlargement) and without (right enlargement) extracellular ring. Scale bar 50 nm.

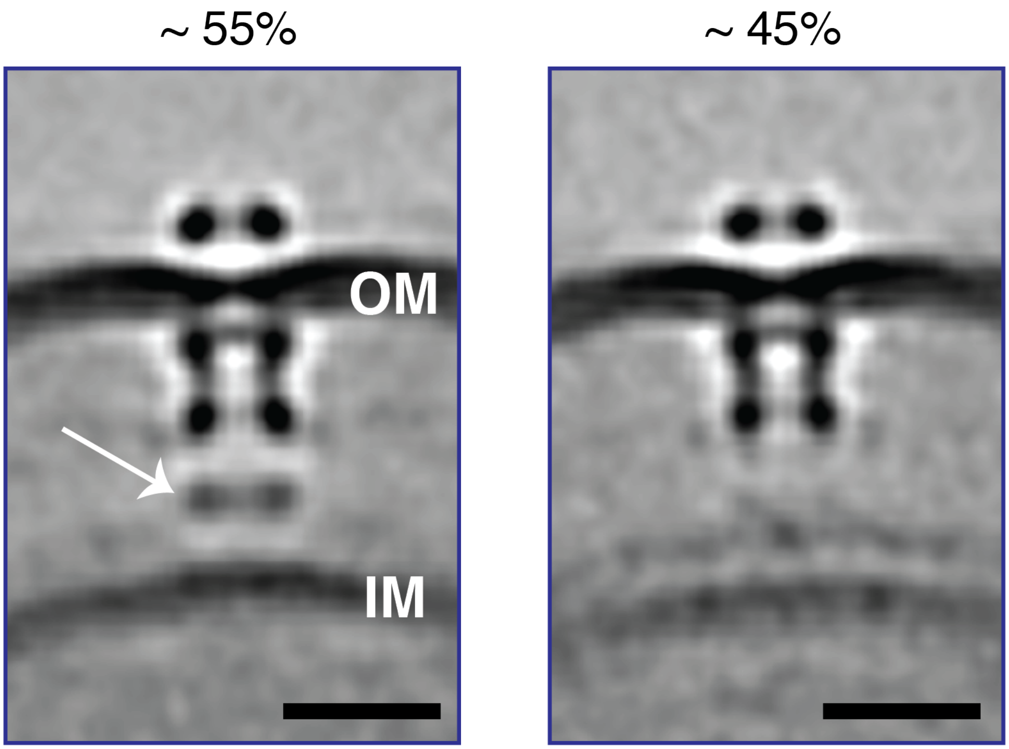


**Figure S6:** Central slices through subtomogram averages of non-piliated T4aP basal bodies with (left, white arrow) and without (right) the lower periplasmic ring. The percentage of each class is indicated above the subtomogram average. Scale bars 20 nm.

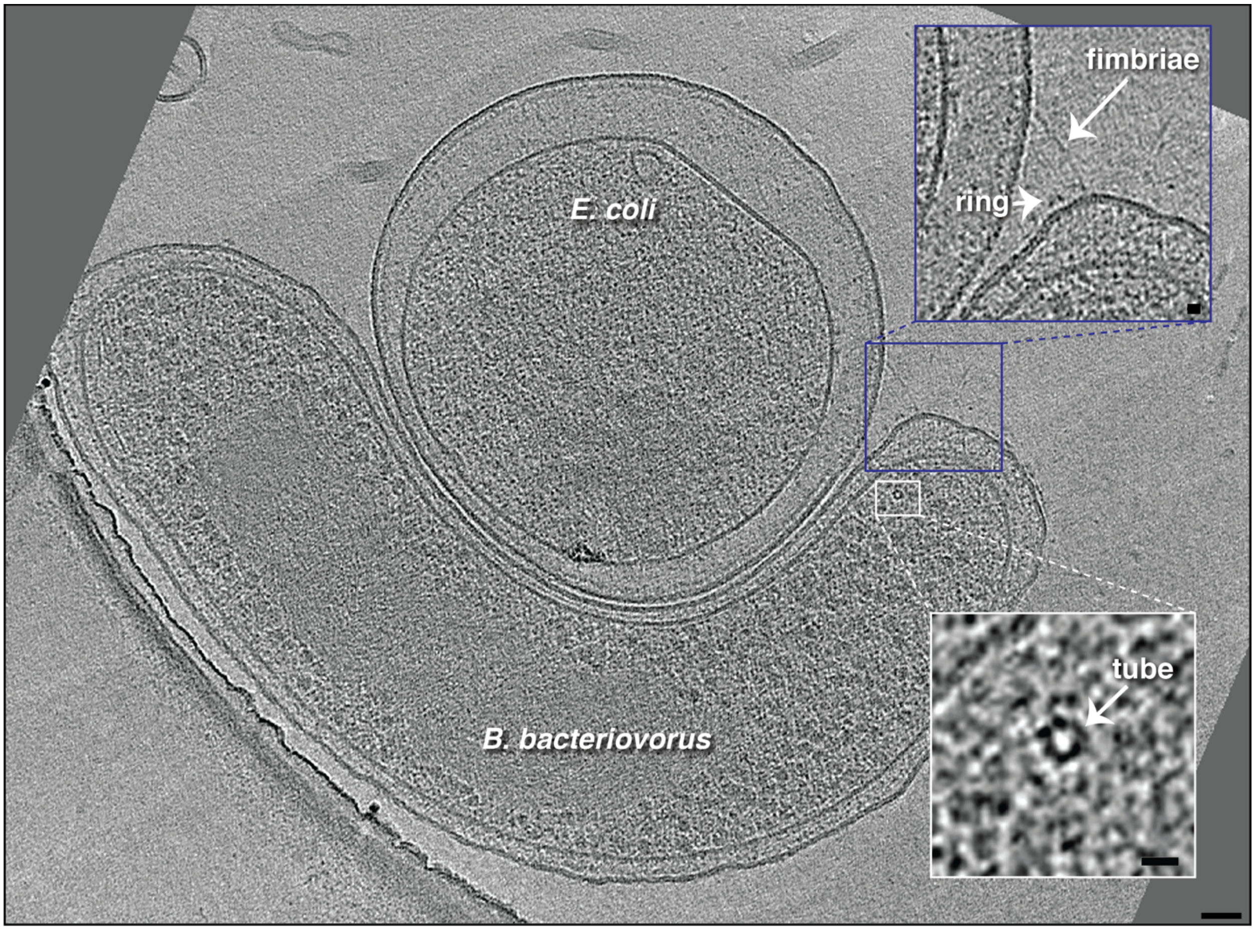


**Figure S7:** A slice through an electron cryo-tomogram of a *B. bacteriovorus* cell near an *E. coli* minicell. Upper, blue-boxed enlargement highlights a fimbria extending to the prey's outer membrane. An extracellular ring of a nearby non-piliated T4aP basal body is highlighted. Lower, white-boxed enlargement highlights a top view of an 8-nm wide cytoplasmic tube (white arrow). Scale bars 50 nm in the main panel, 10 nm in the enlargements.

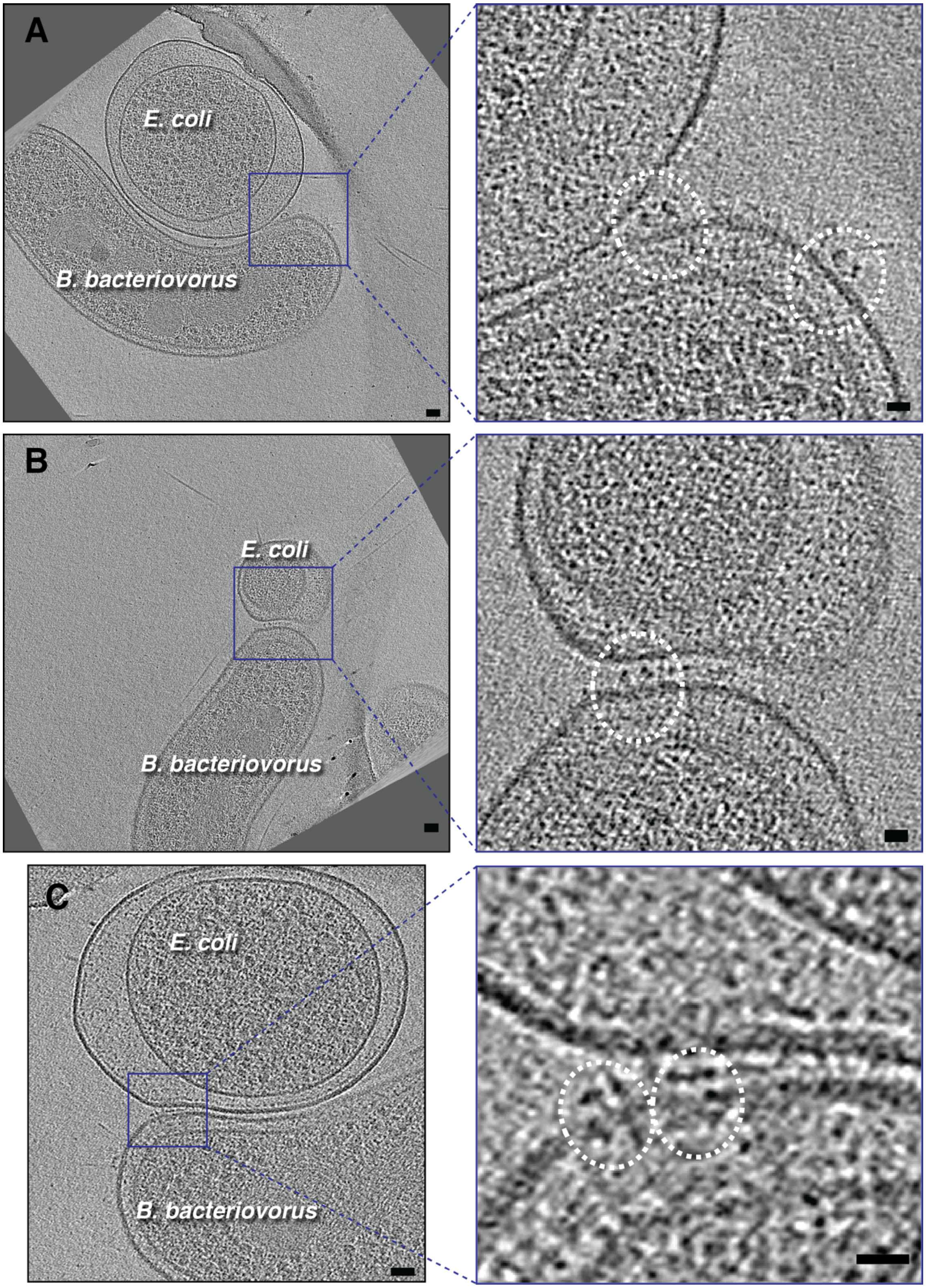
A
E. coli
B. bacteriovorus
B
E. coli
B. bacteriovorus
C
E. coli
B. bacteriovorus

**Figure S8: A-C)** Slices through electron cryo-tomograms of *B. bacteriovorus* near prey (*E. coli* minicells) highlighting rose-like complexes (A and B, white ellipses) and non-piliated T4aP basal bodies (C, white ellipses) in close contact with the prey outer membrane. Scale bars 50 nm in left panels, 20 nm in enlargements.

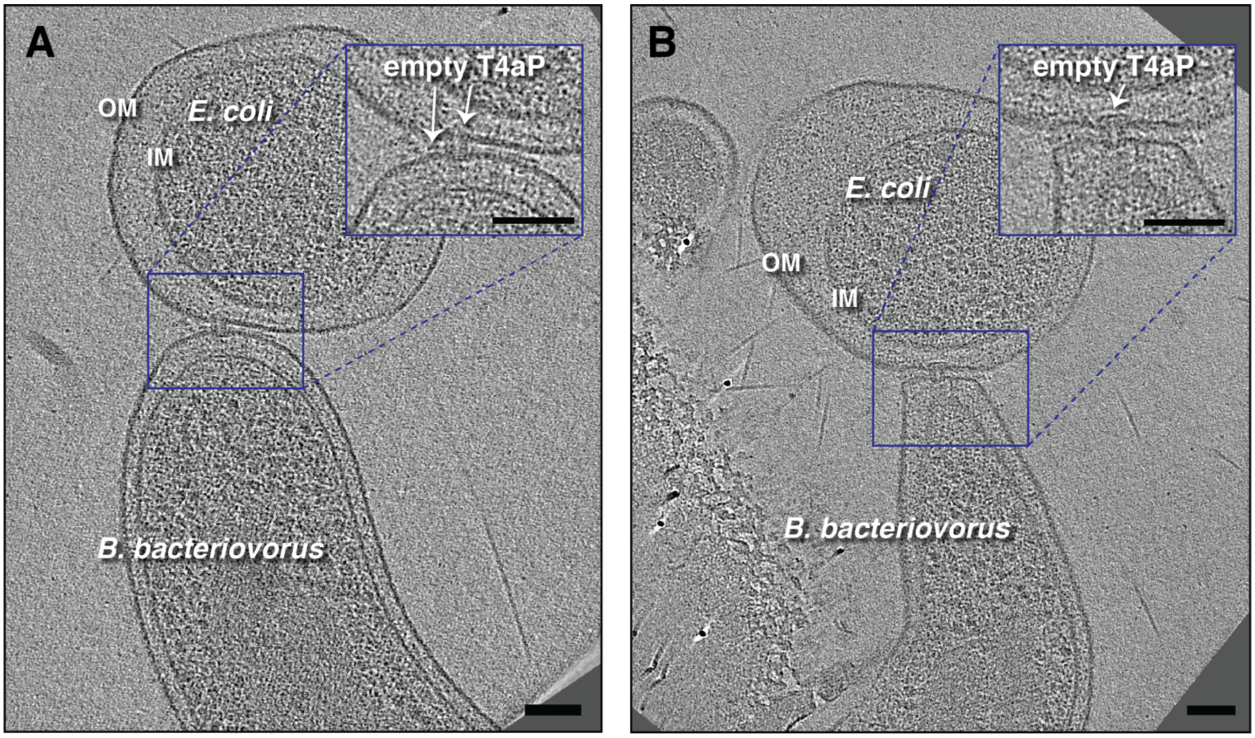


**Figure S9: A, B)** Slices through electron cryo-tomograms of *B. bacteriovorus* attached to prey (*E. coli* minicells) showing non-piliated T4aP basal bodies (white arrows) penetrating to the prey's PG layer. Scale bars 100 nm.

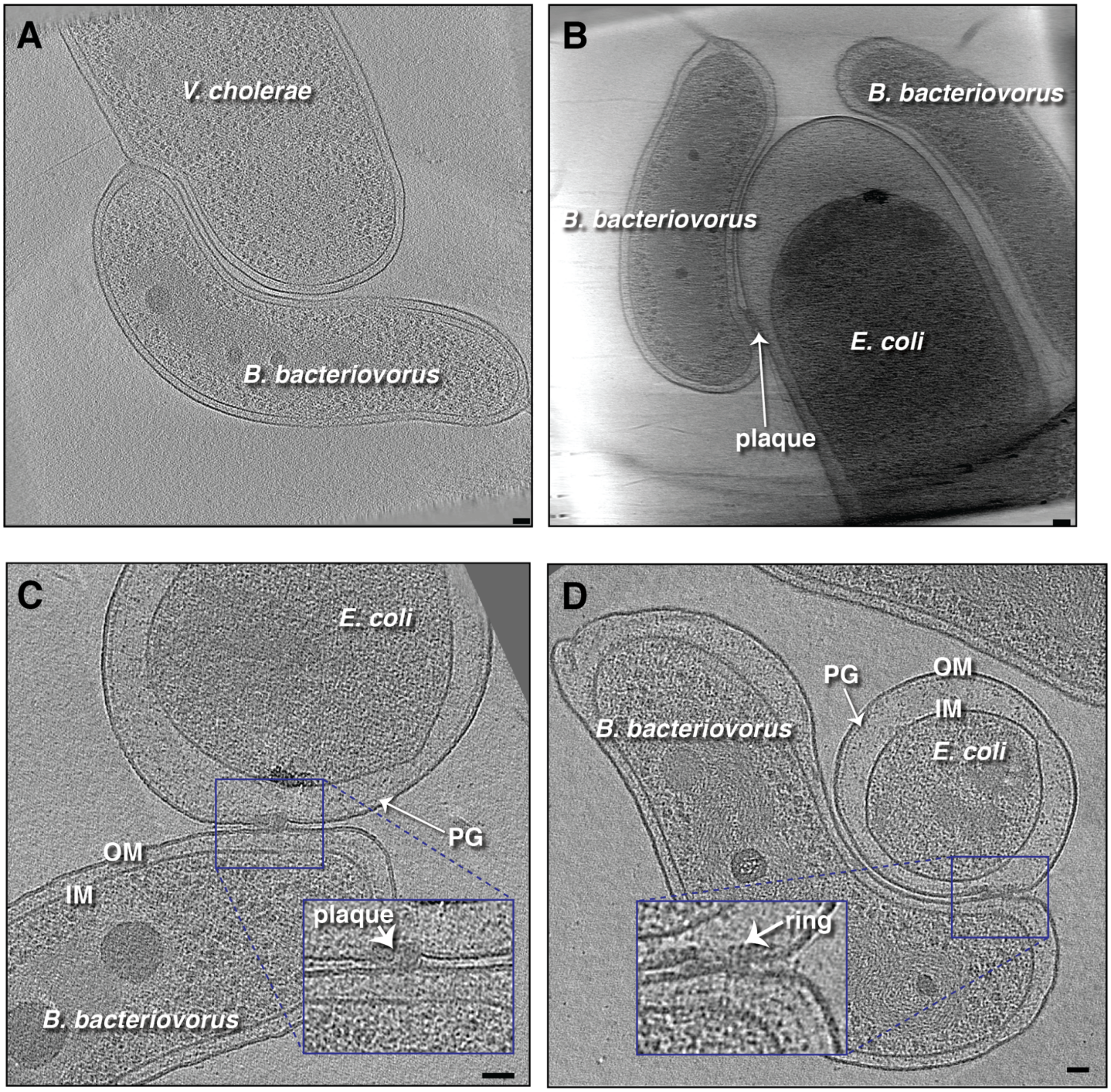


**Figure S10: A, B)** Slices through electron cryo-tomograms showing *B. bacteriovorus* cells attached to full-sized prey (*V. cholerae* in (A) and *E. coli* in (A)). An attachment plaque is visible in B (white arrow). Scale bars 50 nm. **C, D)** Slices through electron cryo-tomograms of *B. bacteriovorus* attached to prey (*E. coli* minicell) with a lateral attachment plaque. PG = peptidoglycan, OM = outer membrane, IM = inner membrane. Scale bars 50 nm.

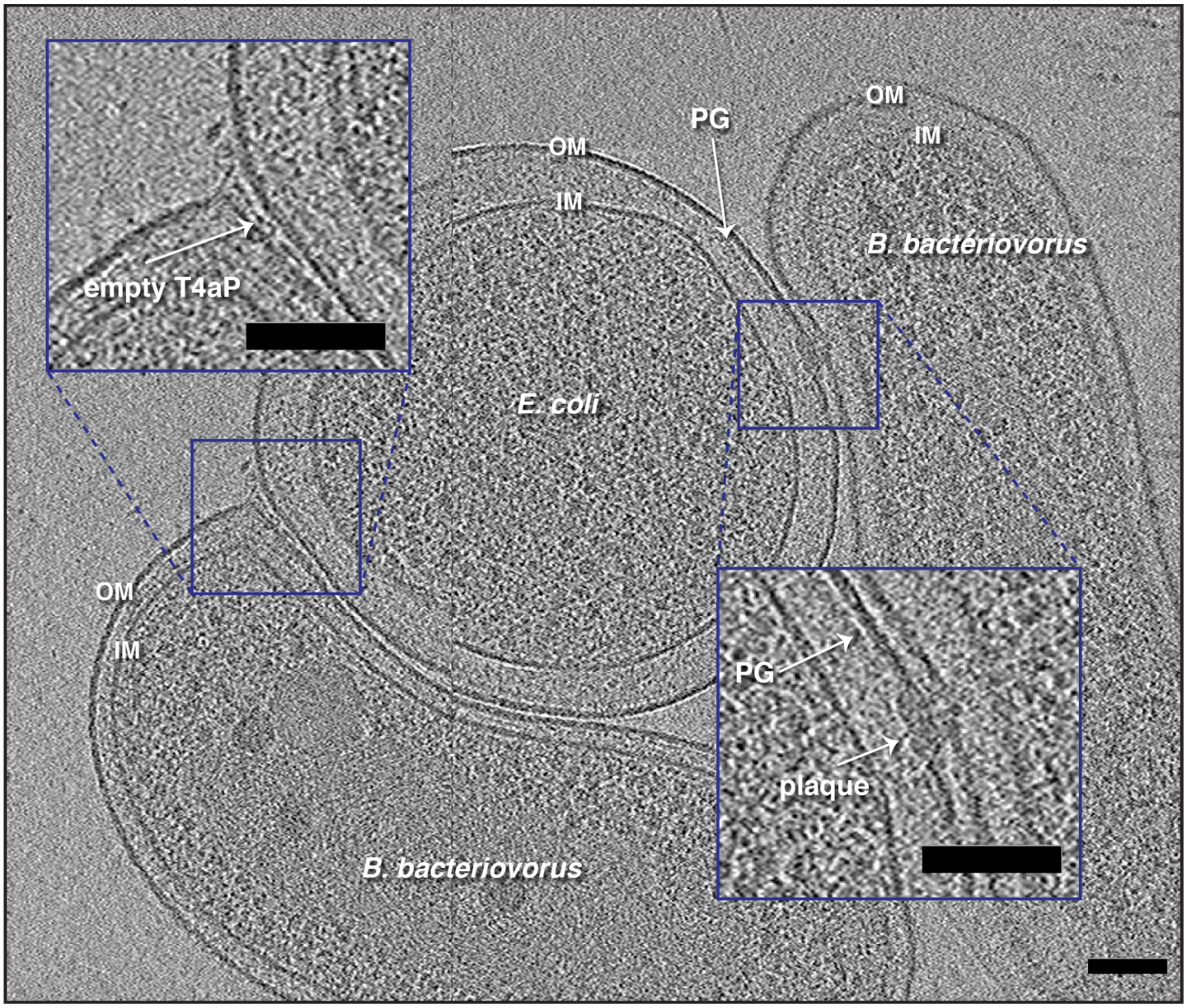


**Figure S11:** A slice through an electron cryo-tomogram showing the presence of two predators attached to a prey (*E. coli* minicell). One of the predators is attached via T4aP (left), while the other via an attachment plaque (right). PG = peptidoglycan, OM = outer membrane, IM = inner membrane. Scale bar is 100 nm.

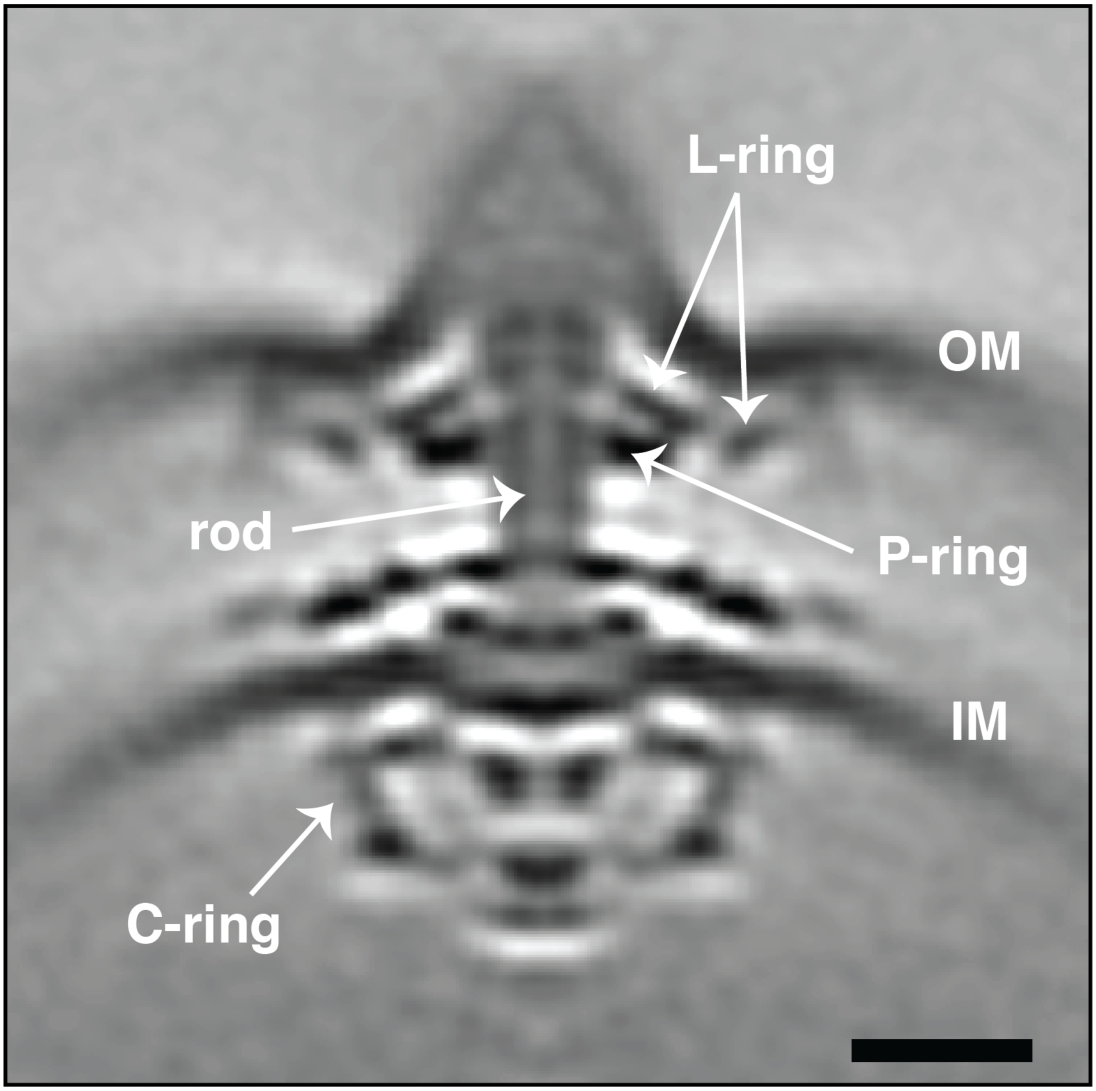


**Figure S12:** A central slice through the subtomogram average of the *B. bacteriovorus* flagellar motor with major parts annotated. Scale bar is 20 nm. OM = outer membrane, IM = inner membrane.

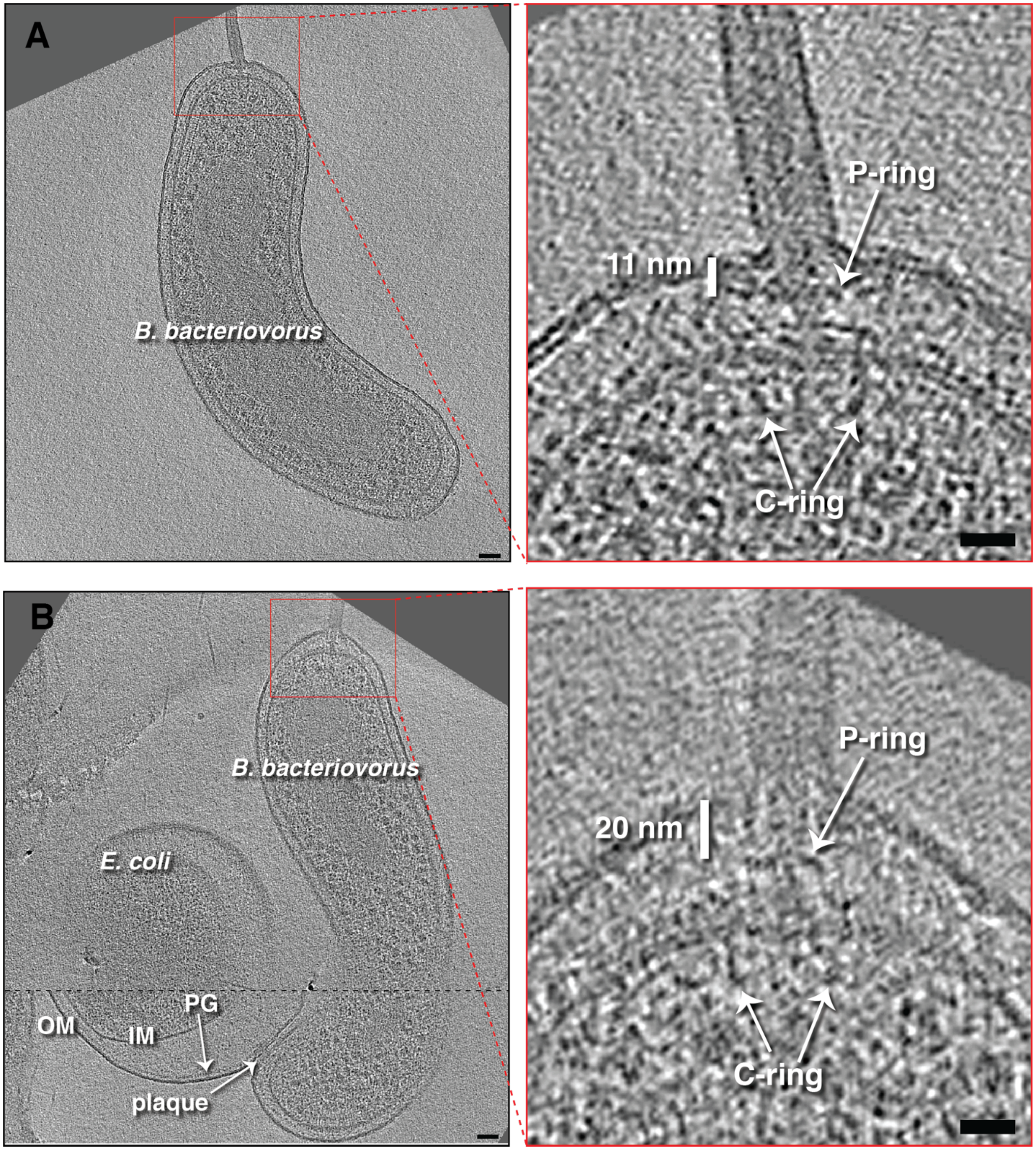


**Figure S13:** A slice through an electron cryo-tomogram (left) and enlarged view (right) of an attack-phase *B. bacteriovorus* cell illustrating the flagellar motor at the flagellated pole. The distance between the flagellar P-ring and the outer membrane is indicated (~11 nm). **B)** A slice through an electron cryo-tomogram (left) and enlarged view (right) of a *B. bacteriovorus* cell attached to a prey with an attachment plaque. Dashed black line indicates a composite of slices through the tomogram at different z-heights. The distance between the flagellar P-ring and the outer membrane is indicated (~20 nm), highlighting an early stage of flagellar resorption. Scale bars 50 nm in left panels, 20 nm in enlargements.

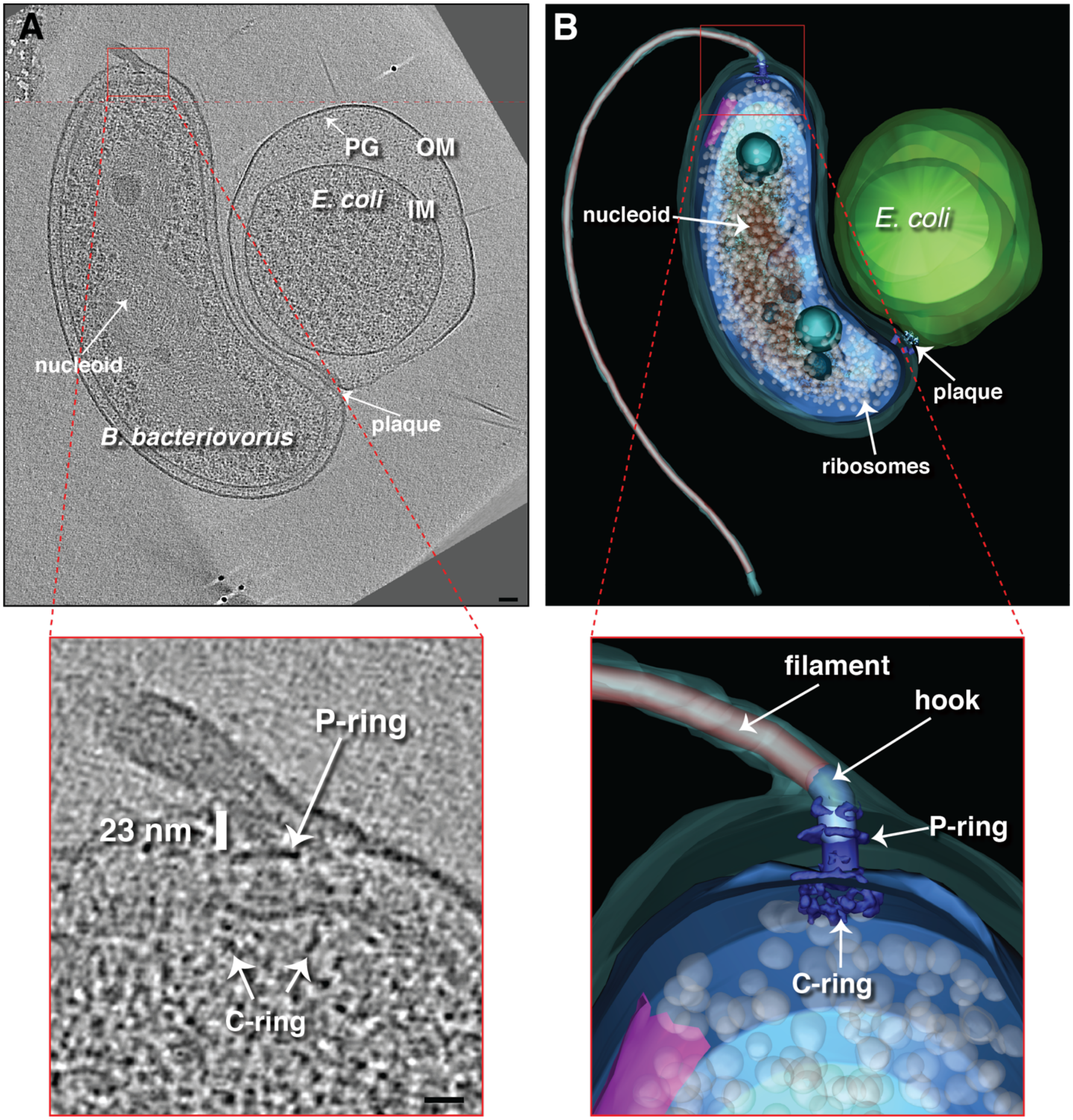


**Figure S14: A)** A slice through an electron cryo-tomogram (top) and enlarged view (bottom) of a *B. bacteriovorus* cell attached to a prey with an attachment plaque at an early stage of flagellar resorption. The distance between the flagellar P-ring and the outer membrane is indicated (~23 nm). **B)** A 3D segmentation of the cryo-tomogram shown in (A) with an enlargement highlighting the flagellar motor. Scale bars 50 nm in the top panel and 20 nm in the enlargement.

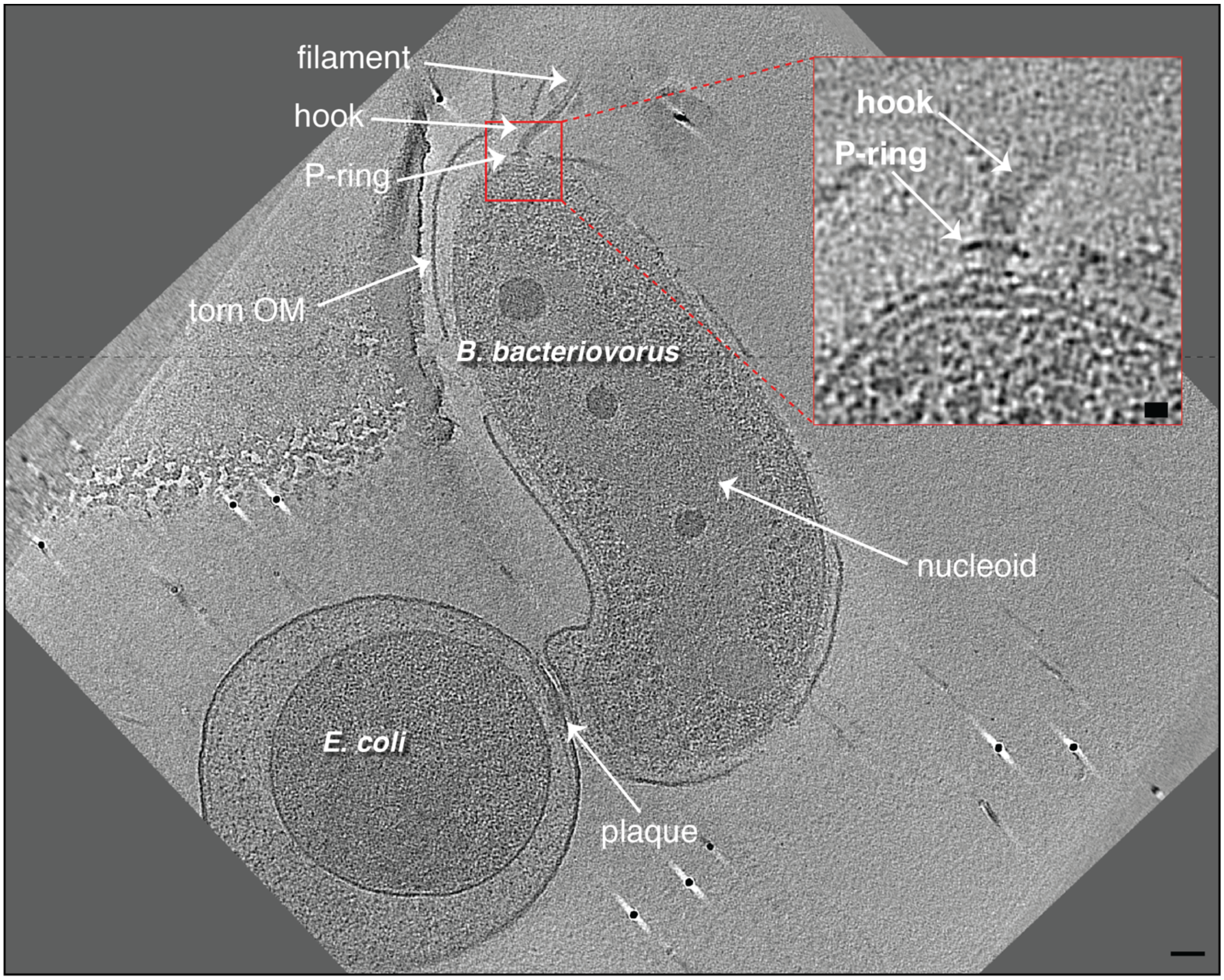


**Figure S15:** A Slice through an electron cryo-tomogram and enlarged view (inset) of a *B. bacteriovorus* cell attached to *E. coli* minicell. Note the torn outer membrane and presence of the P-ring but not the L-ring in the flagellar motor. Dashed black line indicates a composite of slices through the tomogram at different z-heights. Scale bar is 50 nm in the main panel, 10 nm in the inset.

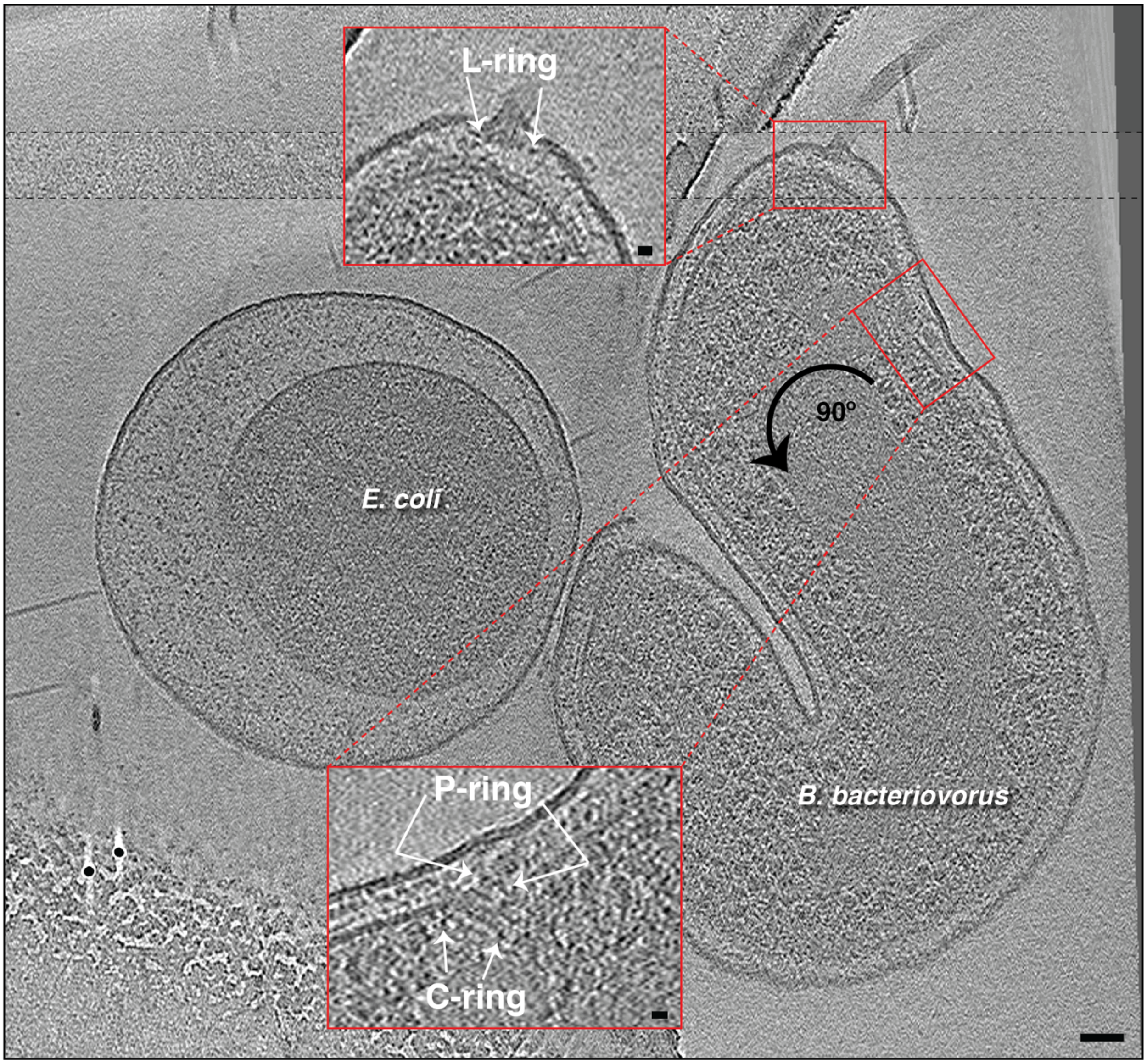


**Figure S16:** A slice through an electron cryo-tomogram of a *B. bacteriovorus* cell attached to *E. coli* minicell illustrating the absorption of the predator's flagellum. Upper enlargement highlights the flagellar L-ring in the outer membrane. Lower enlargement shows the internalized flagellar motor lacking the L-ring. Dashed black lines indicate a composite of slices through the tomogram at different z-heights. Scale bar is 50 nm in the main panel, 10 nm in the insets.

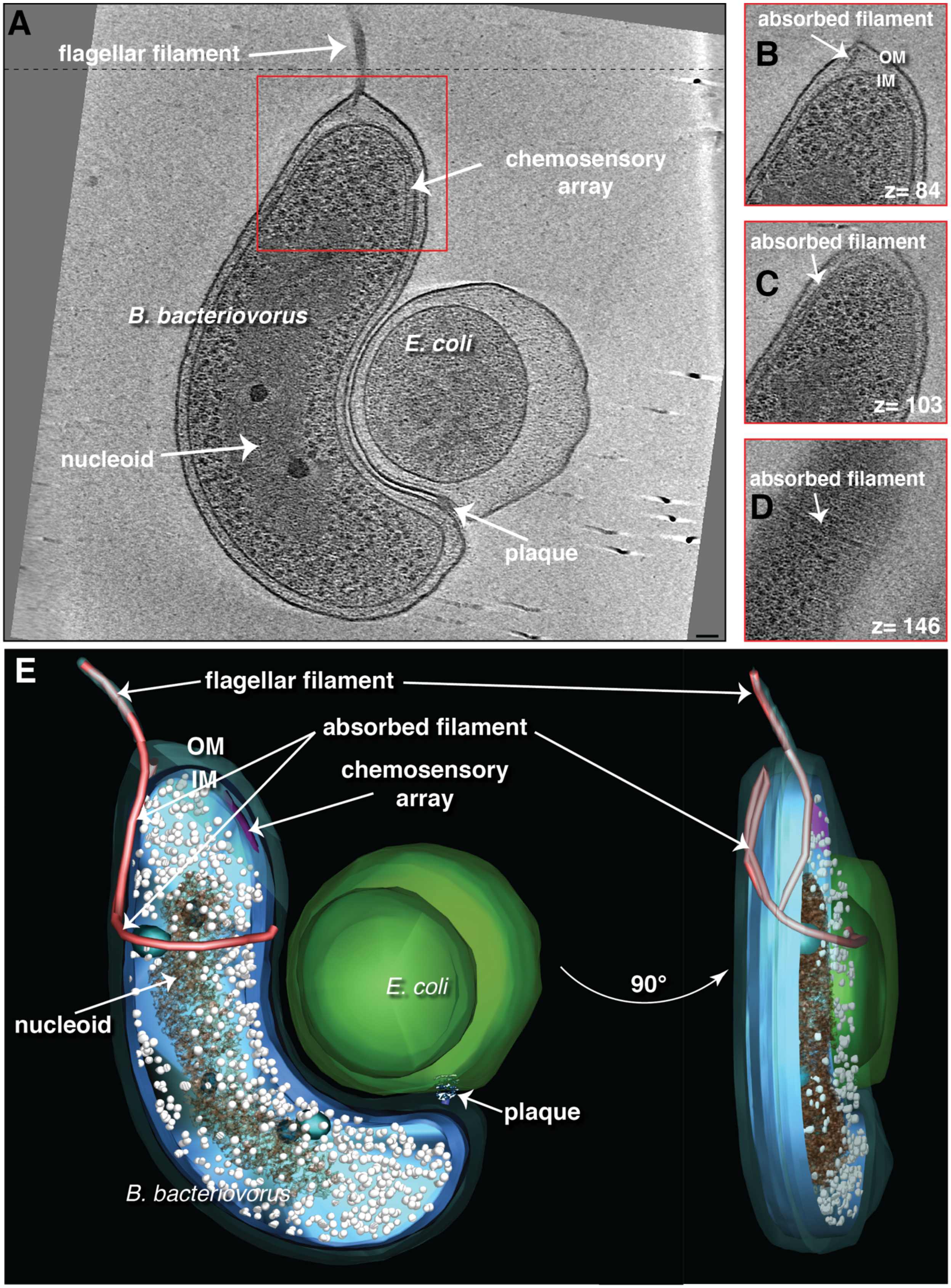

A
flagellar filament
chemosensory array
B. bacteriovorus
E. coli
nucleoid
plaque
absorbed filament
B
OM
IM
z= 84
absorbed filament
C
z= 103
absorbed filament
D
z= 146
E
flagellar filament
absorbed filament
OM
IM
chemosensory array
E. coli
nucleoid
plaque
B. bacteriovorus
90°

**Figure S17: A)** A slice through an electron cryo-tomogram of a *B. bacteriovorus* cell attached to an *E. coli* minicell with an attachment plaque illustrating flagellar resorption. Scale bar is 50 nm. **B-D)** Slices at different z-heights of the red-boxed area in the cryo-tomogram in (A) showing the absorbed flagellar filament wrapped around the cell. **E)** Rotated views of a 3D segmentation of the tomogram shown in (A) highlighting the retracted periplasmic flagellar filament.

331

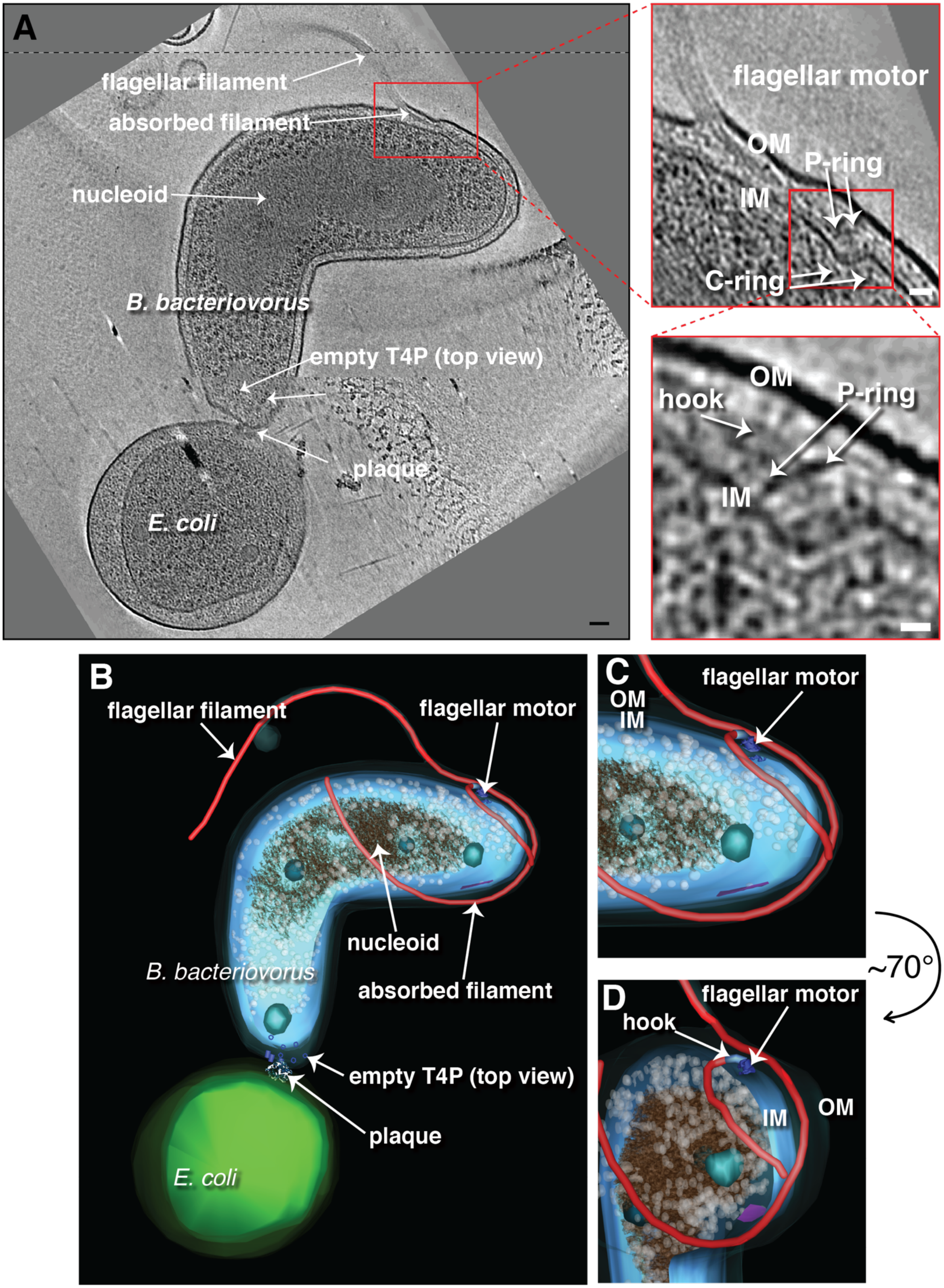
A
flagellar filament
absorbed filament
nucleoid
B. bacteriovorus
empty T4P (top view)
plaque
E. coli
flagellar motor
OM
P-ring
IM
C-ring
OM
hook
P-ring
IM
B
flagellar filament
flagellar motor
nucleoid
B. bacteriovorus
absorbed filament
empty T4P (top view)
plaque
E. coli
C
OM
IM
flagellar motor
~70°
D
flagellar motor
hook
IM
OM

**Figure S18: A)** A slice through an electron cryo-tomogram of a *B. bacteriovorus* cell attached to a prey with an attachment plaque at a late stage of flagellar resorption. The flagellar entry hole is shifted to the side of the predator cell. Dashed black line indicates a composite of slices through the tomogram at different z-heights. Successive enlargements on the right highlight the features of the absorbed flagellar motor. Scale bars 50 nm in left panel, 20 nm in top enlargement, 10 nm in bottom enlargement. **B-D**) Various views of a 3D segmentation of the cryo-tomogram shown in (A) highlighting the absorbed flagellum (including the motor).

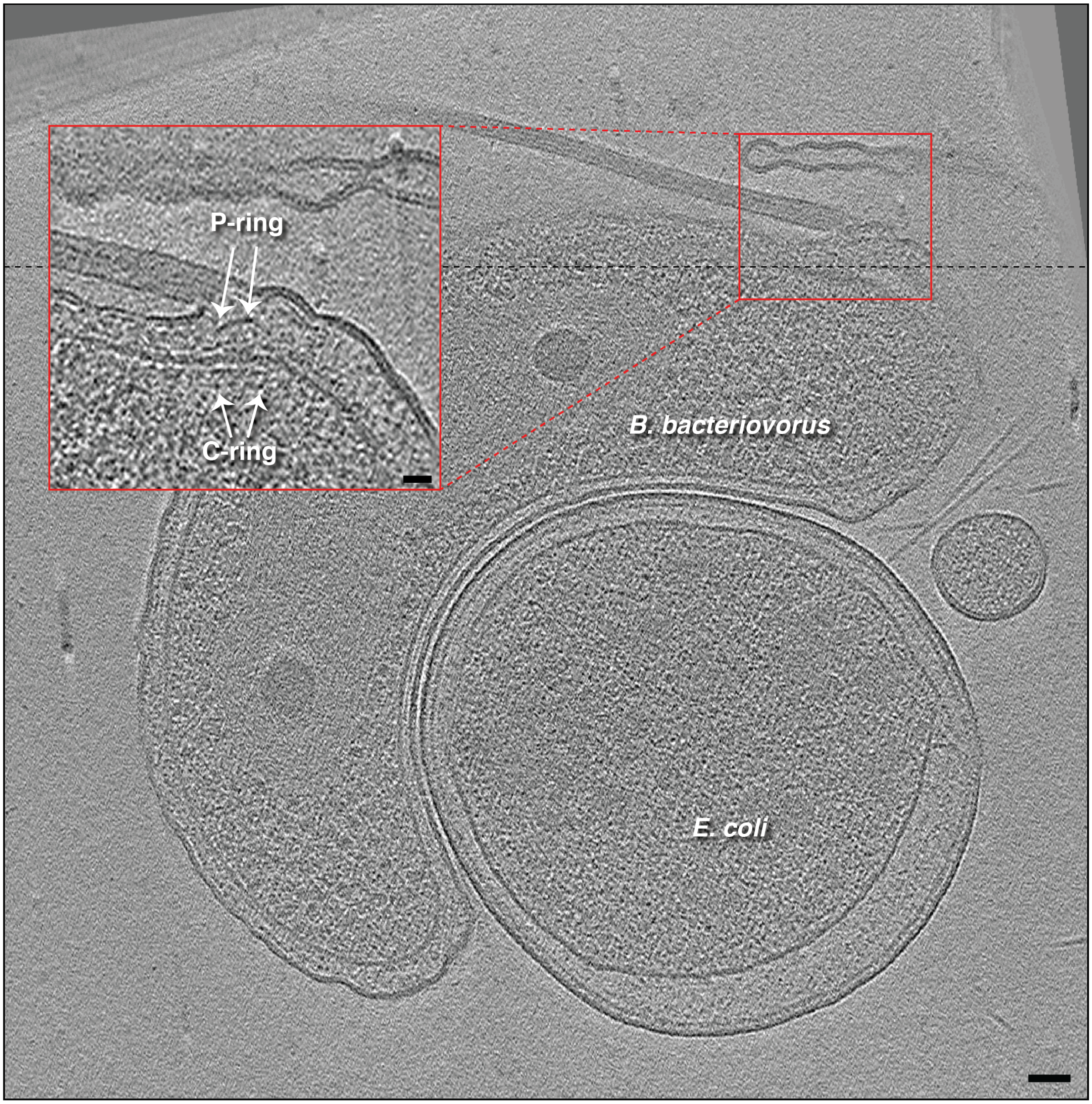


**Figure S19:** A slice through an electron cryo-tomogram of a *B. bacteriovorus* cell attached to an *E. coli* minicell illustrating the absorption of the predator's flagellum and the shift of the flagellar exit hole to the side of the predator's cell. Dashed black line indicates a composite of slices through the tomogram at different z-heights. Enlargement shows the internalized flagellar motor with the P-ring and cytoplasmic C-ring labeled. Scale bar 50 nm in the main panel, 20 nm in the inset.

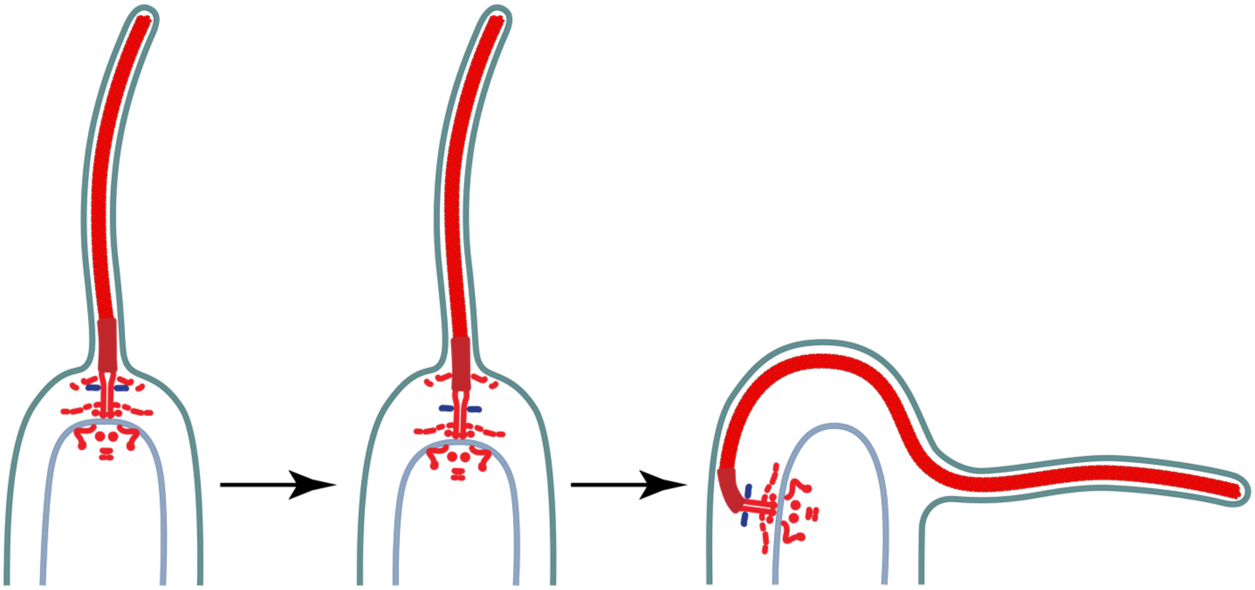

**Figure S20:** A schematic representation of the process of flagellar retraction that occurs during *B. bacteriovorus* prey invasion based on our cryo-ET imaging. The P-ring is highlighted in blue.

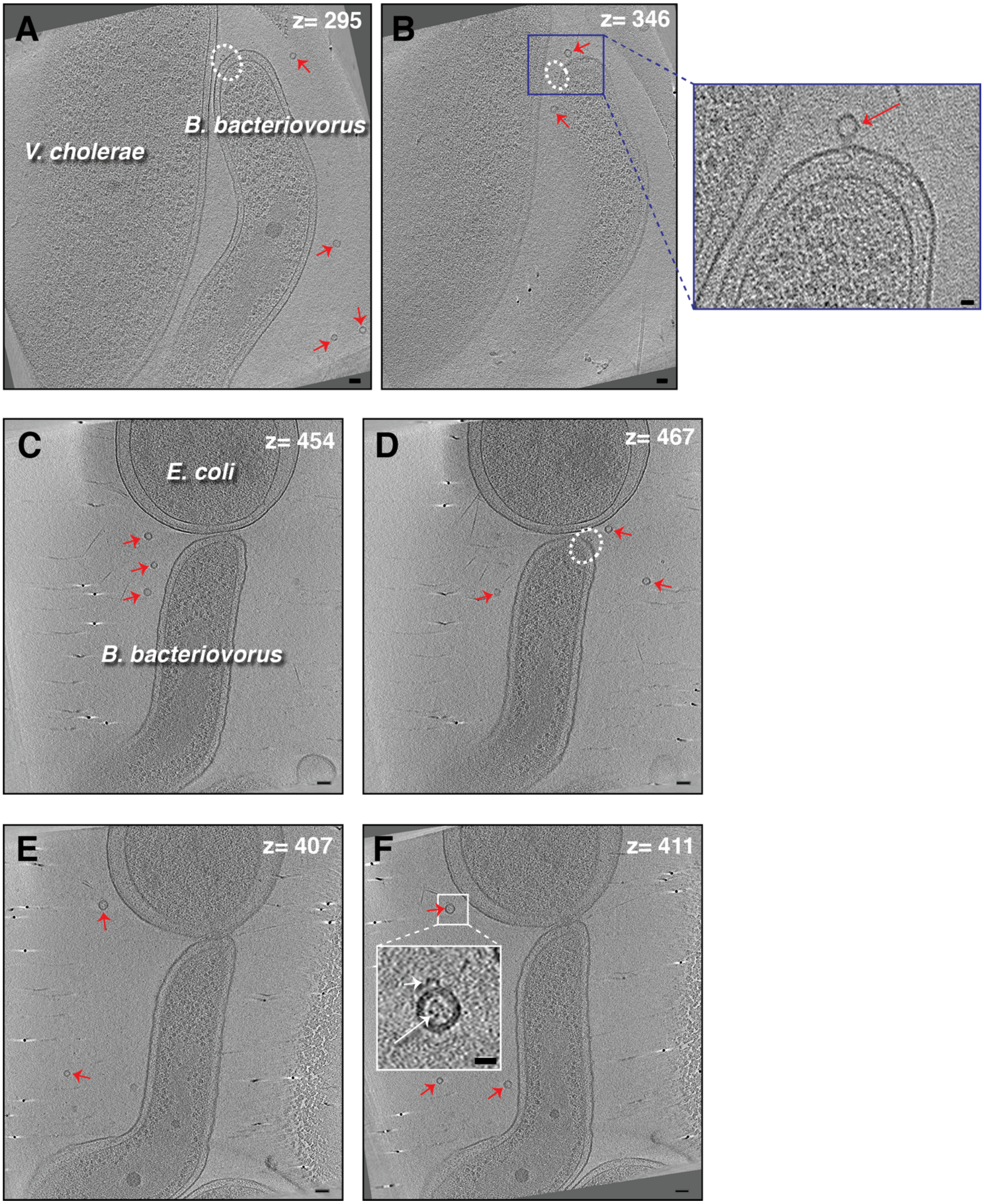


**Figure S21: A, B)** Slices at different z-heights through an electron cryo-tomogram of *B. bacteriovorus* near a *V. cholerae* cell highlighting multiple uniformly-sized vesicles (red arrows)

in the vicinity of the cells or budding from the *B. bacteriovorus* pole (shown in the enlargement in panel B). Dashed ellipses indicate a rose-like complex. **C-F)** slices at different z-heights through an electron cryo-tomogram of *B. bacteriovorus* attached to an *E. coli* minicell highlighting multiple 20-30 nm-diameter vesicles near the cells. The enlargement in (F) shows dark densities inside and outside the vesicle (white arrows). Scale bars 50 nm in main panels of (A-F), 20 nm in enlargements.

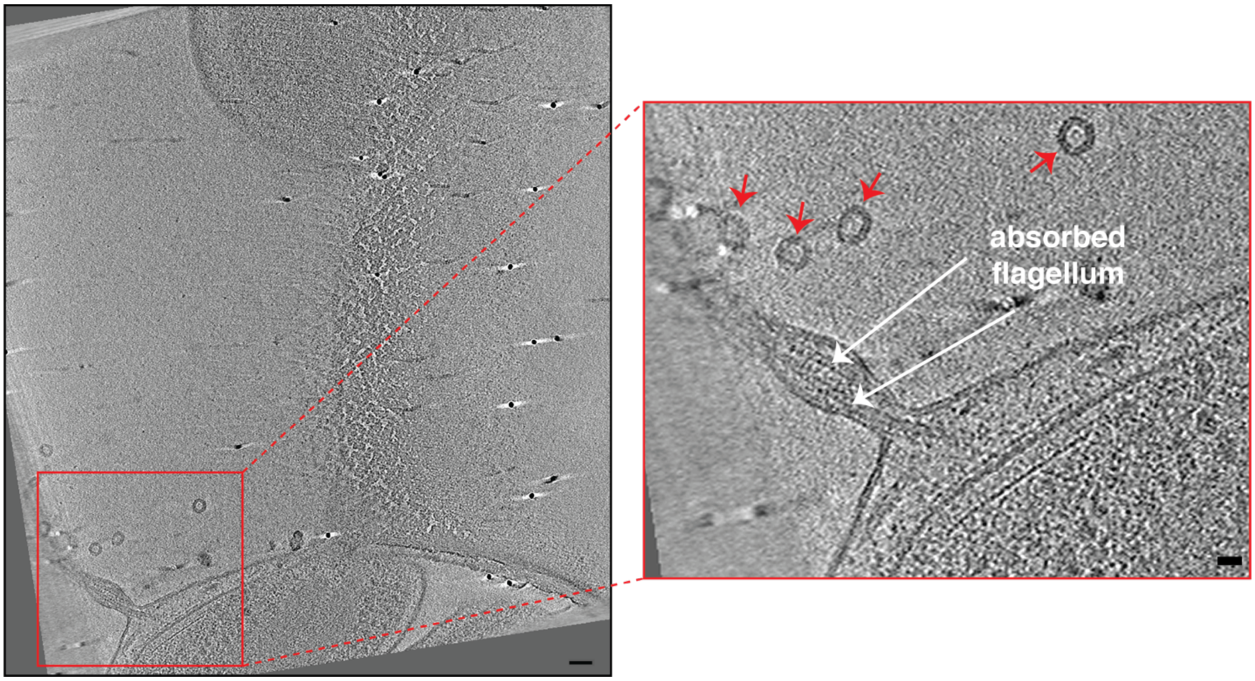


**Figure S22:** A slice through an electron cryo-tomogram of *B. bacteriovorus* attached to an *E. coli* minicell (different z-slices of the same tomogram are shown in Figure S21 C-F) indicating multiple homogenously-sized vesicles in the vicinity of the resorbing sheathed flagellum of the predator. Scale bars 50 nm in left panel, 20 nm in the enlargement.

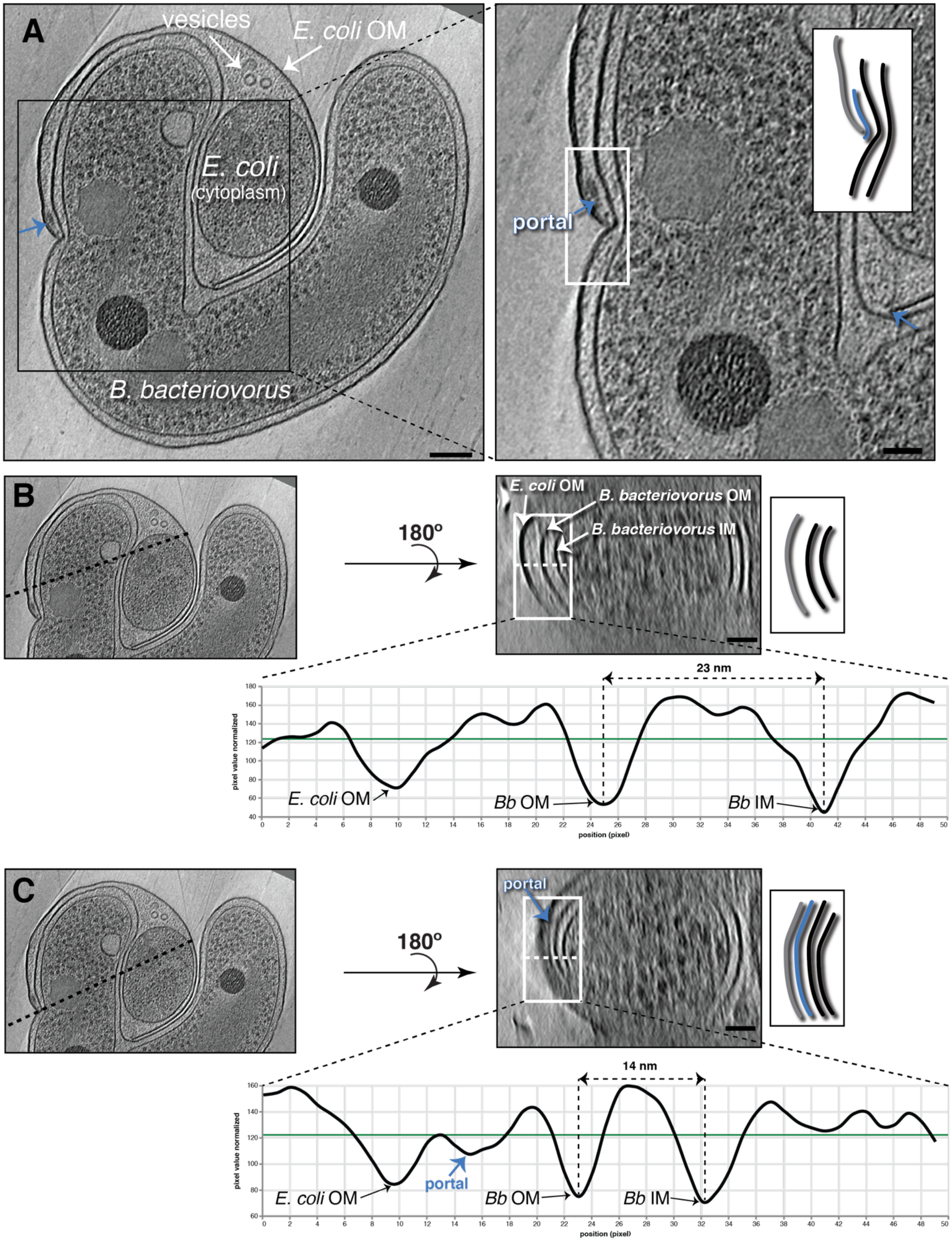
A
vesicles
E. coli OM
E. coli
(cytoplasm)
B. bacteriovorus
portal
B
180º
E. coli OM
B. bacteriovorus OM
B. bacteriovorus IM
23 nm
pixel value normalized
position (pixel)
E. coli OM
Bb OM
Bb IM
C
180º
portal
14 nm
pixel value normalized
position (pixel)
E. coli OM
portal
Bb OM
Bb IM

**Figure S23: A)** A slice through an electron cryo-tomogram (left) and enlarged view (right) showing a stalled invasion by a *B. bacteriovorus* of an *E. coli* minicell. Blue arrows and inset schematic highlight the portal. **B)** Right (top): Cross-section through the *yz* plane of the cryo-tomogram shown in (A) along the black dotted line indicated on the left. Note that this slice does not include the portal. Right (bottom): Average density profile taken along the white dashed line (inside the white rectangle) in the top panel. The distance between the predator's inner (IM) and outer membranes (OM) is indicated (~23 nm). **C)** Similar to (B) but for a *yz* slice where the portal is visible. The distance between the predator's inner and outer membranes is indicated (~14 nm). The schematics in the right panels of (A-C) represent the white-boxed areas in the corresponding slices, with the portal shown in blue, the prey outer membrane in grey, and the predator inner and outer membranes in black. Scale bars 100 nm in the left panel of (A), and 50 nm in other panels.

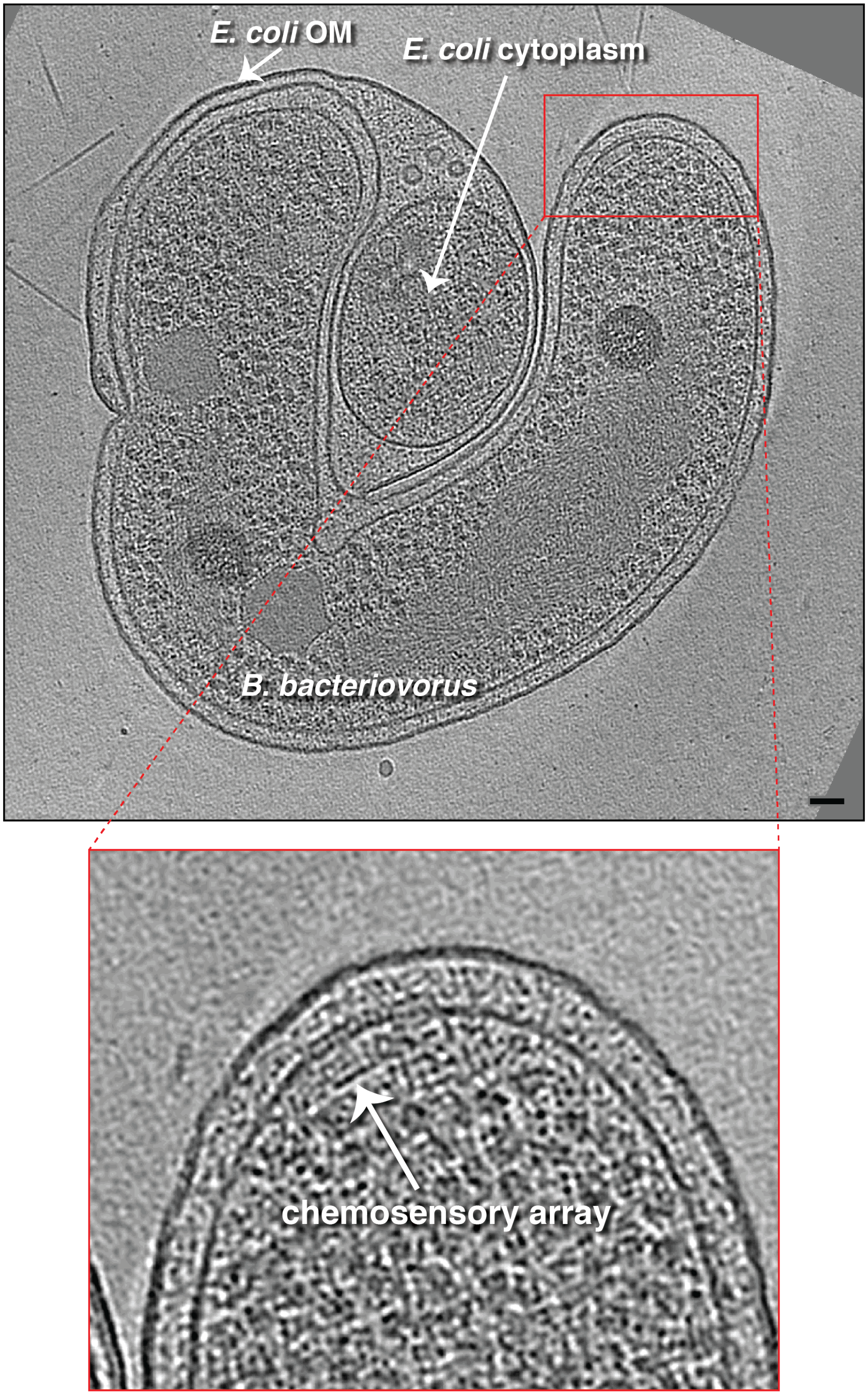
E. coli OM
E. coli cytoplasm
B. bacteriovorus
chemosensory array

**Figure S24:** A slice through an electron cryo-tomogram (top) and enlarged view (bottom) of a stalled invasion by a *B. bacteriovorus* cell of an *E. coli* minicell, highlighting a short chemosensory array (white arrow). Scale bar 50 nm.

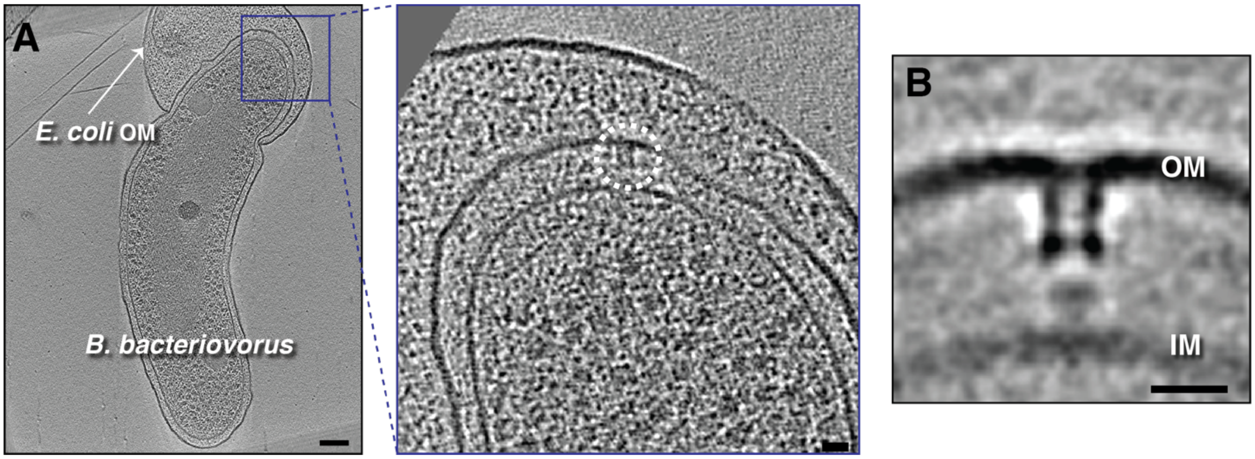


**Figure S25: A)** A slice through an electron cryo-tomogram (left) and enlarged view (right) of a stalled invasion by a *B. bacteriovorus* cell of an *E. coli* minicell highlighting a secretin-containing particle (putative T4aP) lacking the extracellular ring (white circle). **B)** A central slice through the subtomogram average of putative T4aP particles without extracellular rings found on predator biting poles inside prey. OM = outer membrane, IM = inner membrane. Scale bars 100 nm in left panel of (A), and 20 nm in other panels.

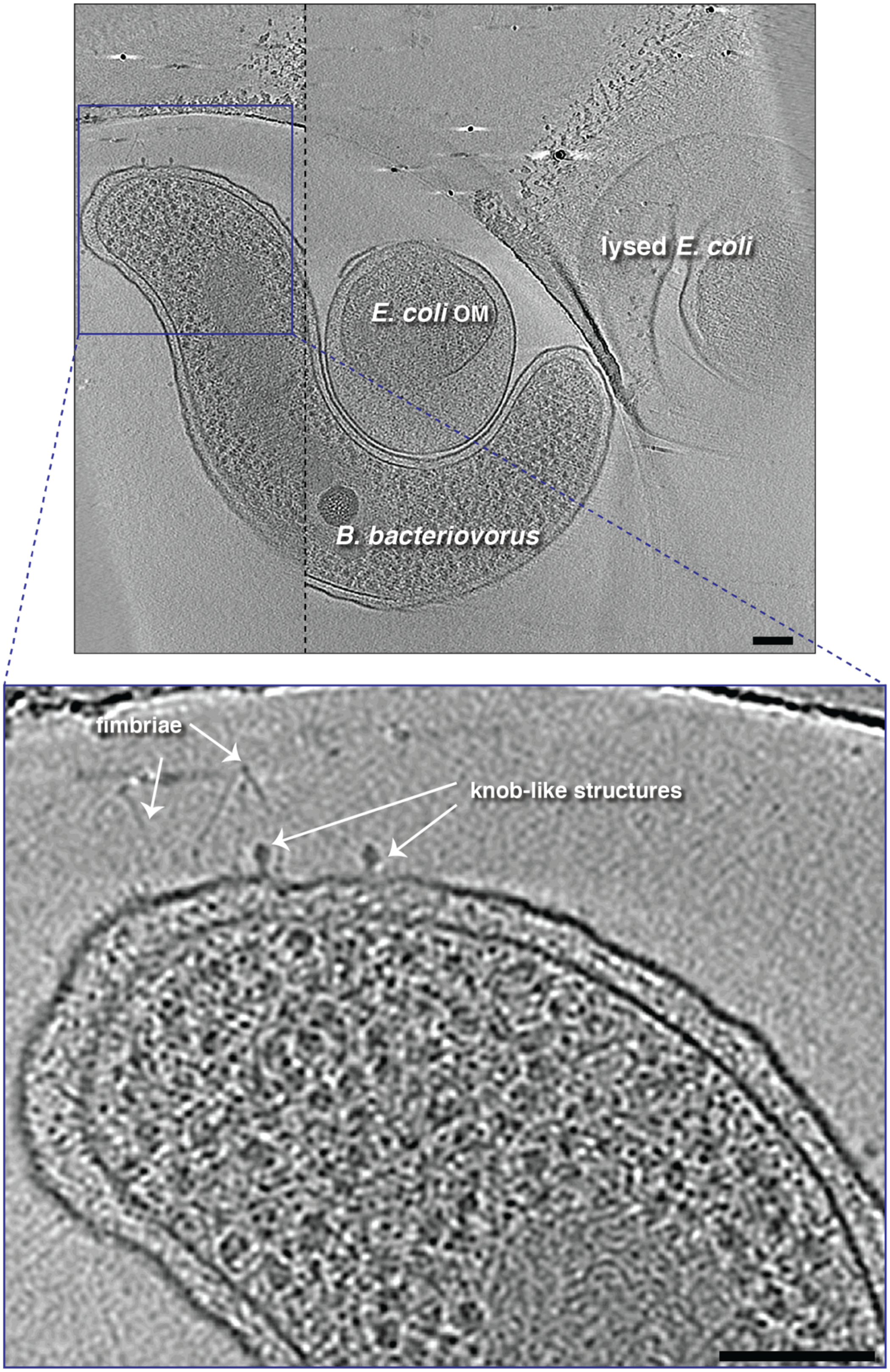
lysed E. coli
E. coli OM
B. bacteriovorus
fimbriae
knob-like structures

**Figure S26:** A slice through an electron cryo-tomogram (top) and enlarged view (bottom) showing a *B. bacteriovorus* cell near a lysed cell, highlighting knob-like densities. Dashed black line indicates a composite of slices through the tomogram at different z-heights. Scale bars 100 nm.

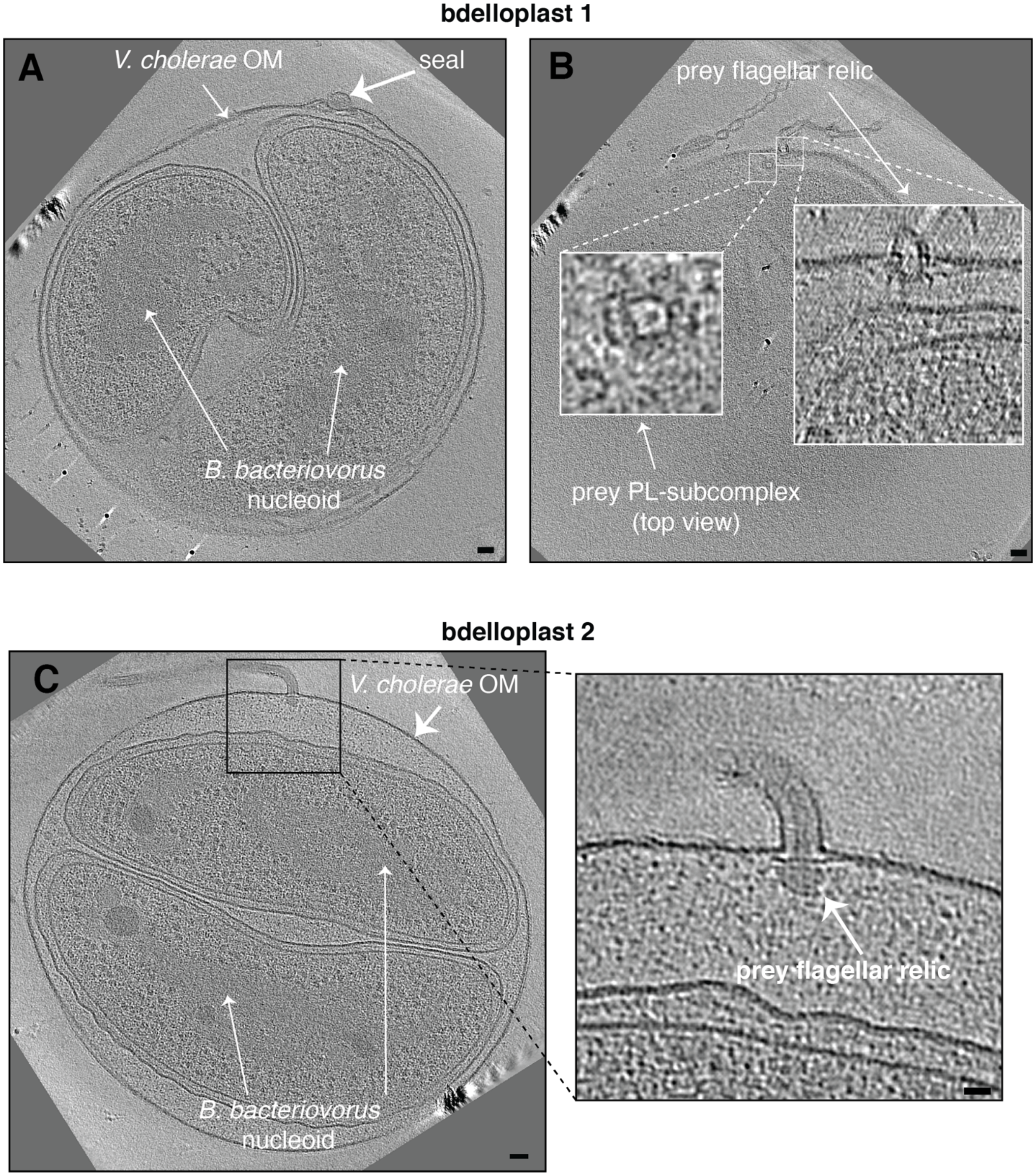


**Figure S27: A, B)** Slices at different z-heights through an electron cryo-tomogram and enlargements of a *V. cholerae* bdelloplast containing two *B. bacteriovorus* cells highlighting the prey flagellar relic and a top view of a PL-subcomplex. **C)** A slice through an electron cryo-tomogram (left) and enlargement (right) of a different bdelloplast highlighting a prey flagellar relic attached to the bdelloplast. Scale bars 50 nm in the main panels, 20 nm in the enlargement.

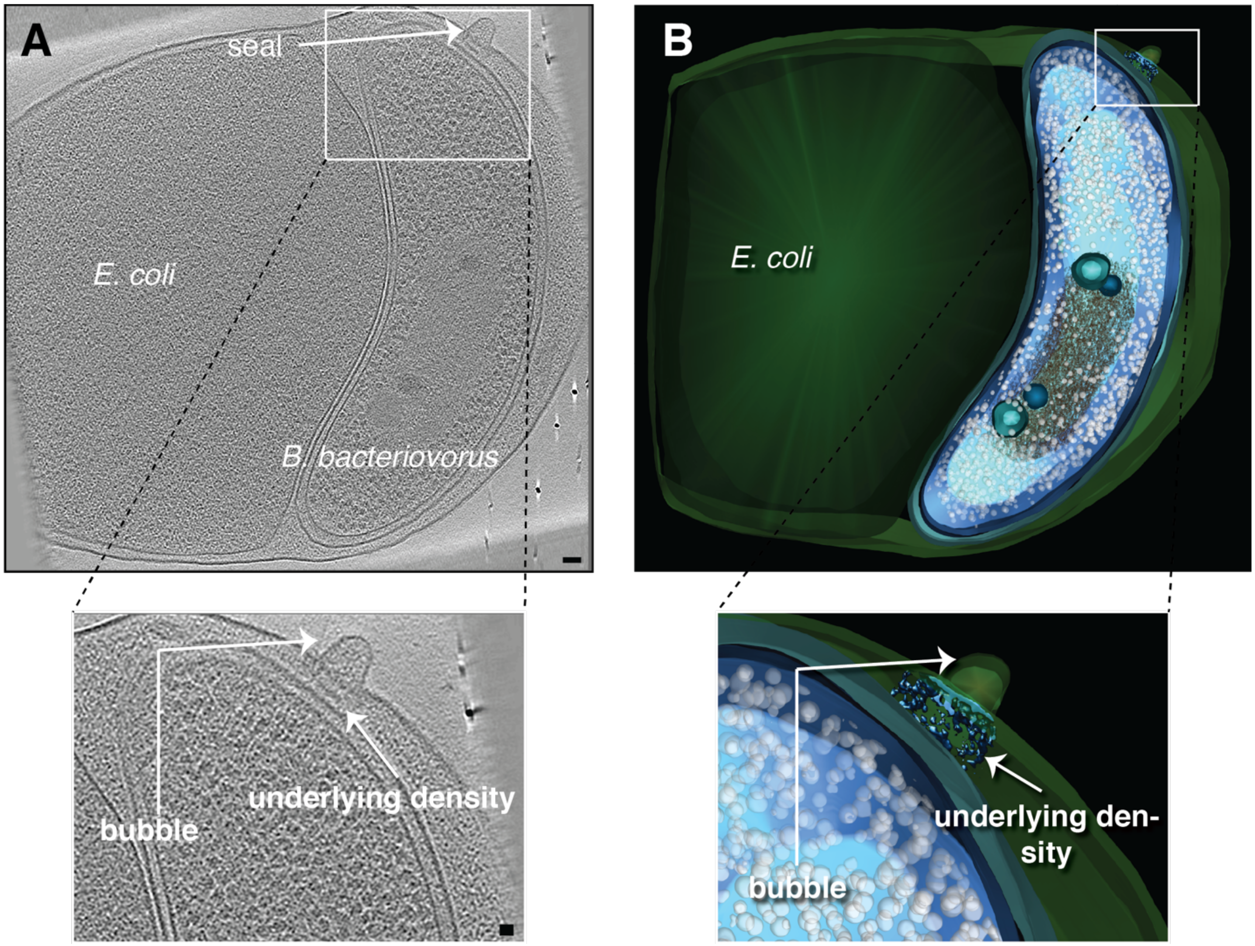


**Figure S28: A)** A slice through an electron cryo-tomogram (top) and enlargement (bottom) of an *E. coli* bdelloplast containing a *B. bacteriovorus* cell highlighting the bdelloplast seal. **B)** A 3D segmentation of the cryo-tomogram shown in (A) and an enlarged view of the seal. Scale bars 50 nm in the top panel of (A), and 20 nm in the enlargement.

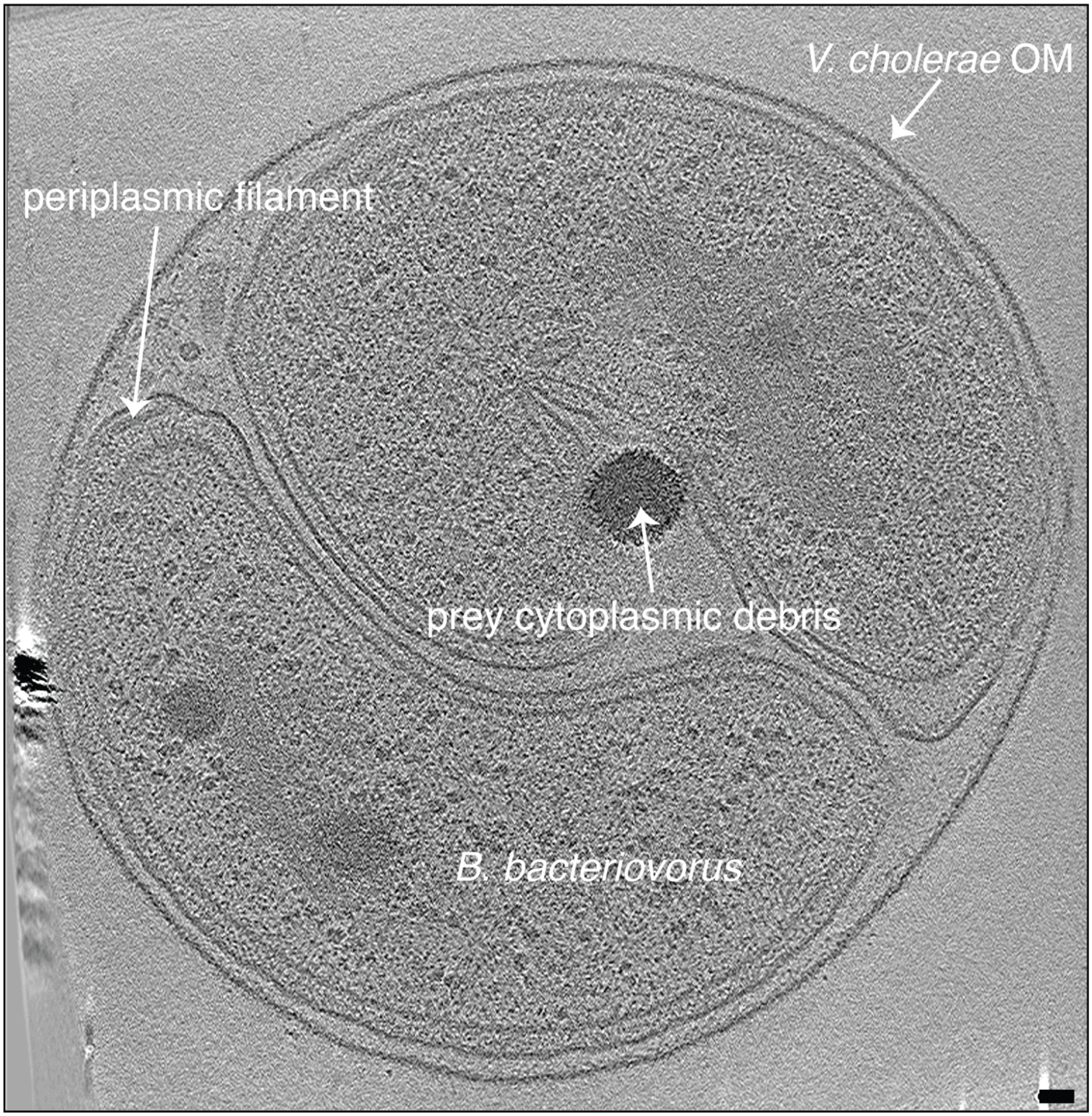


**Figure S29:** A slice through an electron cryo-tomogram of a *V. cholerae* bdelloplast containing two *B. bacteriovorus* cells highlighting a filamentous structure in the polar periplasm of one of the predator cells. Scale bar is 50 nm.

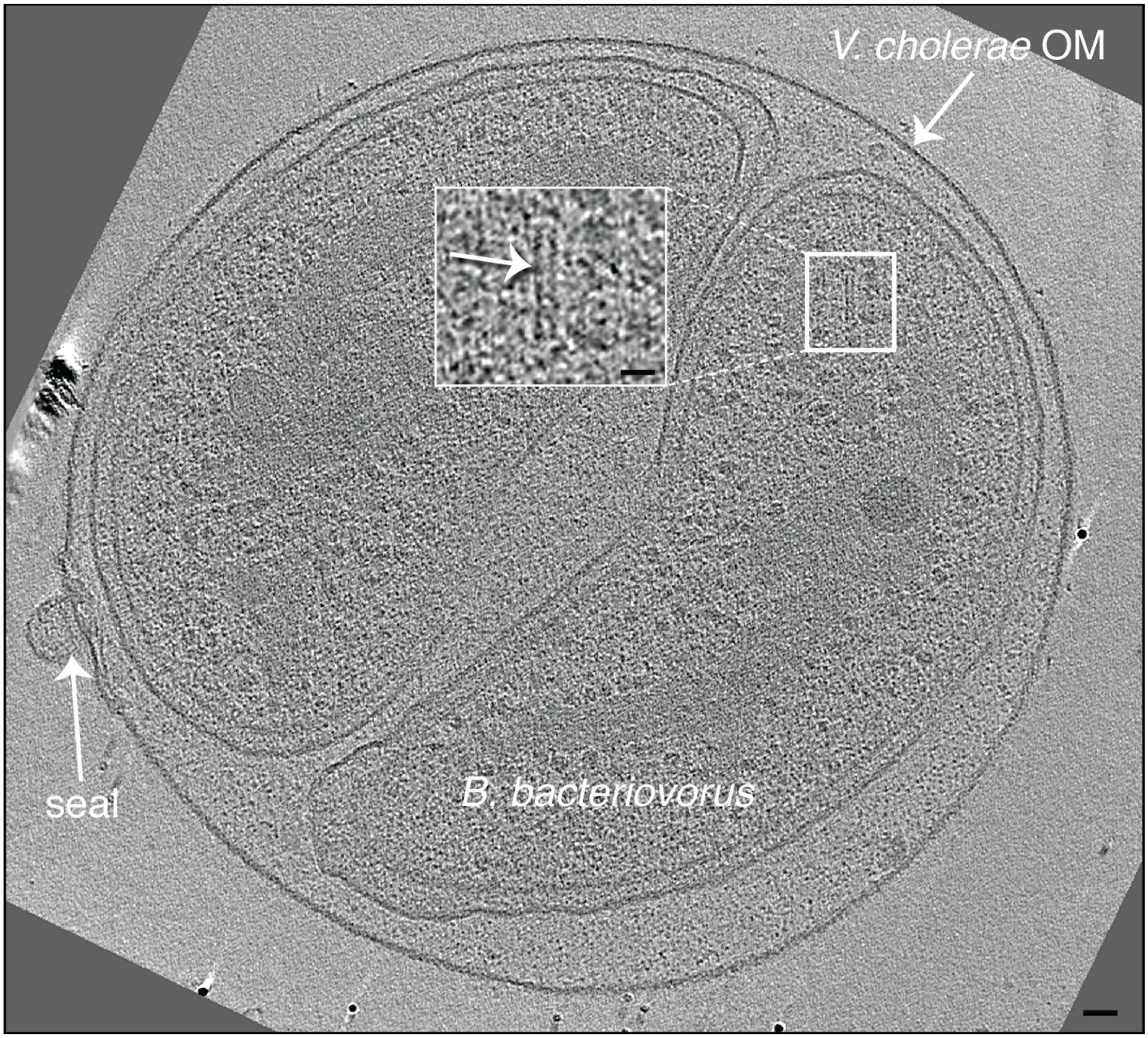


**Figure S30:** A slice through an electron cryo-tomogram and enlarged view (inset) of a *V. cholerae* bdelloplast containing two *B. bacteriovorus* cells highlighting an 8-nm wide tube in the cytoplasm of one of the predator cells (white arrow in the inset). Scale bars 50 nm in main panel, 20 nm in enlargement.

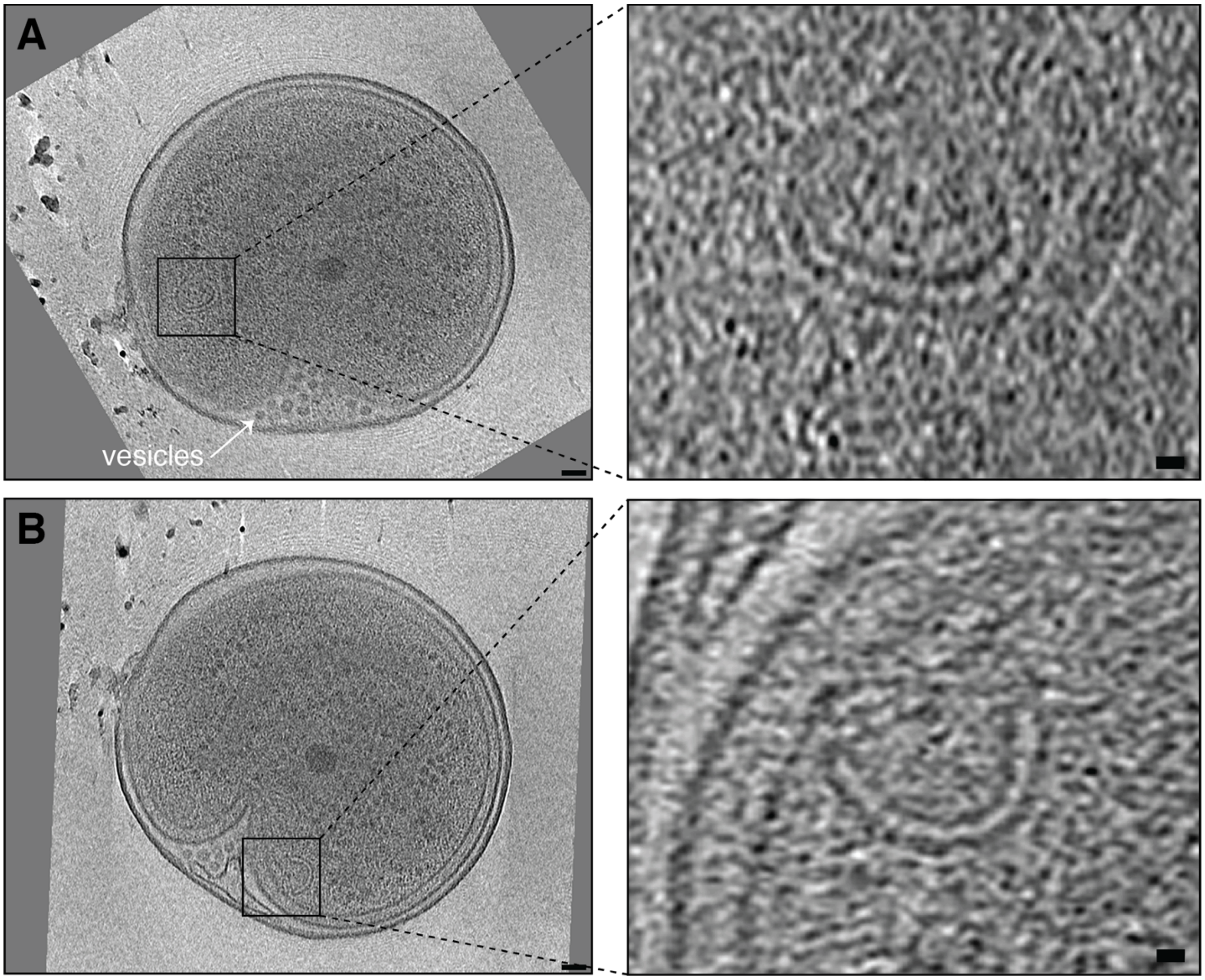


**Figure S31: A, B)** Slices at different z-heights through an electron cryo-tomogram (left) and enlarged views (right) of an end-stage *E. coli* minicell bdelloplast containing a *B. bacteriovorus* cell highlighting nested round- (A) and horseshoe-shaped (B) vesicles in the predator cytoplasm. Scale bars 50 nm in left panels and 20 nm in enlargements.

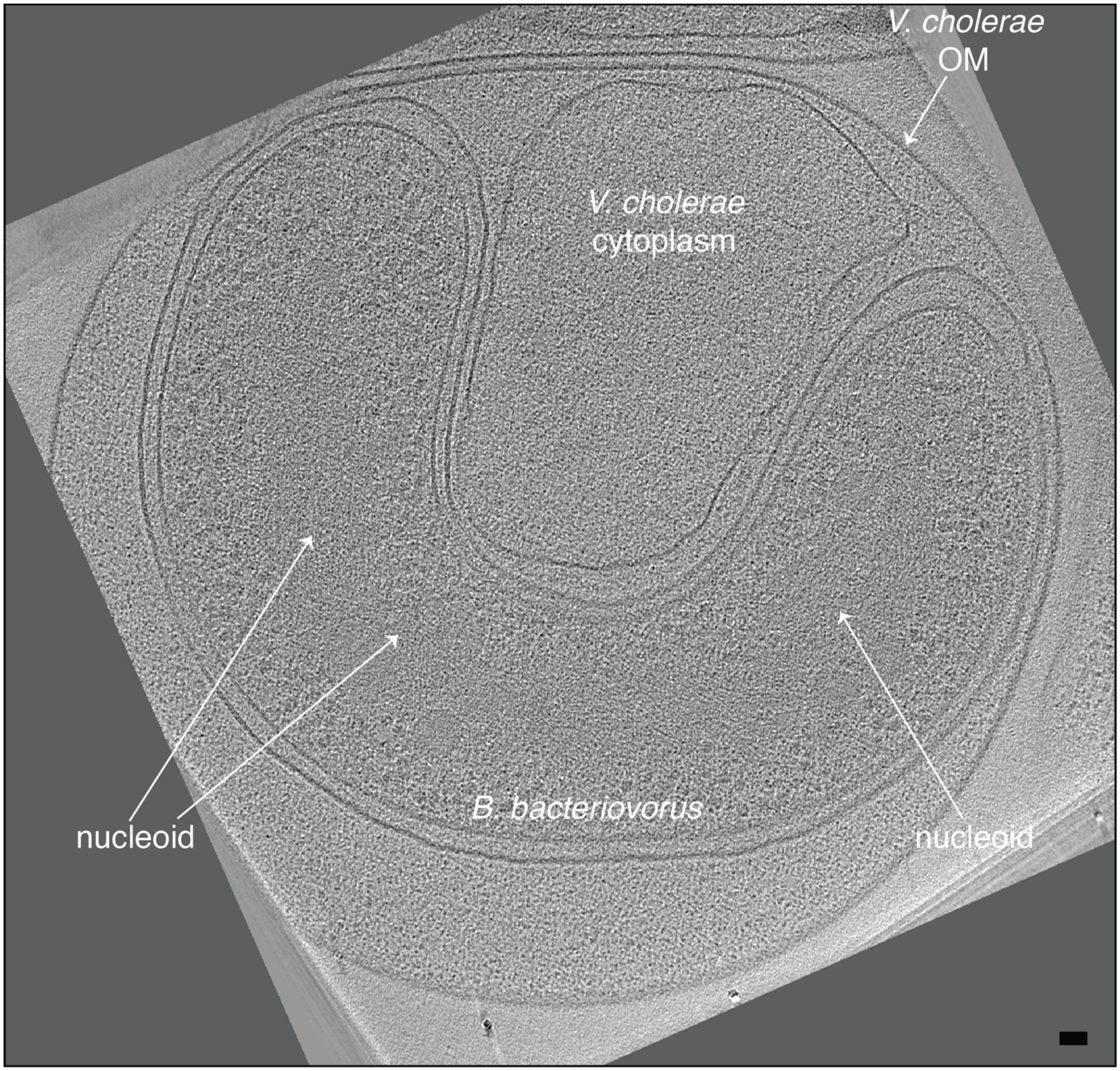


**Figure S32:** A slice through an electron cryo-tomogram of a *V. cholerae* bdelloplast showing an elongated *B. bacteriovorus* cell with a less densely compacted nucleoid than attack-phase cells. Scale bar is 50 nm.

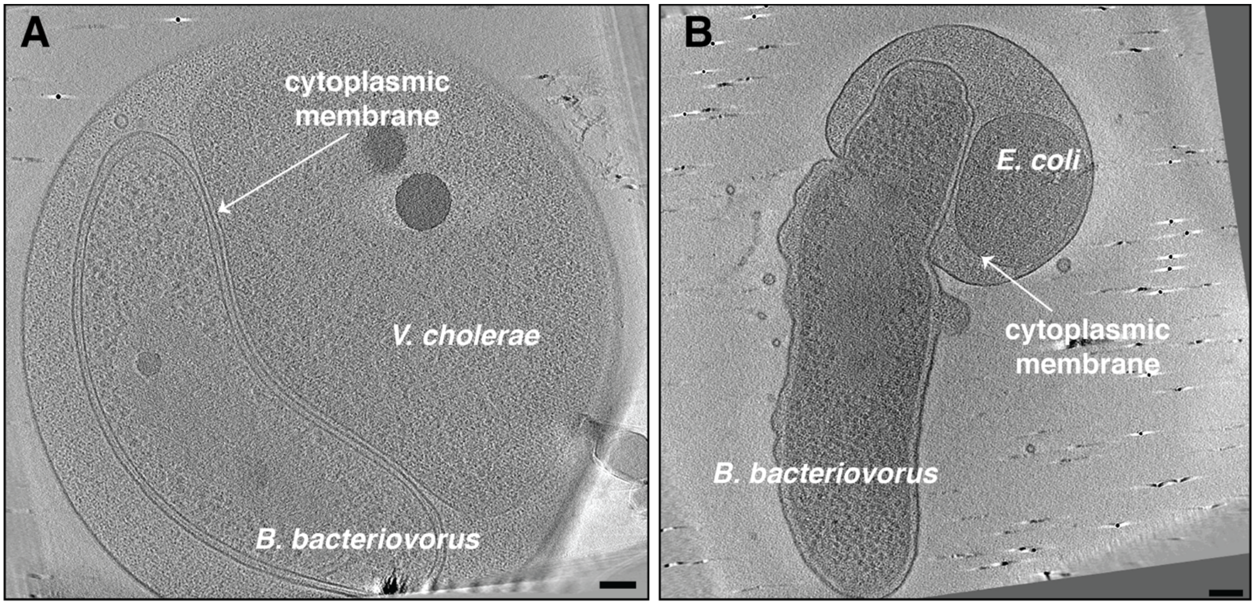


**Figure S33:** Slices through electron cryo-tomograms of a *V. cholerae* bdelloplast containing a *B. bacteriovorus* cell **(A)** and a stalled invasion of an *E. coli* minicell **(B)**, highlighting the proximity of the prey cytoplasmic membrane and the predator outer membrane. Scale bars 100 nm.

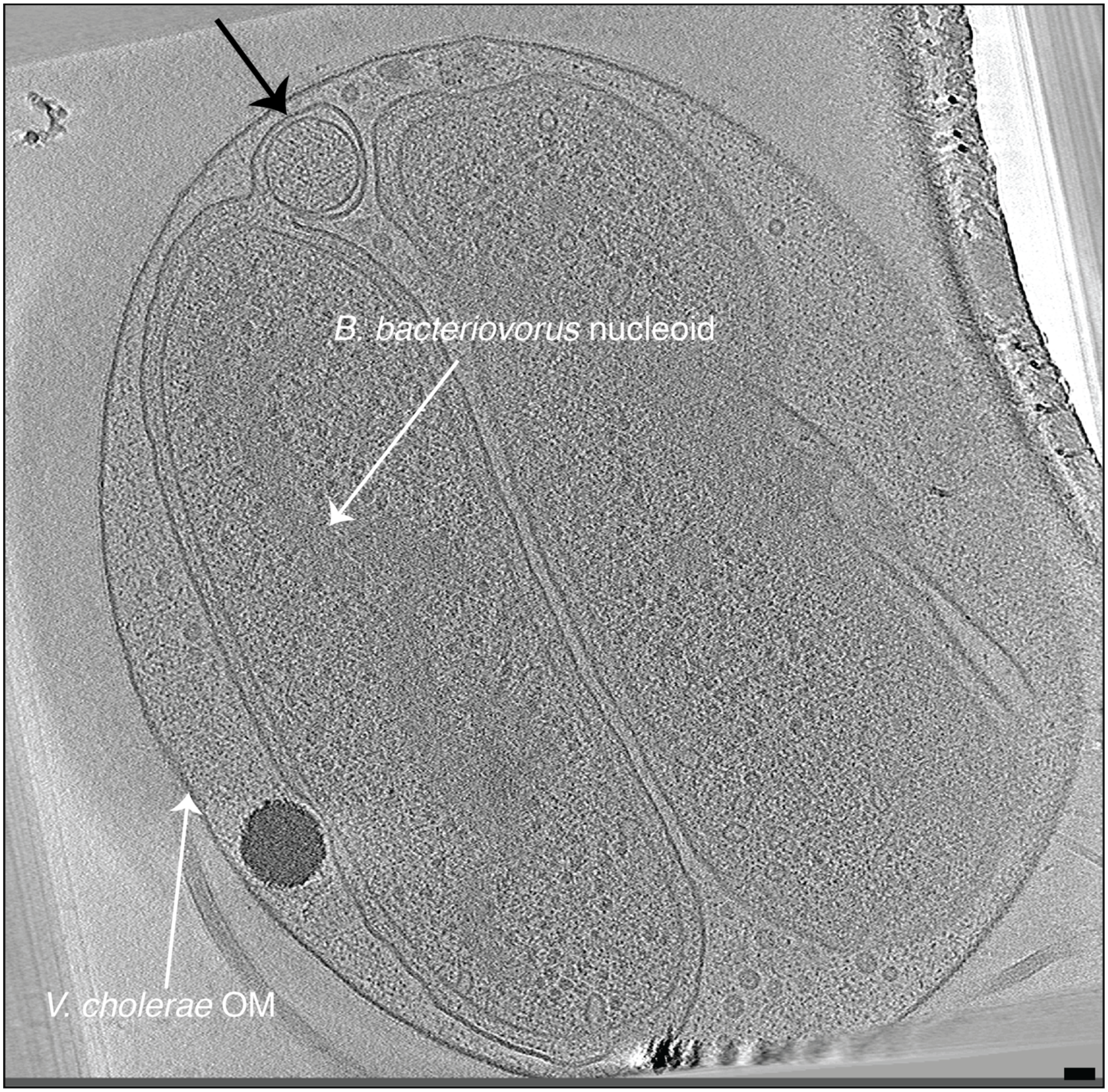


**Figure S34:** A slice through an electron cryo-tomogram of a *V. cholerae* bdelloplast showing a small spherical product (black arrow) resulting from division of the predator. Scale bar is 50 nm.

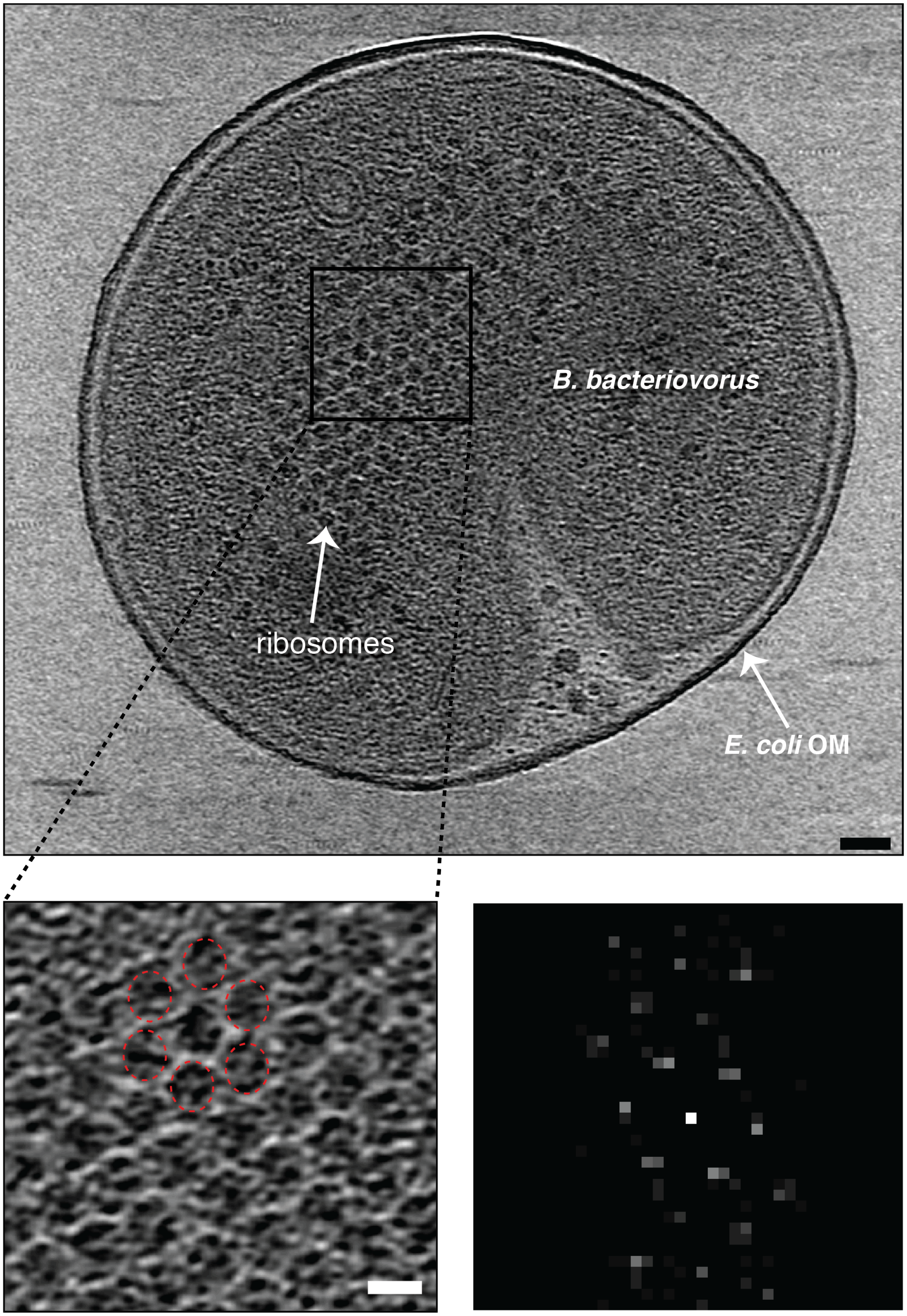
B. bacteriovorus
ribosomes
E. coli OM

**Figure S35: A)** A slice through an electron cryo-tomogram of an end-stage *E. coli* minicell bdelloplast showing a regular hexagonal lattice of ribosomes on the surface of the nucleoid. The enlargement at the bottom left highlights six ribosomes in the pattern. The power spectrum of the enlarged region is shown at the bottom right. Scale bars 50 nm in the top panel and 20 nm in the enlargement.

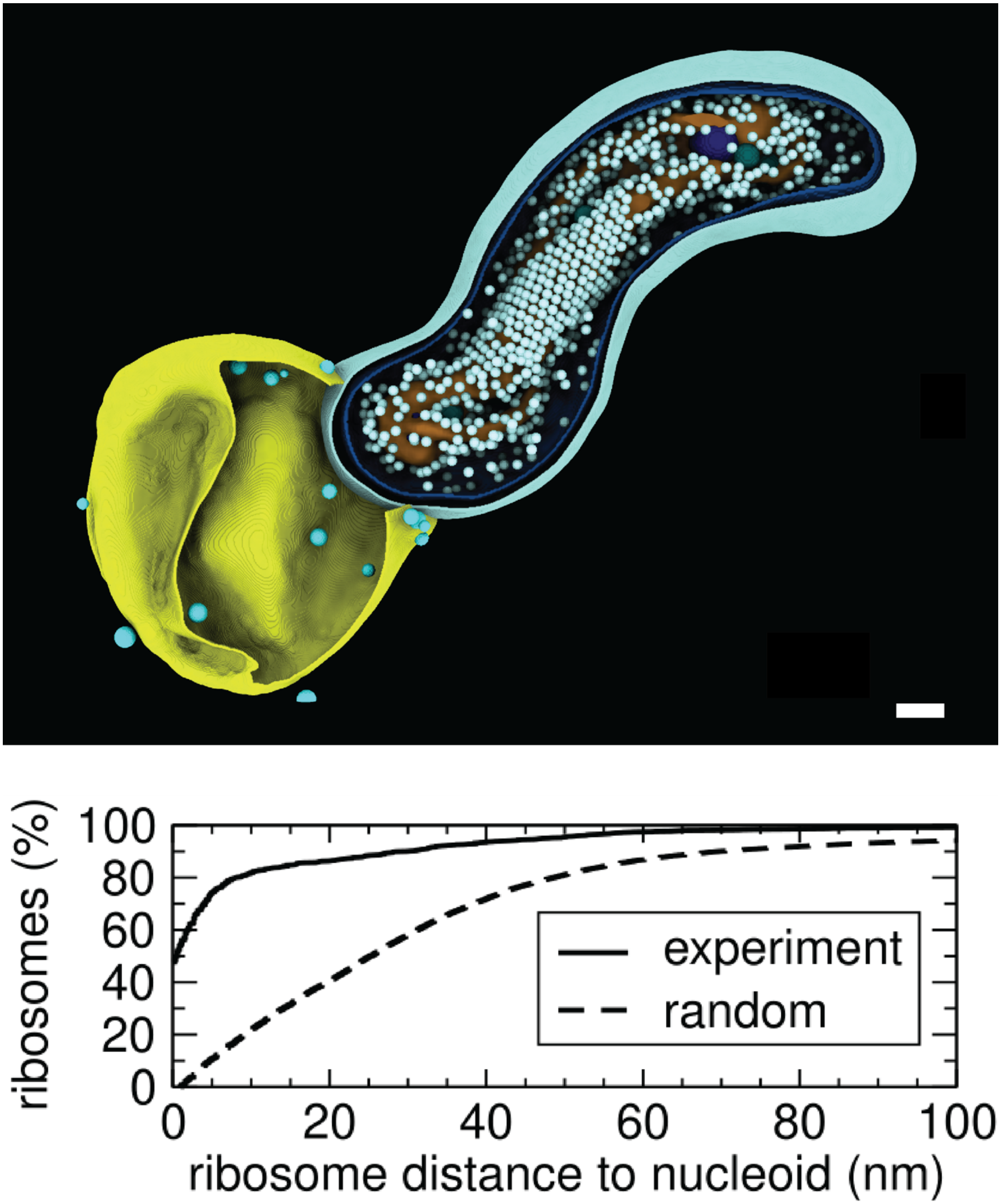


**Figure S36:** Distances of individual ribosomes from the nucleoid surface measured in the 3D segmentation of the shown cryo-tomogram of *B. bacteriovorus* ("experiment," solid line), compared to a simulation of randomly distributed 20 nm-wide spheres packed in the same segmented volume (“random,” dashed line). Scale bar 100 nm. The cryo-tomogram used in this analysis is shown in Movie S17.

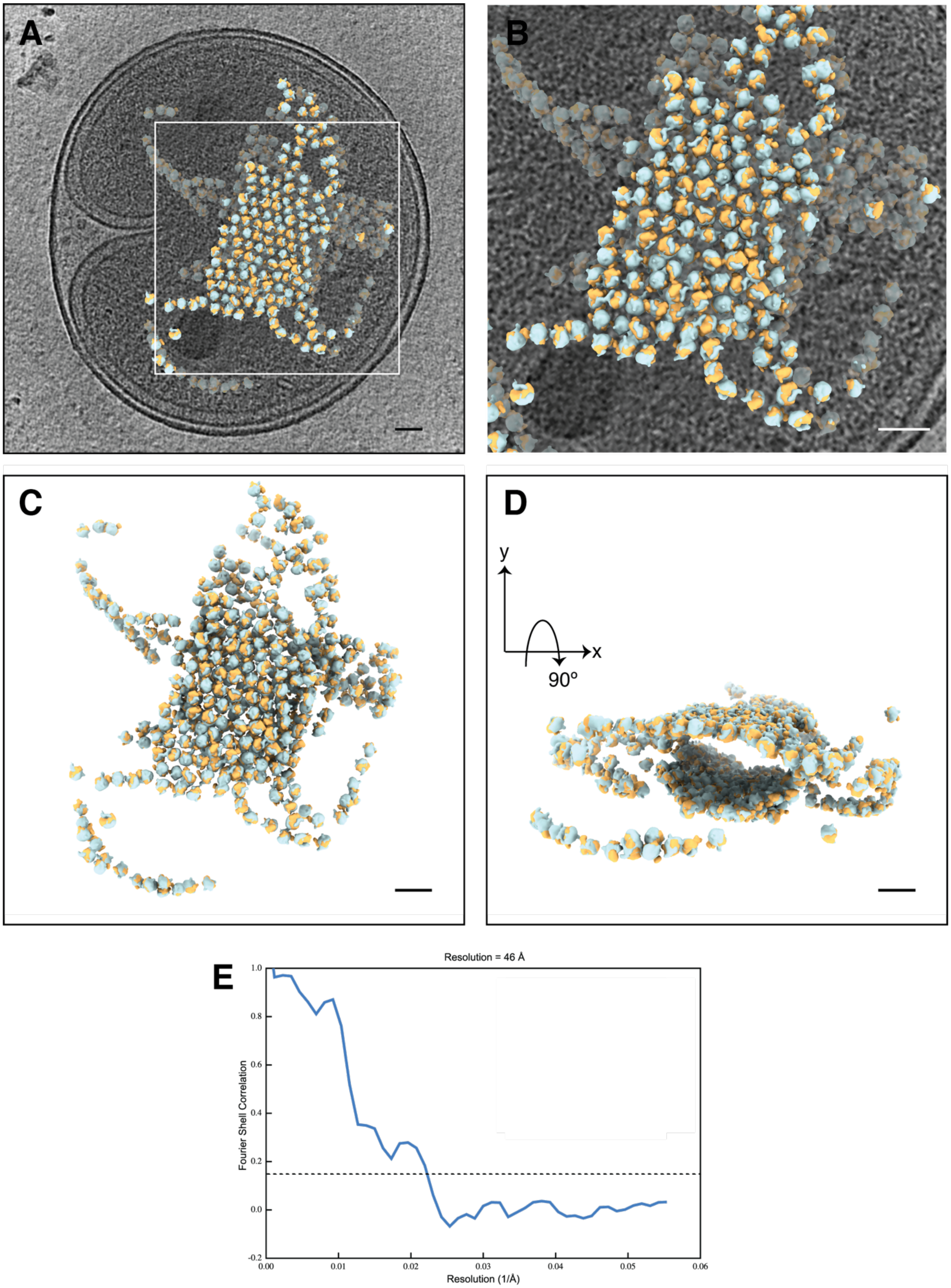
A
B
C
D
y
x
90°
E
Resolution = 46 Å
Fourier Shell Correlation
Resolution (1/Å)
1.0
0.8
0.6
0.4
0.2
0.0
-0.2
0.00
0.01
0.02
0.03
0.04
0.05
0.06

**Figure S37: A)** A slice through an electron cryo-tomogram (left) and enlarged view of the white-boxed area (right) of an end-stage bdelloplast (shown in Movie S15) with a superimposed view of a subtomogram average of regularly-arranged ribosomes on the nucleoid surface mapped back to their original positions and orientations. Cyan is the large ribosomal subunit; light yellow is the small subunit. **C, D)** The same regularly-arranged ribosomes shown in (A and B) rotated at different angles. **E)** Fourier shell correlation of the subtomogram average used in (A) obtained by averaging 775 ribosomes from the regular lattice. Scale bar is 50.

### Movie legends:

**Movie S1:** An electron cryo-tomogram of an attack-phase *B. bacteriovorus* cell highlighting limited arrangement of ribosomes on some parts of the surface of the compacted nucleoid. Note the flagellum and chemosensory array at one pole and T4aP at the opposite pole. Scale bar is 100 nm.

**Movie S2:** An electron cryo-tomogram of a *B. bacteriovorus* cell attached to an *E. coli* minicell and accompanying 3D segmentation. An attachment plaque and T4aP basal bodies (blue cylinders) can be seen at the biting pole, while an early stage of flagellar resorption can be seen at the other pole. Scale bar is 100 nm.

**Movie S3:** An electron cryo-tomogram of a *B. bacteriovorus* cell attached to an *E. coli* minicell and accompanying 3D segmentation. An attachment plaque, rose-like complexes (light green), and T4aP basal bodies (blue cylinders) can be seen at the biting pole, while an intermediate stage of flagellar absorption into the periplasm can be seen at the other pole. Scale bar is 100 nm.

**Movie S4:** An electron cryo-tomogram of a *B. bacteriovorus* cell attached to an *E. coli* minicell and accompanying segmentation. An attachment plaque and T4aP basal bodies (blue cylinders) can be seen at the biting pole, while a late stage of flagellar resorption can be seen at the other pole. The broken periplasmic flagellar filament is wrapped around the cell. Scale bar is 100 nm.

**Movie S5:** An electron cryo-tomogram of a *B. bacteriovorus* cell attached to an *E. coli* minicell by an attachment plaque. A late stage of flagellar resorption can be seen at the other pole with the periplasmic flagellar filament wrapping around the cell. Scale bar is 50 nm.

**Movie S6:** An electron cryo-tomogram of a *B. bacteriovorus* cell attached to an *E. coli* minicell with an attachment plaque. A late stage of flagellar resorption can be seen where the exit hole of the flagellum is far from the motor and the periplasmic flagellar filament wraps around the cell. Scale bar is 50 nm.

**Movie S7:** An electron cryo-tomogram of two *B. bacteriovorus* cells, one of them attached to an *E. coli* minicell with an attachment plaque. A late stage of flagellar resorption can be seen with the periplasmic flagellar filament wrapping around the cell. No flagellar motor could be identified in this cryo-tomogram. Scale bar is 50 nm.

**Movie S8:** An electron cryo-tomogram of a stalled invasion by a *B. bacteriovorus* cell of an *E. coli* minicell. A portal can be seen surrounding the entry hole and many vesicles are present inside the prey. Scale bar is 100 nm.

**Movie S9:** An electron cryo-tomogram of an end-stage stalled invasion by a *B. bacteriovorus* cell of an *E. coli* minicell, in which all the prey cytoplasm has been consumed. A portal with associated membrane blebs can be seen at the entry hole. The ribosomes exhibit a regular hexagonal packing around much of the nucleoid surface. Scale bar is 100 nm.

**Movie S10:** An electron cryo-tomogram of an *E. coli* bdelloplast, and accompanying 3D segmentation, highlighting the seal at the entry hole. A significant part of the prey's cytoplasm is still present. Scale bar is 100 nm.

**Movie S11:** An electron cryo-tomogram of a *V. cholerae* bdelloplast highlighting multiple uniformly-sized vesicles inside the bdelloplast. A significant part of the prey's cytoplasm is still present. Scale bar is 50 nm.

**Movie S12:** An electron cryo-tomogram of a *V. cholerae* bdelloplast containing two newly-divided *B. bacteriovorus* cells highlighting the bdelloplast seal and prey flagellar relic. Inside the bdelloplast, uniformly-sized vesicles and a dense sphere of yet-undigested prey cytoplasm are present. Scale bar is 100 nm.

**Movie S13:** An electron cryo-tomogram of an end-stage *E. coli* minicell bdelloplast highlighting the seal, multiple uniformly-sized vesicles and ribosomes in a regular hexagonal arrangement on the nucleoid surface. Scale bar is 100 nm.

**Movie S14:** An electron cryo-tomogram of an end-stage *E. coli* minicell bdelloplast highlighting multiple uniformly-sized vesicles and a hexagonal lattice of ribosomes around the nucleoid. No prey cytoplasm remains. Scale bar is 100 nm.

**Movie S15:** An electron cryo-tomogram of an end-stage *E. coli* minicell bdelloplast highlighting multiple uniformly-sized vesicles and a hexagonal lattice of ribosomes around the nucleoid. No prey cytoplasm remains. Scale bar is 100 nm.

**Movie S16:** An electron cryo-tomogram of an end-stage *E. coli* minicell bdelloplast containing two newly-divided *B. bacteriovorus* cells and accompanying segmentation. No prey cytoplasm remains and many ribosomes can be seen regularly arranged around the nucleoids of the two predator cells. Scale bar is 100 nm.

**Movie S17:** An electron cryo-tomogram of a *B. bacteriovorus* cell associated with a lysed *E. coli* minicell and accompanying segmentation, highlighting the regular arrangement of many ribosomes around the nucleoid. Scale bar is 100 nm.

**Movie S18:** A summary of the invasion cycle of *B. bacteriovorus* based on the *in situ* cryo-ET data presented in this study.

**Table S1:** Number of tomograms collected of various stages of the *B. bacteriovorus* lifecycle.

| Stage | No. of Tomograms |
|---|---|
| Attack phase | 171 |
| Predator close or attached to a prey | 166 |
| Stalled invasion of *E. coli* minicells | 18 |
| Early to mid-stage bdelloplasts (some prey cytoplasm still present) | 29 |
| End-stage bdelloplasts (all prey cytoplasm consumed) | 25 |

**Table S2:** Number of examples observed of various features described in this study.

| Feature | No. of observations |
|---|---|
| Flagellar motor | 1 per attack phase cell |
| Cytoplasmic tubes (8 nm in diameter) | 2 on average per cell |
| Fimbriae | Multiple per attack-phase cell |
| Piliated T4aP | 32 |
| Non-piliated T4aP | 3-6 on average per attack-phase cell |
| Rose-like complex | 2-3 on average per attack-phase cell |
| Unidentified periplasmic tubular structures | 6 |
| Extracellular vesicles near attack-phase cells | 1-10 per cell |
| External knob-like structures | 33 on 2 cells near lysed prey |
| Predator attached to a prey with piliated T4aP | 6 |
| Attachment plaque | 57 |
| Absorbed flagellum (early) | 8 |
| Absorbed flagellum (intermediate) | 4 |
| Absorbed flagellum (late) | 30 |
| Invasion portal | 18 (all in stalled invasions of *E. coli* minicells) |
| Bdelloplast seal | 1 per bdelloplast |
| Prey flagellum relic on bdelloplast | 12 |
| Vesicles in prey | 72 (all in bdelloplasts/stalled invasions) |
| Possible T4aP basal bodies lacking the extracellular ring inside bdelloplast | 25 |
| Small spherical cell division products in bdelloplast | 3 |
| Ribosome lattice on nucleoid | 33 (all in end-stage bdelloplasts/stalled invasions) |

**Materials and Methods:**

**Strains and growth conditions:**

*B. bacteriovorus* HD100 cells were grown and prepared as described in (Lambert and Sockett, 2008). To enrich for attack phase, *B. bacteriovorus* cells were grown in S17-1 prey solution at 30°C, then filtered by a 0.45 µm filter. To enrich for attachment, *B. bacteriovorus* cells were grown in S17-1 prey solution at 30° C, then filtered by a 0.45 µm filter and concentrated by centrifugation. *E. coli* WM3433 (minicell-producing) (Liu et al., 2011) were cultured in LB medium at 30° C. Minicells were enriched by 2-step centrifugation (initially 3,000 x g for 3 minutes and subsequently 9,000 x g for 5 minutes). The two species were then mixed for 15 minutes. To enrich for stalled invasions, the two species were mixed for 30 minutes. To enrich for bdelloplasts, *B. bacteriovorus* cells were grown in S17-1 prey solution at 30°C, then filtered by a 0.45 µm filter. *V. cholerae* strain MKW 1383 was grown overnight in LB medium. Subsequently, *B. bacteriovorus* and *V. cholerae* were mixed and incubated in Ca-HEPES buffer at 30°C for 16 hours.

To enrich for end-stage bdelloplasts, a co-culture of *B. bacteriovorus* HD100 and *E. coli* WM3433 was incubated at 30°C, 220 rpm for 48 h, then was used to inoculate 100 mL HEPES buffer with 4 mL of overnight *E. coli* prey culture and grown for another 48 h. Cells were harvested by centrifugation at 3,500 x g at 4°C for 20 min. The pellet was resuspended in 1 mL ice-cold HG-buffer (HEPES buffer + 10 mg/mL gelatin), loaded on a sucrose gradient (20% sucrose in HG buffer (40 mL), frozen at -20°C and thawed overnight), and centrifuged at 2,000 x g with no brake at 4°C for 30 min. The broad mid-band was collected and washed twice by centrifugation at 10,000 x g for 5 min before resuspension in HEPES buffer for vitrification.

**Cryo-ET sample preparation and imaging:**

Samples were mixed with a solution of BSA-treated 10-nm gold and 3-4 µL were applied to freshly glow-discharged R2/2 carbon-coated 200 mesh copper Quantifoil grids (Quantifoil Micro Tools). Samples were plunge-frozen in a liquid ethane/propane mixture in a Vitrobot Mark III or Mark IV (FEI). For all samples except the end-stage *E. coli* minicell bdelloplast enrichment, imaging was performed with an FEI Tecnai G2 Polara 300 keV field emission gun transmission electron microscope (FEI company, Hillsboro, OR, USA) equipped with a Gatan energy filter and K2 Summit direct electron detector (Gatan, Pleasanton, CA, USA) at Caltech or an FEI Titan Krios equipped with Gatan energy filter and Gatan K2 Summit direct detector at the Janelia Research Campus of HHMI. Tilt-series were collected from -60° to +60° in 1° increments using UCSF Tomography software (Zheng et al., 2007), with a cumulative electron dose of 100 $e^-$/ Å$^2$, a target defocus of -8 µm, and a pixel size of 3.3 Å, 3.9 Å, 4 Å or 4.2 Å.

End-stage bdelloplast-enriched samples were imaged using the fast-incremental single exposure (FISE) method (Chreifi et al., 2019; Eisenstein et al., 2019) with SerialEM software (Mastronarde, 2005) using a dose-symmetric tilt scheme adapted from (Hagen et al., 2017). Data was collected using a Titan Krios 300 keV field emission gun transmission electron microscope (Thermo Fisher Scientific) equipped with a Gatan imaging filter and a K2 Summit direct detector in counting mode (Gatan) at Caltech. The tilt range was -60° to +60° with 3° tilt increment, target defocus of -8 µm, pixel size of 4.52Å, frame rate of 0.05 sec/frame, and a total dose of 130 $e^-$/Å$^2$. FISE tilt-series were gain-normalized and motion-corrected using the alignframes function of IMOD (Kremer et al., 1996).

**Image processing and subtomogram averaging:**

Three-dimensional reconstructions of tilt-series were performed using either the IMOD software package (Kremer et al., 1996), or automatically with RAPTOR through the Jensen Lab processing pipeline (Ding et al., 2015). Subtomogram averaging was done using the PEET program (Nicastro, 2006) with 2-fold symmetrization applied along the particle Y-axis. The number of particles averaged were: 132 particles for the rose-like complex, 335 particles for the non-piliated T4aP basal body, 79 particles for the flagellar motor.

For the subtomogram average of the ribosomes in end-stage bdelloplasts, the selected tilt-series were aligned, contrast transfer function corrected, and reconstructed using EMAN2.91 (Chen et al., 2019; Tang et al., 2007). A total of 775 nucleoid-surrounding ribosomes from 2 tomograms were selected manually. The averaged structure of the ribosome was produced according to the "gold-standard" protocol using a subtomogram refinement pipeline in EMAN2 (Chen et al., 2019). During alignment, particles shifts were limited to ~30 Å and a loose spherical mask was applied. The final, mask-corrected Fourier shell correlation (FSC), was estimated in RELION3 using a soft-edge mask (Fig. S39E) (Zivanov et al., 2018). To analyze spatial relationships between nucleoid-coating ribosomes, the averaged map was placed back into particle locations in one of the original tomograms using EMAN2 (Chen et al., 2019). Ribosome maps were dusted, segmented, and visualized in their original orientations within the tomogram using ChimeraX (Pettersen et al., 2021a).

To classify non-piliated T4aP particles based on the presence/absence of the lower periplasmic ring, we used the built-in principal component analysis (PCA) and k-means clustering functions of the PEET package (Heumann et al., 2011). After an initial T4aP subtomogram average was obtained, a binary mask was drawn on the region of interest (the lower periplasmic ring) and used for PCA analysis. Subsequently, the particles were separated into two classes using 5 to 20 components and re-aligned based on the class number. The best clustering parameters were selected based on the Akaike information criterion (AIC) and Bayesian information criterion (BIC) values and by visually inspecting the realigned particles.

Average density profiles for the portal were automatically calculated with a custom script, sideview-profile-average, written by Davi Ortega (https://www.npmjs.com/package/sideview-profile-average) and described in (Ortega et al., 2020).

**Tomogram segmentations:**

Segmentations shown in Figures 4, 7, S15, S18, S19, S30 and Movies S2, S3, S4, S8, S10, S12 were produced manually using IMOD (Kremer et al., 1996). For Figures 8 and S38 and Movies S16 and S17, features were segmented automatically using the VISFD segmentation tools (Jewett, 2021a) and PoissonRecon (Calakli and Taubin, 2012; Kazhdan and Hoppe, 2013), and edited and visualized using ChimeraX (Pettersen et al., 2021b). A tutorial is available demonstrating how to use VISFD to segment tomograms of *B. bacteriovorus* cells (Jewett, 2021b). Note that manual corrections were often needed after the automatic segmentations.

To segment the nucleoid of *B. bacteriovorus* (Figures 8 and S38 and Movies S16 and S17), nucleoid density was inferred from the absence of other objects nearby such as ribosomes and proteins. For every voxel in the image, we calculated the fluctuations in brightness within a sphere of radius 20 nm. (See the VISFD documentation for the “-fluct” filter.) To detect the nucleoid, we searched for regions in the image with relatively uniform brightness (where the fluctuations in nearby brightness were comparatively low). However, this simple method can mistake other homogeneous regions in the cell for nucleoid, such as regions above and below the cytoplasm in the Z direction (The upper and lower boundary of the cell is not visible due to the "missing wedge" artifact.) These regions were removed manually using the volume editing tools included with ChimeraX (Pettersen et al., 2021a).

Ribosomes were detected using the method of scale-free blob detection (Lindeberg, 1998), to detect dark blobs in the image of size between 120-215 Å (similar to the size of a ribosome). Dark blobs whose centers were closer than 12 nm to other blobs were discarded. Unfortunately, ribosomes are not sufficiently dark to be reliably detected using this method, especially in crowded environments. Furthermore, other dark objects of similar size can be mistaken for ribosomes. Hence, ribosomes which were not detected in the original image automatically were detected manually in IMOD (Kremer et al., 1996). Approximately 1/3 of the ribosomes were detected manually. Subsequently, ribosomes were approximated as spherical objects.

*B. bacteriovorus* membranes and their interior volumes were segmented automatically using VISFD using the ridge detection and tensor voting method (Martinez-Sanchez et al., 2014) to locate the visible surface of the membranes, followed by screened Poisson surface reconstruction (Calakli and Taubin, 2012) to generate closed surfaces and their interior volumes. The details of how this was done are explained in the VISFD tutorial (Jewett, 2021b). Manual corrections were applied when required using the volume editing tools included with ChimeraX (Pettersen et al.,

2021a).

**Plotting the distances between the ribosomes and the nucleoid surface:**

We first measured the distance from the center of each ribosome to the nearest identified voxel in the nucleoid. Then we subtracted 10 nm from these distances, because each ribosome is approximately 20 nm wide, and plotted these numbers on the horizontal axis. The same distances were then measured using randomly generated spheres. To generate them, spheres of 20 nm in diameter were added to the cytoplasmic volume at random locations. If a newly added sphere overlapped with the nucleoid or another sphere, or if it did not lie within the inner membrane, then it was discarded and a new randomly located sphere was generated. This process was repeated until the number of random spheres matched the number of ribosomes in the cytoplasmic volume for each tomogram (1,109 ribosomes for Figure S36 and 2,304 for Figure 6).
